# Genomic regions associate with major axes of variation driven by gas exchange and leaf construction traits in cultivated sunflower (*Helianthus annuus* L.)

**DOI:** 10.1101/2022.04.01.486477

**Authors:** Ashley M. Earley, Andries A. Temme, Christopher R. Cotter, John M. Burke

## Abstract

Stomata and leaf veins play an essential role in transpiration and the movement of water throughout leaves. These traits are thus thought to play a key role in the adaptation of plants to drought and a better understanding of the genetic basis of their variation and coordination could inform efforts to improve drought tolerance. Here, we explore patterns of variation and covariation in leaf anatomical traits and analyze their genetic architecture via genome-wide association (GWA) analyses in cultivated sunflower (*Helianthus annuus* L.). Traits related to stomatal density and morphology as well as lower order veins were manually measured from digital images while the density of minor veins was estimated using a novel deep learning approach. Leaf, stomatal, and vein traits exhibited numerous significant correlations that generally followed expectations based on functional relationships. Correlated suites of traits could further be separated along three major principal component (PC) axes that were heavily influenced by variation in traits related to gas exchange, leaf hydraulics, and leaf construction. While there was limited evidence of colocalization when individual traits were subjected to GWA analyses, major multivariate PC axes that were most strongly influenced by several traits related to gas exchange or leaf construction did exhibit significant genomic associations. These results provide insight into the genetic basis of leaf trait covariation and showcase potential targets for future efforts aimed at modifying leaf anatomical traits in sunflower.

**Significance Statement:** Using traditional and automated/high-throughput (using a novel deep learning approach) phenotyping methods we studied leaf anatomical variation in sunflower. Genome-wide association (GWA) analyses identified numerous genomic regions underlying individual trait variation and regions underlying major multivariate axes of phenotypic variation. These results illustrate the value of employing a multivariate approach to GWA analyses and shed light on the extent to which leaf trait (co-)variation can be genetically decoupled to explore novel phenotypic space.

## Introduction

Stomata and leaf veins are central players in plant water relations. Veins distribute water throughout the leaf and stomata control the rate of transpiration (Sack and Scoffoni, 2013). As such, these traits are generally thought to play a key role in the adaptation of plants to drought (e.g., Bertolino *et al*., 2019; Sack and Scoffoni, 2013), a major agricultural stress that limits plant growth and productivity worldwide (NOAA; IPCC, 2014). Plant performance in water-limited conditions is known to be influenced by stomatal and vein traits (Lei *et al*., 2018; Scoffoni *et al*., 2011; Buckley, 2019). Stomatal conductance (*g_s_*), a measure of the conductance of CO2 and water vapour through the stomata, is determined by the physiological control of stomatal opening and closing as well as the size and distribution of stomata (Faralli *et al*., 2019). Similarly, vein length per area (VLA) affects leaf hydraulic conductance (*K_leaf_*), or the ratio of water flow rate to the water potential gradient across the leaf, and thus greatly affects how long stomata can remain open without drying out the leaf (Sack and Scoffoni, 2012). While substantial work has been done on the developmental and regulatory pathways determining stomatal density and morphology (e.g., Gudesblat *et al*., 2012; Bergmann and Sack, 2007) less is known about the genetic basis of the observed relationships between stomatal and leaf vein traits.

Leaf veins distribute water from the petiole throughout the leaf. In the intracellular space, water then moves through the mesophyll to the ultimate site of transpiration, the stomata. It is important to note here that stomata can be distributed on both the top (adaxial) and bottom (abaxial) sides of the leaf, and that species with stomata on both sides (i.e., amphistomatous) tend to have greater gas exchange capacity than species with stomata on a single side (Xiong and Flexas, 2020). An increase in the number of stomata on the upper side leads to increased maximum photosynthetic rates and to increased rates of transpiration due to increased CO2 diffusion (Muir, 2018; Xiong and Flexas, 2020). Due to these effects on water supply and loss, a close relationship between vein density and stomatal size/density is expected to be important for the optimization of water transport and transpiration (Fiorin *et al*., 2016; Bertolino *et al*., 2019). In particular, a shorter vein-to-stomata distance is thought to improve gas exchange and photosynthetic performance (Brodribb *et al*., 2007; de Boer *et al*., 2012). This distance is affected by traits such as stomatal density, vein density (estimated as vein length per area; VLA), leaf thickness, and cell shape (Brodribb *et al*., 2007; de Boer *et al*., 2012). As a general rule, vein and stomatal density are positively correlated (Sack and Scoffoni, 2013) while stomatal size and density are negatively correlated (Shahinnia *et al*., 2016; Doheny-Adams *et al*., 2012). Smaller epidermal cell size is also known to correlate with increases in stomatal and vein densities (Simonin and Roddy 2018; Brodribb *et al*. 2013) Such correlations are observed within (Carins Murphy *et al*., 2014) and across species (Zhang *et al*., 2012), though the extent to which these trait correlations are conditioned by genetic correlations (i.e., linkage or pleiotropy of major effect loci) remains an open question.

At the whole leaf level, plant species exhibit a range of strategies related to the cost of leaf construction. Traits such as leaf mass per area (LMA) and VLA play a role in leaf construction in addition to affecting leaf hydraulics (Xing *et al*., 2021; Poorter *et al*., 2009; John *et al*., 2017). These strategies occur along a major axis of leaf trait variation (the Leaf Economics Spectrum, LES), which ranges from resource conservative (i.e., ‘slow’) with a greater investment in leaf construction to resource acquisitive (i.e., ‘fast’) with a smaller investment in leaf construction (Wright *et al*., 2004; Reich, 2014; Díaz *et al*., 2016). A key indicator trait for the location of species along this spectrum is LMA, with leaf hydraulic traits (allocation to major vs. minor veins in particular) accounting for a portion of the observed variation in LMA (John *et al*., 2017). Investigating the variation present in both finer scale aspects of leaves as well as whole leaf traits promises to broaden our understanding of the evolution of correlated suites of leaf traits.

To date, most studies focusing on within species variation in leaf anatomy have either relied on relatively small sample sizes or have largely focused on stomatal traits to the exclusion of vein traits (e.g., Khan *et al*., 2003; Ries *et al*., 2012; Shi *et al*., 2021; Haworth *et al*., 2021). The general dearth of studies that explicitly examine the genetic basis of stomatal and vein traits, particularly in tandem, is likely due to challenges associated with the large-scale phenotyping of such traits. One study on wild tomato found that strongly correlated leaf traits were not controlled by the same QTL, suggesting that natural selection had favored particular trait combinations (Muir *et al*., 2014). However, another study on wild *Arabidopsis* accessions found that stomatal density correlated with various other leaf traits, including stomatal index and pavement cell density, and that they seemed to share a common genetic basis (Delgado *et al*., 2011). The lack of a clear pattern across species leaves as an open question the extent to which variation in such traits can be decoupled, thereby allowing them to vary independently.

Here, we describe patterns of phenotypic variation in leaf traits in cultivated sunflower (*Helianthus annuus* L.) and investigate their genetic architecture via genome-wide association (GWA) analyses. Domesticated from the common sunflower (also H. annuus; Wills and Burke, 2006; Blackman *et al*., 2011), cultivated sunflower is one of the world’s most important oilseed crops (FAO, 2018). Often grown in rainfed regions, sunflower productivity is frequently dependent on natural patterns of precipitation. While sunflower is generally regarded as being drought resistant due to its ability to root deeply (Connor and Sadras, 1992), drought is considered to be a major yield-limiting factor across the range of production (Hussain *et al*., 2018), making traits underlying plant-water relations a vital avenue of research. Prior work on sunflower has shown substantial variability and plasticity in leaf anatomical traits (Wang *et al*., 2020). Previous genomic analyses however, have been largely limited to higher level traits such as leaf mass, leaf area, LMA, and leaf mass fraction (Temme *et al*., 2020; Masalia *et al*., 2018). To overcome limitations in the analysis of vein traits, we developed a neural network deep learning approach to increase phenotyping efficiency from digital images (see Xu *et al*., 2020 for a similar approach), enabling an investigation of the genomic basis on finer scale leaf anatomical traits.

In this study, we sought to: (1) quantify phenotypic variation in stomatal and leaf vein traits and test for trait correlations in sunflower, (2) identify genomic regions underlying these traits using genome-wide association analyses, and (3) determine the extent to which observed trait correlations are due to a shared genetic basis. Our results provide insight into the genetic complexity of these traits and the degree to which observed trait correlations are constrained by linkage or pleiotropy of major effect loci and serve as a valuable first step toward optimizing leaf trait combinations via breeding.

## Results

We sampled leaves from a diversity panel of 239 cultivated sunflower genotypes from the Sunflower Association Mapping (SAM) population (Mandel *et al*. 2011) grown under greenhouse conditions. These leaves were then used to collect measurements for a variety of leaf anatomical traits (Figure 1). Traits of interest included stomatal density and size and vein length per area (VLA) for major and minor veins (Figure 1) as well as traits including leaf mass per area (LMA), midrib density, and plant biomass. With the exception of minor veins, which were traced using a novel deep learning neural network (Methods S1), all traits were measured manually. Trait correlations were examined using bivariate and multivariate analyses and genetic associations were determined using a custom GWA pipeline. See Experimental Procedures for full details.

**Figure 1:**
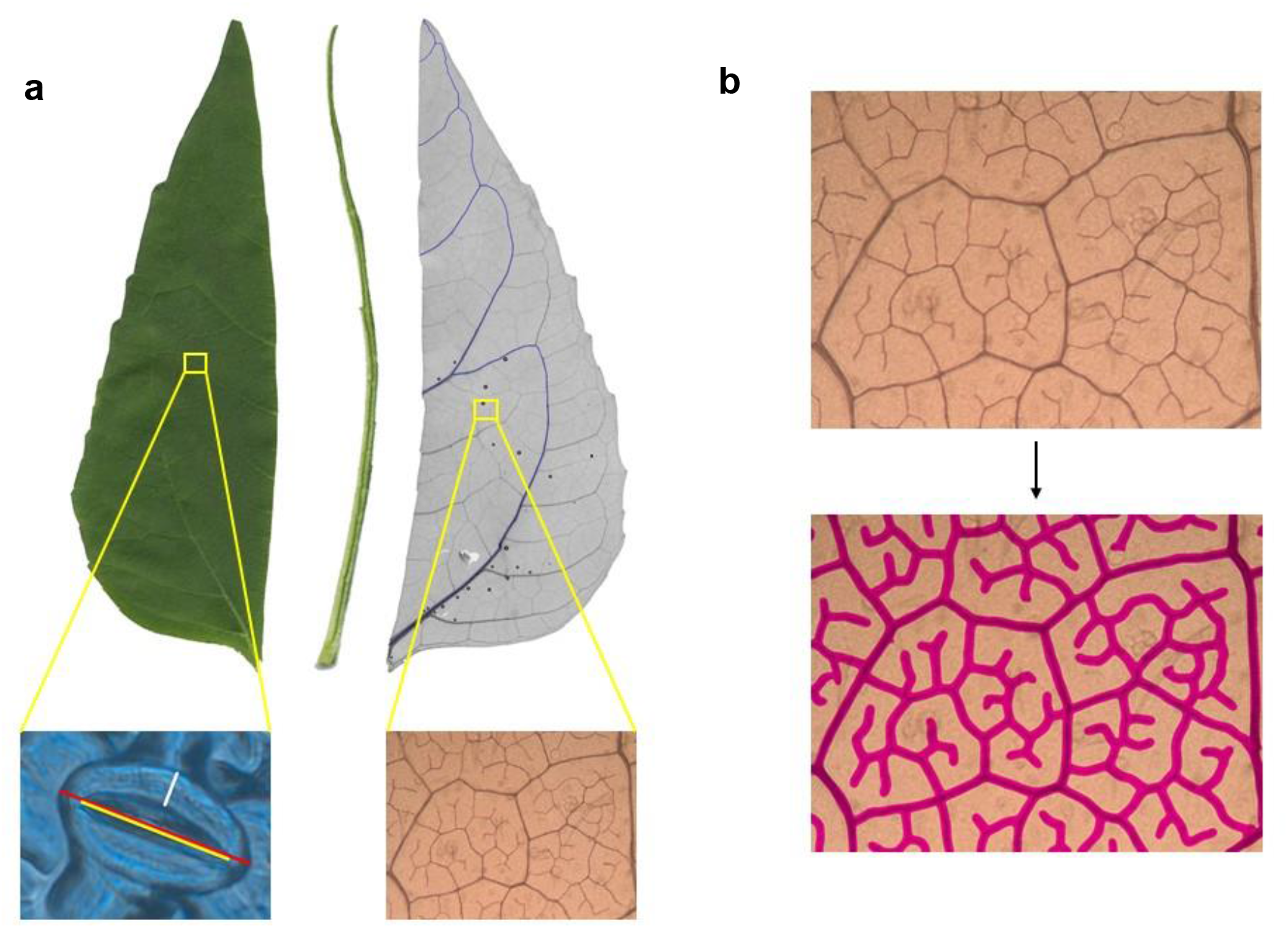
(a) Left: Image of dissected leaf. Right: Cleared and stained leaf showing major veins (green). Left Inset: A single stoma taken using the 100X objective. Colored lines indicate measurements taken: stomate length (red), pore length (yellow), and guard cell width (white). Right Inset: A microscope image of minor veins taken using the 5X objective. (b) Top: Image of minor veins taken at 5X. Bottom: Image of computer-traced minor veins using our deep learning approach. See Experimental Procedures for details.

### Patterns of phenotypic variation and trait correlations

Significant genotypic effects were detected for all traits measured except midrib mass fraction (Table 1). Table 1 lists the median, minimum, and maximum values found for all traits measured, demonstrating substantial trait variation across the population. Comparing manually measured VLA to the results from the neural network analysis revealed a strong correlation (Pearson’s r = 0.97), demonstrating the accuracy and validity of these computer-generated measurements (Figure S2).

**Table 1:**
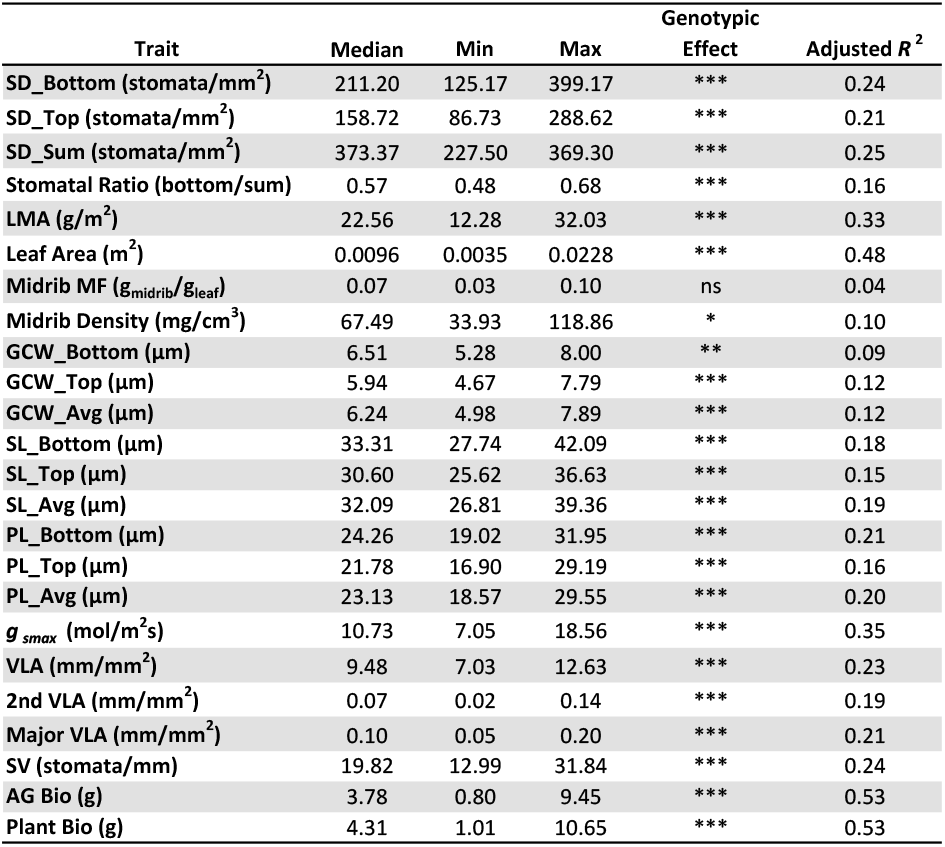
List of all traits measured along with the mean and range of trait values. Significance for genotypic effects (***P ≤ 0.001, **P ≤ 0.01, * P ≤ 0.05, ns = not significant) and adjusted R^2^ from the model are also presented. SD = stomatal density; LMA = leaf mass per area; MF = mass fraction; GCW = guard cell width; SL = stomatal length; PL = stomatal pore length; g_smax_ = theoretical maximum stomatal conductance; VLA = vein length per area; SV = stomata per vein length; AG = aboveground; Bio = biomass.

Trait correlations were examined using both bivariate and multivariate analyses. Stomatal density and size traits from the top and the bottom of each leaf were strongly correlated with each other (Figure 3a; Figure S4; e.g., SD_Top vs. SD_Bottom, PL_Top vs. PL_Bottom); as such, stomatal sum (SD_Top + SD_Bottom) and averages of the top and bottom for other traits were used for analyses moving forward. Bivariate analysis revealed numerous significant trait correlations among stomatal and vein traits (Figure 2; Figure 3). Notably, stomatal density and length were negatively correlated (Figure 3b), stomatal density and VLA were positively correlated (Figure 3c), and stomatal length and VLA were negatively correlated (Figure S4). VLA was positively correlated with both average stomatal density and theoretical *g_smax_* (Figure 3d). The only traits that were significantly correlated with leaf area were second VLA and major VLA and biomass traits indicating that trait scaling with leaf size did not play a major role in observed patterns of anatomical variation. For all bivariate plots see Figure S4.

**Figure 2:**
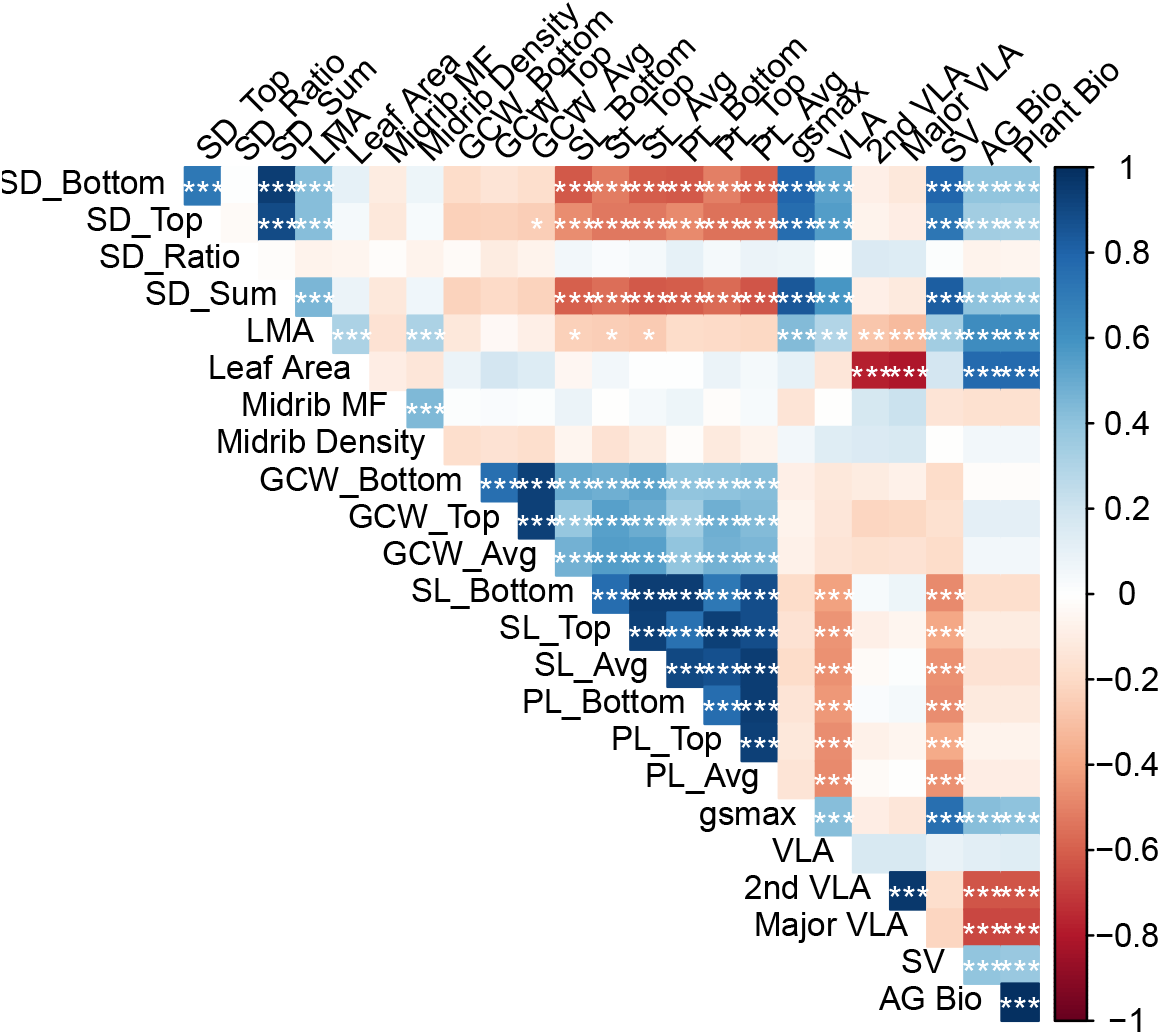
Correlation matrix of leaf traits. Values were calculated using genotype marginal means and significance tests were corrected for multiple comparisons using a Bonferroni correction. Positive correlations are in blue and negative correlations are in red. Shading gives a relative indication of the magnitude of the estimate. ***P ≤ 0.001, **P ≤ 0.01, * P ≤ 0.05. Trait abbreviations are as defined in Table 1.

**Figure 3:**
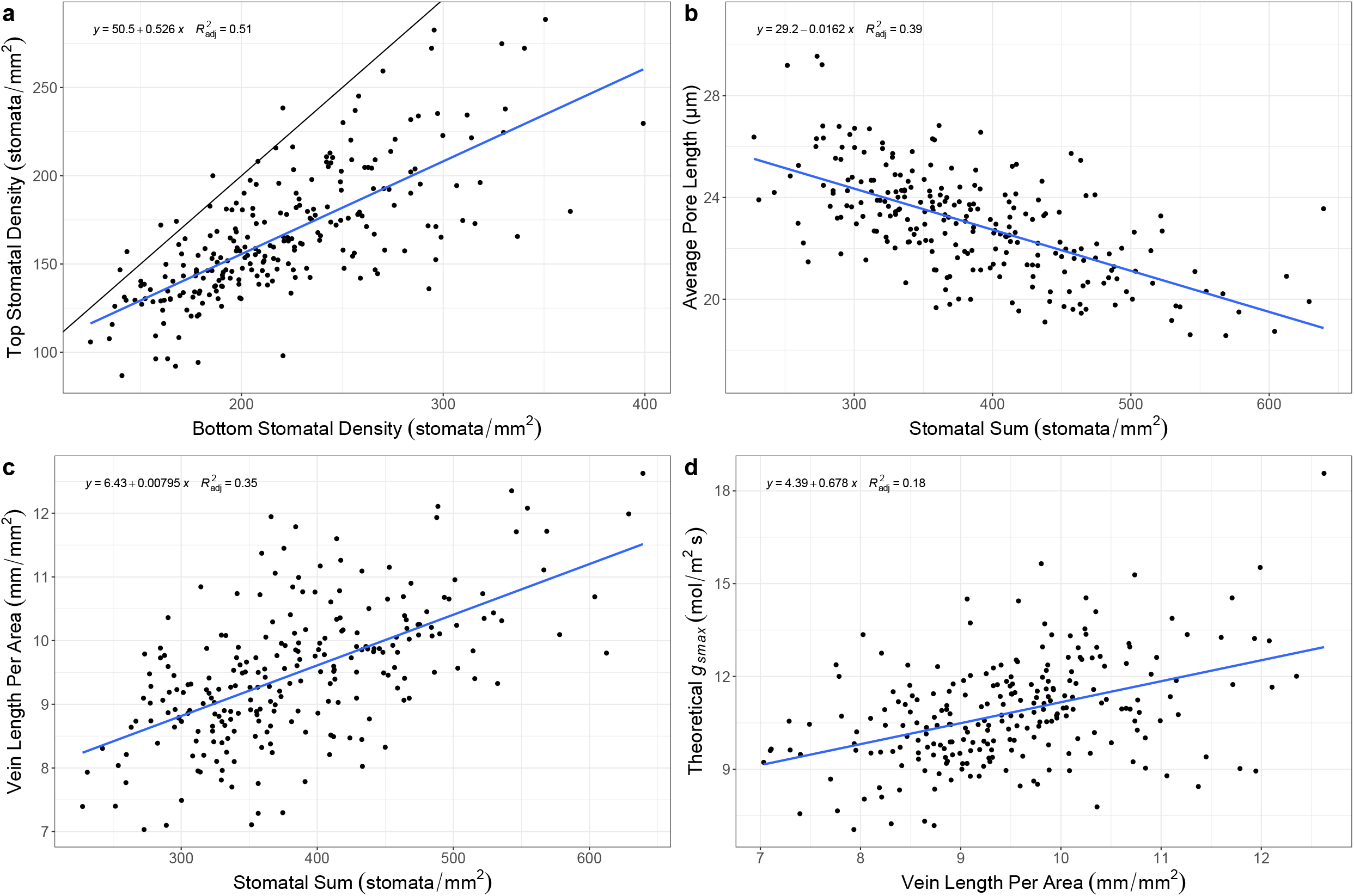
Example bivariate trait plots. In all panels, the blue line is the best fit. (a) Top vs. bottom stomatal density (black line is 1:1). (b) Average (top and bottom) stomatal pore length vs. stomatal sum. (c) Vein length per area vs. stomatal sum. (d) Theoretical stomatal conductance (g_smax_) vs. vein length per area.

Multivariate trait correlations were analyzed via PCA to determine major axes of variation. Because stomatal trait values on the top and bottom of the leaf were strongly correlated, we again used an average of the top and bottom values for stomatal size traits including length, pore length, and guard cell width (Figure 4; see Figure S5 for a full PCA including the top and bottom traits separately). The first three PCs combined to explain 62.6% of trait variation (Figure S6) with PC1 explaining 30.3%, PC2 explaining 22.6%, and PC3 explaining 10.0% of variation. The top three traits contributing to each of the major axes were: PC1 – stomatal density (SD_Sum), stomata per vein length (SV), and whole plant biomass (Plant Bio); PC2 – 2nd VLA, Major VLA, and leaf area; and PC3 – midrib density, midrib mass fraction, and LMA (Table 2). While all traits have a loading score on each PC axis, we sought to infer some functional significance for each of the major axes of variation. Given the observed trait loadings, PC1 appears to be most heavily influenced by traits related to gas exchange functioning such as stomatal density sum and stomata per vein length. In contrast, PC2 is strongly influenced by traits related to hydraulic functioning including major VLA and second VLA. Finally, PC3 is heavily influenced by traits related to cost of leaf construction including LMA and midrib traits.

**Figure 4:**
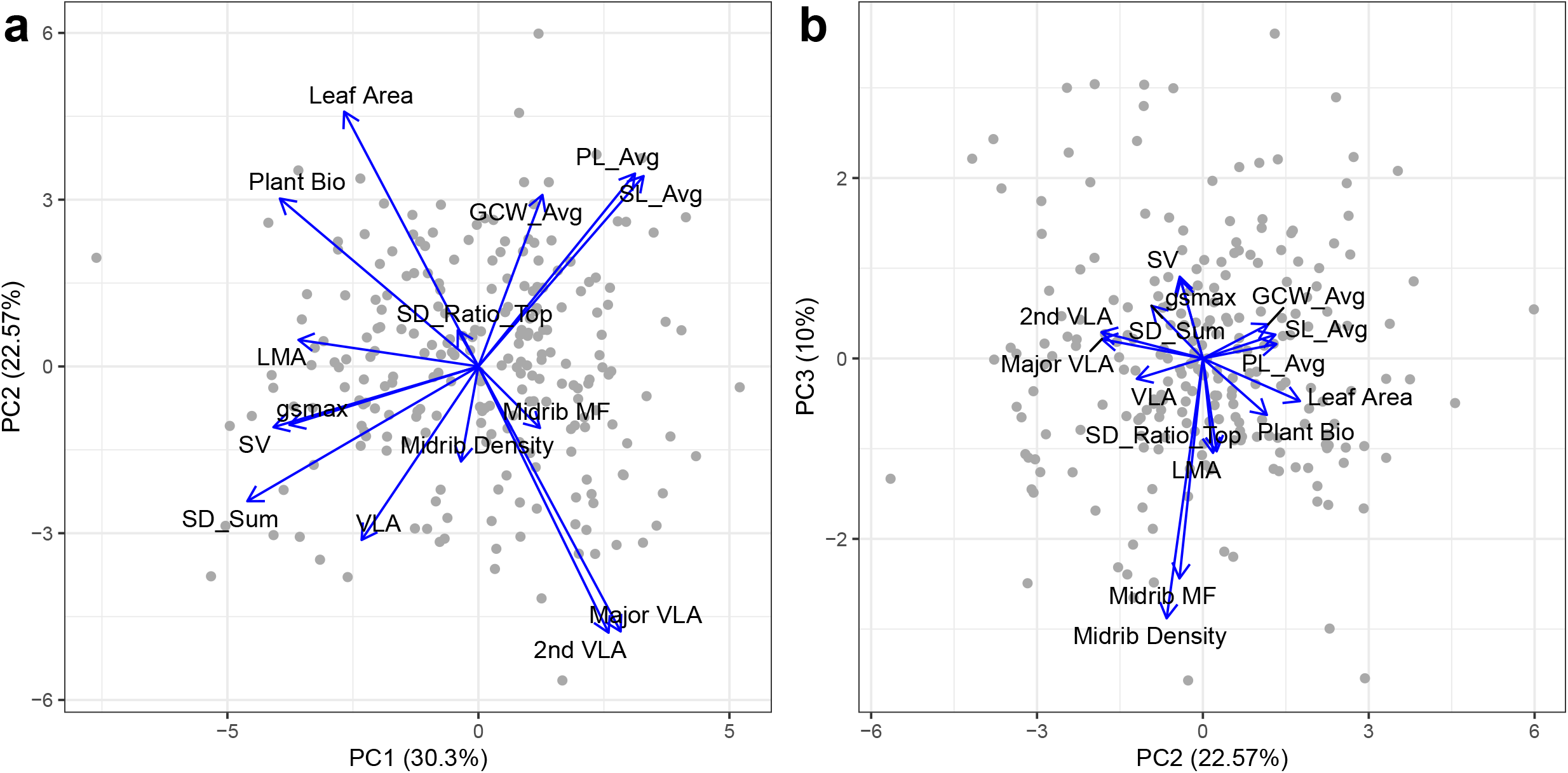
Principal component analysis (PCA) of leaf traits using estimated marginal means for each trait. (A) PC1 vs. PC2. (B) PC2 vs. PC3. Traits names that include _Avg (stomatal length [SL], pore length [PL], guard cell width [GCW]) are averages of the values from the top and bottom of the leaf and SD_Sum is the sum of stomatal density from the top and bottom. Trait abbreviations are as defined in Table 1.

**Table 2:** Trait loadings (fraction of trait variation explained by principal component) of first three principal components (PC). The top three traits per PC are highlighted in bold and with the ranking of the traits for each PC in parentheses. Trait abbreviations are as defined in Table 1.

| Traits | PC1 | PC2 | PC3 |
| --- | --- | --- | --- |
| SD_Sum | **16.08 (1)** | 4.46 (9) | 1.76 (8) |
| SD_Ratio | 0.13 (14) | 0.32 (14) | 5.48 (4) |
| LMA | 9.75 (5) | 0.17 (15) | **5.66 (3)** |
| Leaf Area | 5.42 (9) | **15.94 (3)** | 1.16 (9) |
| Midrib MF | 1.14 (13) | 0.93 (11) | **30.73 (2)** |
| Midrib Density | 0.09 (15) | 2.22 (10) | **42.88 (1)** |
| GCW_Avg | 1.23 (12) | 7.23 (7) | 0.74 (10) |
| SL_Avg | 8.21 (6) | 8.90 (5) | 0.36 (12) |
| PL_Avg | 7.38 (7) | 9.13 (4) | 0.13 (15) |
| *g<sub>smax</sub>* | 10.75 (4) | 0.84 (13) | 3.89 (6) |
| VLA | 4.11 (11) | 7.39 (6) | 0.26 (13) |
| 2nd VLA | 5.10 (10) | **17.40 (1)** | 0.44 (11) |
| Major VLA | 6.09 (8) | **17.24 (2)** | 0.24 (14) |
| SV | **12.65 (2)** | 0.91 (12) | 4.25 (5) |
| Plant Bio | **11.87 (3)** | 6.92 (8) | 2.03 (7) |

### Genetic architecture of observed trait variation

As a first step toward understanding the genetic basis of variation in leaf anatomical traits, we estimated narrow sense heritability for all traits under consideration (Table 3). Midrib taits and VLA (major and minor) had very low heritabilities (0.05 to 0.11) and the confidence intervals of these estimates all overlapped with zero. Stomatal traits had somewhat higher heritability, ranging from 0.08 to 0.20. Heritability values largely reflected our GWA results (see below) in those traits with the lowest heritability estimates had few significant associations.

**Table 3:**
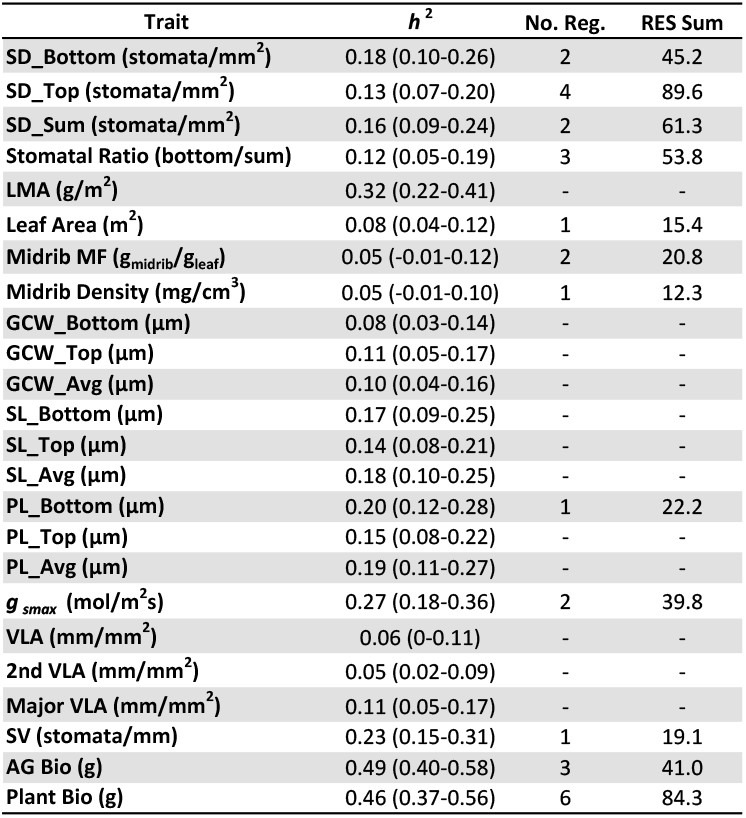
List of all traits measured along with estimated narrow sense heritabilities, numbers of significantly associated regions, and total relative effect sizes (RES Sum). Trait abbreviations are as defined in Table 1.

Our GWA analyses revealed significant associations for 12 out of 24 traits and 2 of the 3 major PC axes (Table 3, Figure 5). There were suggestive associations for several other traits, as well (Figure 5c; Figure S8). Based on observed patterns of LD, we identified a total of 24 independent genomic regions with a significant effect on one or more traits (Figures 5c and S7). Trait co-localization within these regions varied. For example, there were multiple regions on chromosome 11 that associated with both aboveground plant biomass and whole plant biomass, and a region on chromosome 3 (i.e., region 03-01) was significantly associated with *g_smax_* while being suggestive for stomatal density and VLA. Interestingly, PC1 (traits primarily related to gas exchange) had a single significant association (on chromosome 12; region 12-01) but no suggestive associations; moreover, this single region was not suggestive for any other traits (Figure 5a). Stomata per vein length (SV), a top contributor to PC1, had a single significant association along with four suggestive regions, none of which corresponded to PC1 (Figure 5b). Similarly, PC3 had a single significant association (on chromosome 3; region 03-02) that did not correspond to any other traits.

**Figure 5:**
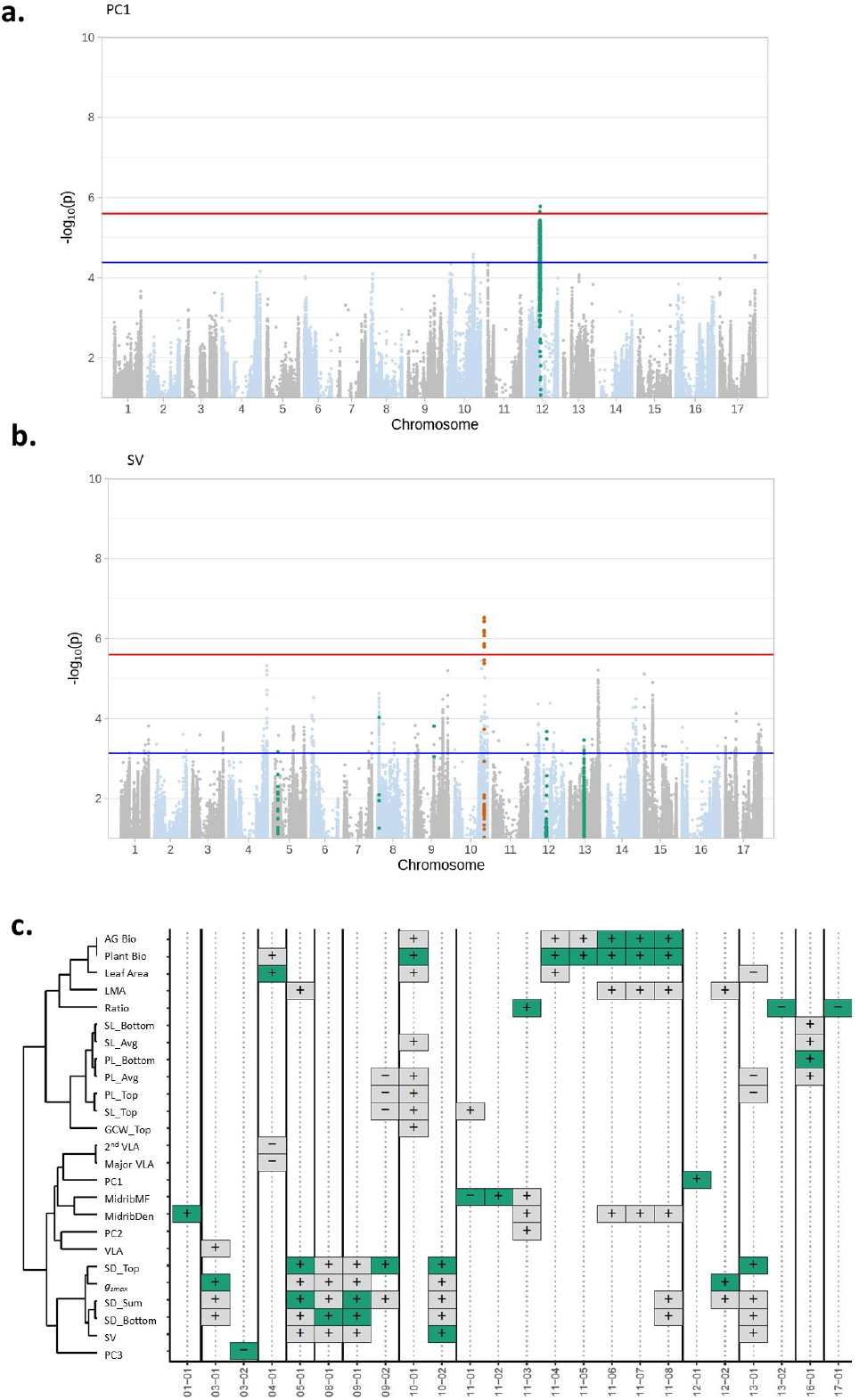
Examples and summary of GWA results. (a) Manhattan plot of PC1 showing the single significantly associated region on chromosome 12. (b) Manhattan plot of stomata per vein length (SV) showing the single significantly associated region on chromosome 10 and suggestive associations on chromosomes 5, 6, 8, and 13. In both plots, the red line is the significance threshold based on the modified Bonferroni correction and the blue line is the suggestive threshold based on the top 0.1% of all SNPs. Differently colored dots represent all SNPs in a region that are significant or suggestive for at least one trait. (c) Visual summary of all GWA results highlighting colocalization within regions. The dendrogram to the left is based on hierarchical clustering of trait correlations. Green and gray boxes indicate significant or suggestive associations with a given trait, respectively. The sign (+/-) refers to the sign of β (the effect of the minor allele on the trait value). Regions are numbered numerically within chromosomes and box/region sizes are arbitrary. Ratio refers to the stomatal ratio. Trait abbreviations are otherwise as defined in Table 1.

Overall, we found somewhat limited evidence for trait colocalization despite the prevalence of significant trait-trait correlations (Figure 5c). Indeed, no region was significantly associated with more than 1-2 traits. However, the inclusion of suggestive associations revealed more support for a common genetic basis. Traits with the largest number of colocalizations were stomatal size and density-related traits. For example, region 13-01 had seven associations including a significant association for stomatal density (top) and suggestive associations for leaf area, pore length (top and average), stomatal density (bottom and sum), and SV (Figure 5c). No significant associations were identified for VLA, although there was a suggestive association in region 03-01, which was significant for *g_smax_* and suggestive for stomatal density (both bottom and sum). Allelic effects in this case were consistent with observed trait correlations, with an increase in *g_smax_* being associated with an increase in VLA and stomatal density.

Relative effect sizes (RES) for individual associations ranged from 9% to 35% of the observed range of trait variation. Summing across regions, traits with significant associations had total RES values of ca. 12-90%. Stomatal density top had the largest amount of variation explained at 89.6% and midrib density the least with 12.3%. Significantly associated regions varied in size, ranging from a single SNP to 6.52 Mbp, though they tended to cluster at the lower end of the range with the majority being < 200 kbp (mean = 592.7 kbp, median = 77.54 kbp; Table S2). These regions contained anywhere from 1 to 135 genes; here again, the significant regions tended to cluster at the lower end of this range (mean = 12.3 genes, median = 2 genes; Table S2). The most gene-rich regions (i.e., 10-01, 12-01) were significantly associated with variation in plant biomass and PC1, respectively (Figures 5C, S4, and S9). A full list of gene names and annotations is available in Table S2.

## Discussion

Stomata and leaf veins are central to plant-water relations and thus potentially important players in determining the performance of plants under water stress. Here, we investigated patterns of variation and covariation in stomatal and vein traits across a diversity panel of inbred cultivated sunflower breeding lines with the goal of improving our understanding of covariation between these traits and their underlying genetic basis. We additionally sought to determine the extent to which observed trait correlations result from genomic co-localization which would indicate that trait relationships are genetically constrained and difficult (if not impossible) to disrupt in the interest of producing novel combinations. In quantifying variation in stomatal and vein traits, we observed numerous correlations amongst traits (Figure 2 and 3). Across traits, we identified three primary axes of variation that we interpreted as being most heavily influenced by traits involved in gas exchange, hydraulics, and leaf construction (Figure 2; Table 2). Subsequently, we performed GWA analyses to examine the genetic architecture of these traits and axes. We found somewhat limited overlap in significant genomic associations across traits, though we did identify significant associations for two of the three multi-trait axes (Figure 5; Table 3).

### Patterns of phenotypic variation and trait correlations

Leaf anatomical traits are known to vary widely across species and environments. For example, stomatal length has been shown to vary globally between 10-80 μm with density varying between 5-1000 stomata/mm^2^ (Shahinnia *et al*., 2016; Hetherington and Woodward, 2003). Additionally, a global dataset of 796 species revealed broad variation in estimates of VLA, ranging from 0.1-24.4 mm/mm^2^ (Sack and Scoffoni, 2013). In cultivated sunflower, we documented substantial variation in stomatal size and density as well as VLA. Indeed, observed trait values covered 14% (227.5-369.3 stomata/mm^2^) of the global range in stomatal density, 18% (26.8-39.4 μm) of the global range in stomatal length, and 23% (7.0-12.6 mm/mm^2^) of the global range in VLA (Table 1). While there is a general lack of large datasets describing intraspecific variation in these sorts of traits, particularly for VLA, the ranges that we observed in cultivated sunflower appear quite wide. For example, a global collection of 62 wild accessions of *Arabidopsis thaliana*, grown under benign conditions, covered just 4.3% (17-59 stomata/mm^2^) of the global range of stomatal densities (Delgado *et al*., 2011) while a collection of 330 accessions from across the European range of the same species covered less than 12% (87-204 stomata/mm^2^) of the global range (Dittberner *et al*., 2018). The wide range of variability observed herein is perhaps even more noteworthy given that cultivated sunflower has experienced genetic bottlenecks associated with domestication and improvement that reduced levels of genetic variability as compared to its wild progenitor (Liu and Burke, 2006; Mandel *et al.*, 2011; Park and Burke, 2020).

When analyzed together, leaf anatomical traits exhibited many significant bivariate trait correlations that generally followed expectations based on the literature and their known roles in plant-water relations (Sack and Scoffoni, 2013; Doheny-Adams *et al*., 2012; Shahinnia *et al*., 2016; Figure 2). For example, our data showed that stomatal density and VLA are positively correlated. This was expected as there tends to be a balance between stomata and veins such that water use and carbon acquisition are optimized (Carins Murphy *et al*., 2014; Brodribb *et al.*, 2007; Sack and Scoffoni, 2013). Additionally, stomatal size and density were negatively correlated, as expected, since the total area allocated to stomata affects total stomatal conductance and thus total photosynthesis (Harrison *et al*., 2019; Shahinnia *et al*., 2016; Figure 2) both within (Doheny-Adams *et al*., 2012) and across species (Hetherington and Woodward, 2003). Overall plant size, estimated as biomass, correlated positively with stomatal density and related traits (e.g., *g_smax_* and SV) but negatively with 2nd/major VLA, with larger plants tending to have higher stomatal density and lower 2nd/major VLA. Conversely, stomatal size was unrelated to plant biomass despite its correlation with other leaf traits of interest. Besides scaling with plant mass (Wang *et al*., 2020), the potential scaling of traits with leaf area is of interest as correlations can arise as a byproduct of trait values scaling with size. Given that stomatal and vein traits were not significantly correlated with leaf area, however, it appears that observed correlations between these traits exist independently of variation in leaf size (Figure 2).

When compared to minor VLA, lower order vein traits (i.e., 2nd and major VLA) exhibited a distinct pattern of trait correlations. These traits were not significantly correlated with any stomatal traits; rather, they exhibited significant correlations with traits related to the investment in leaf production (i.e., LMA and leaf area; Figures 2 and 3). Contrary to expectations based on cross-species comparisons (e.g., Walls, 2011; Kawai and Okada, 2016), 2nd and major VLA were negatively correlated with LMA (Figure 2) suggesting that variation in LMA at this scale may be driven by other, perhaps unmeasured traits such as leaf thickness. Similarly, 2nd and major VLA were negatively correlated with leaf size. This pattern was, however, expected given that major veins are typically formed early in leaf development before being pushed apart as leaf expansion accelerates (Sack and Scoffoni, 2013). In contrast, minor veins are expected to show no such relationship (consistent with our results) because they can be initiated throughout leaf development.

As compared to bivariate analyses, multivariate analyses provide a more holistic view of trait relationships along with possible impacts of leaf anatomical variation on ‘higher level’ traits such as biomass, leaf size, and LMA. When analyzed via PCA, nearly two-thirds of the observed trait variation was captured by the first three PC axes. As noted above, these axes are most heavily influenced by suites of traits involved in gas exchange, hydraulics, and leaf construction. More specifically, plants with lower stomatal density (estimated as stomatal sum) and fewer stomata per vein length (SV), which are primary players in stomatal conductance, tended to be smaller overall. In terms of hydraulic traits, and consistent with the results of our bivariate analyses, plants with greater second and major VLA tended to have smaller leaves. Interestingly, this trait combination is thought to confer greater leaf drought tolerance (Scoffoni *et al*., 2011). Finally, in terms of leaf construction traits, plants that produced more costly leaves (i.e., leaves with higher LMA) tended to have a greater relative investment in major structural features including midrib density and midrib mass fraction (Figure 4; Table 2), likely reflecting an increase in the mechanical strength of such leaves (Méndez-Alonzo *et al*., 2013).

### Genetic architecture of observed trait variation

Estimates of narrow sense heritability ranged from 0.05 to 0.49 across traits. Notably, vein (including midrib) traits had low heritability estimates while stomatal traits had higher estimates (Table 3). The highest estimates were for biomass-related traits, LMA, and *g_smax_* indicating a more substantial contribution of additive genetic effects to observed variation in these traits as compared to others (Kruijer *et al*., 2015). Somewhat surprisingly given the highly significant effect of genotype on VLA, the heritability estimate for that trait was not significantly different from zero (Tables 1 and 3), indicating little to no contribution of additive genetic effects to observed variation. Midrib density and midrib mass fraction had similarly low heritability estimates, though the evidence of a genotypic effect on the former was less clear, and genotype had no apparent effect on the latter. Interestingly, heritability estimates for the composite trait SV (i.e., stomata per vein length) were noticeably higher than estimates for vein traits alone indicating an additive genetic component of the observed variation in this trait. Collectively, these results suggest that traits with the lowest heritability estimates have limited potential for improvement via breeding while others are likely to be more amenable to such efforts.

Consistent with our heritability estimates, the two biomass-related traits exhibited the largest number of significant associations in our GWA analyses (Table 3; Figure 5c; Figure S8). These traits colocalized with leaf area, midrib density, and LMA but not with any other vein or stomatal traits. Similarly, we identified significant associations for multiple stomatal traits, including stomatal density (bottom, top, and sum) and pore length (bottom), many of which colocalized with suggestive associations (i.e., SNPs with −log_10_(p) values in the top 0.1%) for other stomatal traits. Consistent with the low heritability estimates for traits related to vein density (i.e., VLA, 2nd VLA, and major VLA), no significant genomic associations were found for any of these traits. There was, however, one suggestive association for VLA that colocalized with a significant association for *g_smax_* (region 03-01) and suggestive associations for stomatal density (bottom and sum), consistent with a presumed functional relationship between these traits. Despite the lack of significant associations for vein-related traits considered on their own, we identified one significant and five suggestive associations for the composite trait SV (i.e., stomata per vein length). These regions tended to colocalize with stomatal traits suggesting that variation in stomatal characteristics is the primary driver of this trait relationship (Figure 5c, Table S1).

Taken together, our trait-by-trait analyses revealed limited evidence for colocalization between stomatal and vein traits despite the existence of widespread and significant correlations between such traits. While this result could be due, at least in part, to the high stringency of our significance threshold and thus the failure to detect true positives – a common challenge in GWA analyses (Gupta *et al*., 2019) – the identification and inclusion of suggestive regions in our analyses should have helped to mitigate this issue. Nonetheless, observed trait correlations did not appear to be accompanied by clear patterns of genomic colocalization on a single trait basis suggesting a largely independent genetic basis of our traits of interest. However, when multivariate trait relationships were taken into account, our GWA analyses revealed significant associations for two of the three major PC axes (i.e., PC1 and PC3) and a suggestive association for the third (i.e., PC2). For PC1, which is most heavily influenced by traits related to gas exchange (i.e., stomatal density and SV) along with plant biomass, the single significant association (i.e., region 12-01) colocalized with suggestive associations in 2nd VLA and SV. In contrast, the analysis of PC3, which had midrib density and mass fraction as well as LMA as its top three contributors, identified a novel association (i.e., region 03-02). This region was not identified as being significantly or suggestively associated with any of the individual traits analyzed herein, illustrating the potential value of employing a multivariate approach to GWA analyses (see also, e.g., Yano *et al*., 2019; Ma *et al*., 2021). These results also highlight the challenges associated with genetically decoupling certain traits to produce novel phenotypic combinations even though individual trait analyses revealed largely independent genetic architectures.

In terms of effect sizes, the significantly associated regions that we identified tended to have had small to moderate effects (estimated as RES) on trait values. In fact, only three trait/region combinations individually accounted for > 25% of the observed range of trait values across the population (stomatal density [top and sum] in region 05_01 and stomatal density [sum] in region 09_01; Table S1). This result is perhaps not surprising given the relatively low heritability estimates observed for many traits. It is worth noting, however, that the traits with the highest heritability estimates (i.e., biomass-related traits and LMA) tended to be associated with regions of relatively minor effect (i.e., RES < 15%) suggesting that they have a complex genetic basis.

In contrast, stomatal traits were associated with some of the largest RES values, suggesting the presence of genes of larger effect and a simpler genetic basis overall. This result was mirrored in the multivariate analyses with the single association underlying PC1 (region 12-01), which is heavily influenced by stomatal traits, accounting for nearly 21% of the observed range of variation in this ‘trait’ across the population. Unfortunately, most of the genes contained within the significantly associated regions identified herein did not yield obvious candidates for the traits of interest.

While many of the genes in regions of interest were annotated as hypothetical proteins or otherwise showed no clear connection with leaf anatomy, two regions did contain potential genes of interest. Region 10-01, which is significant for total biomass and suggestive aboveground biomass, leaf size, and several stomatal size traits (Figure 5b), includes a gene annotated as a *Putative transcription factor SSXT* (Ha412HOChr10g0435021; GO:0048366 [leaf development]; Table S2). Members of this gene family are thought to play a role in cell size determination in leaves in *Arabidopsis* (Nozaki *et al*., 2020). This is, however, one of the one of the larger regions that we identified and contains 135 genes so, while this gene appears to be a promising candidate for one or more of the size-related traits that map to this region, this result should be interpreted with caution until functional evidence is available to support its effect on one or more of the associated traits. Nonetheless, it is interesting to note that epidermal cell size is also known to be negatively associated with vein and stomatal densities such that smaller epidermal cells facilitate greater stomatal and vein densities (Brodribb *et al*. 2013; Murphy *et al*. 2017; Simonin and Roddy 2018). The other region containing a potential gene of interest, 11-01, contains a *Putative Epidermal Patterning Factor-like protein (EPF;* Ha412HOChr11g0479421; GO:0010052 [guard cell differentiation]; Table S2). This region is significant for MidribMF and suggestive for top stomatal length. EPFs are known to be involved in the density of guard cells and epidermal cells (Hara *et al*., 2009), but how this might relate to midrib mass fraction is unclear. Establishment of a (potential) role for these genes in producing variation in any of the leaf anatomical traits analyzed herein awaits further investigation. Nonetheless, the genomic regions identified during the course of this work, particularly those with larger effects, represent potential targets for future efforts aimed at modifying leaf anatomical traits in sunflower.

## Experimental Procedures

### Plant Material

The cultivated sunflower lines analyzed in this study comprise the sunflower association mapping (SAM) population (Mandel *et al*., 2011), which includes 288 inbred lines that capture ca. 90% of allelic diversity in crop sunflower (Mandel *et al*., 2013). This population has since been subjected to whole genome re-sequencing, thereby enabling the identification of a genome-wide collection of single nucleotide polymorphisms (SNPs) from the full set of lines (Hübner *et al*., 2019).

### Experimental Design

In the summer of 2017, 239 inbred lines from the SAM population (4 four replicates each; N = 4 x 239 = 956 total individuals) were grown in the greenhouse in a randomized block design. The plants used in this study correspond to the control plants from Temme *et al*. (2020) and detailed plant growth methods are described therein. Briefly, 239 of the 288 lines in the mapping population were used due to greenhouse space constraints and to remove lines with greater than expected levels of heterozygosity. Following germination, all plants were grown for one week in seedling trays to allow for establishment before being transplanted into 2.83 L pots (TP414; Stuewe & Sons, Tangent, OR) filled with a 3:1 mixture of sand and a calcined clay mixture (Turface MVP, Turface Athletics). Pots were fertilized with 40g Osmocote Plus (15-12-9 NPK; ScottsMiracle-Gro, Marysville, OH) and 5 mL each of gypsum (Performance Minerals Corporation, Birmingham, AL) and lime (Austinville Limestone, Austinville, VA) powders for supplemental Ca^2+^. All pots were well-watered, and plants were allowed to grow for three additional weeks before being harvested at four weeks old. Plants were grown under typical summer temperatures and natural light levels in Georgia. At harvest, biomass was collected and dried in ovens at 60°C for at least 72 hours. Roots were washed and dried in the same manner. Dried samples were weighed to calculate total and aboveground biomass. During harvest, the two most recently fully expanded leaves (MRFEL) were also collected from each plant. One leaf was arbitrarily designated for use in the estimation of LMA and the other was used for leaf anatomy analyses. Due to our focus on sampling at an equivalent stage during leaf development, we chose the MRFEL at the time of harvest such that the specific leaf pair varied across the population (though generally leaves came from the top 90% of the plant with the number of underdeveloped leaves above varying). The adaxial and abaxial (hereafter top and bottom) surfaces of one half of one MRFEL per plant were pressed into dental putty (President Dental Putty; Coltène/Whaledent Inc., Cuyahoga Falls, OH) to create an impression of the epidermis of each leaf surface to allow for the visualization and analysis of stomatal traits following the general methods of (Weyers and Johansen, 1985). The other half of the same leaf was stored in Formalin-Acetic Acid-Alcohol (FAA) fixative for imaging and analysis of vein traits.

To estimate LMA, the designated MRFEL from each plant was scanned on a flatbed scanner at 300 dpi (Temme *et al*., 2020). This image was then used to calculate leaf area using ImageJ v1.52b (Schindelin *et al*., 2012). The leaf was then dried at 60°C for 48 hours and weighed (the petiole was not included). Using both the mass and area measurements, LMA was calculated as LMA = dry mass/unit area (g/m^2^).

For stomatal traits, clear nail polish was applied to the epidermal impressions of the top and bottom leaf surfaces and subsequently peeled off using clear tape and placed on microscope slides (Hilu and Randall, 1984; Weyers and Johansen, 1985). Slides were imaged using a Zeiss Axioskop 2 microscope along with ZEN software (Carl Zeiss Microscopy) under the 100X objective to enable estimation of stomatal size. A second set of images (four different fields of view per impression) were taken using the 20X objective to enable estimation of stomatal density. Size estimates were based on 10 stomata per leaf, separately for the top and bottom surfaces of each leaf, for a total of 20 stomata per plant. Stomatal length, pore length, and guard cell width were measured for each stomate using ImageJ (Figure 1a). Stomatal densities were estimated by counting the number of stomata in each of the four fields of view (counting partial stomata on only two sides of each image) per side of each leaf (eight images total per plant). Stomatal ratio was then calculated as number of bottom stomata/total stomata and stomatal sum was calculated as number of top stomata + number of bottom stomata. For consistency with the literature, we used stomatal sum instead of average density of stomates (e.g., Muir, 2018; Richardson et al., 2020). Finally, maximum stomatal conductance (*g_smax_*), the theoretical maximum rate of gas exchange if all stomata were fully open (calculated as sum of top and bottom), was calculated based on stomatal density and size measurements following the approach of Dow *et al*. (2014). This was used instead of directly measuring *g_s_* since direct measurements were not feasible for such a large sample size.

For vein traits, the half of each leaf that was fixed in FAA was cleared and stained for analysis using a modification of established procedures (Berlyn *et al*., 1976; Scoffoni and Sack, 2013). Leaves were cleared in 5% NaOH at 55°C for 5-7 days. Subsequently, leaves were rinsed with deionized water and run through an ethanol dilution series of 30%, 50%, and 70% to dehydrate. After dehydration, leaves were stained with a 0.01% safranin dye solution for 30 minutes to make the veins more visible. Images of the stained leaves were captured using both a flatbed scanner at 2400 dpi to image the entire leaf half and a microscope (Zeiss Axioskop 2) using the 5X objective on a small section of leaf with three to four different fields of view per leaf (Vasco *et al.*, 2014; Figure 1 a-b). Second-order vein length was measured by manually tracing veins that branched off the midrib (primary vein) and joined together in arches (Ellis *et al*., 2009). Major vein length was estimated by adding the length of the midrib to the second-order vein length.

Minor vein lengths were estimated from a subset of the microscopic images by manually tracing the veins with ImageJ. The subset that was manually traced was then used to train a deep learning algorithm (see below) to process the rest of the images. The results of these analyses of minor veins were used to calculate vein length per area (VLA). Midrib density (expressed as mg/cm^3^) was estimated from the mass of the midrib and an estimate of its volume; the latter value was calculated from the length of the midrib and its diameter at the base of the leaf and assuming a conical shape. Midrib mass fraction is the ratio of mass of the midrib to mass of the leaf. A composite trait of stomata number per vein length was also calculated as SV = average stomatal density (i.e., calculated as the average of the top and bottom density estimates) divided by VLA (Zhao et al., 2017).

### Image Segmentation Using a Neural Network

A modified version of the U-Net (Xu *et al*., 2020; Ronneberger *et al*., 2015) deep neural network was used to segment leaf-vein pixels from background pixels (Figures S1-S3). Network structure, hyperparameter tuning, and training details are included in Methods S1. Briefly, a test set of 85 images was created by randomly selecting one image from each of 85 randomly selected (without replacement) genotypes. These images were not used during training of the network or hyperparameter estimation. All remaining, manually traced (i.e., hand-segmented) images were used in the training set (747 images). Training was performed on randomly selected 572 pixels wide x 572 pixels high x 3 color (RGB) channels regions of the bright-field leaf images. Each region was normalized to a mean of 0 and a standard deviation of 1. The network was trained with a batch size of 1 (Ronneberger *et al*., 2015) for 1900 batches with a learning rate of 10^-4^.

Per-pixel in-vein predictions were performed by mirror padding each full size (2584×1936 pixel) bright field image, normalizing to a mean of zero and standard deviation of one, and passing the images through the trained network. To reduce the creation of small, non-real branches during medial-axis thinning, the probabilities were filtered with a Gaussian kernel with a standard deviation of 12, about the width of a typical vein. The smoothed images were segmented with a cutoff of 0.2. The resulting segmented images were thinned to single-pixel-wide lines using the medial-axis transform (Bucksch, 2014). Vein lengths were then calculated as the sum of all pixels within the thinned line. The hand-segmented and network-segmented vein lengths of the testing set images have a Pearson correlation of 0.97 (Figure S2). All code can be found at https://github.com/aatemme/burke_leaf_veins/. For additional methodological details, see Methods S1.

### Data Analysis

All data analysis was conducted using R v3.4.3 (R Core Team, 2013). Bivariate plots were made for all pairwise comparisons using the R package *ggplot2* (Wickham, 2016) to show the range of trait variation. A two-way ANOVA with genotype and block as the main effects was performed to test for variation among genotypes and to calculate estimated marginal trait means for genotypes after removing block effects. Marginal means calculated with the R package *emmeans* (Lenth, 2020) were also used to estimate trait correlations and to create a correlation matrix using the R package *corrplot* (Wei and Simko, 2017). A principal component analysis (PCA) was conducted using the function prcomp() and the package *ggfortify* (Tang *et al*., 2016; Horikoshi and Tang, 2018) to visualize multivariate correlations. Narrow sense heritabilities (*h^2^*) were calculated using the package *heritability* (Kruijer *et al*., 2019) by inputting the kinship matrix (see below) for all genotypes and individual pot-level trait data.

All traits of interest, including the first three principal components (PCs) from the PCA, were analyzed via GWA analyses to identify genomic regions that are significantly associated with each trait. These analyses were performed using a custom pipeline as described in Temme *et al*. (2020; https://github.com/aatemme/Sunflower-GWAS-v2). SNPs used in these analyses were called as described in Hübner *et al*. (2019) and reordered based on the improved HA412-HOv2 sunflower genome assembly (Todesco *et al*., 2020). SNPs were filtered to retain those with minor allele frequency ≥ 5% and residual heterozygosity of < 10% (Temme *et al*. 2020). The GWA analyses were performed using GEMMA v.98.1 (Zhou and Stephens, 2012) to test for associations while correcting for kinship (as calculated by GEMMA) and population structure (using the first four PCs from an analysis using a subset of independent SNPs in SNPRelate; Zheng *et al*., 2012). The significance threshold was corrected for multiple comparisons using a modified Bonferroni correction based on the number of multi-SNP haplotypic within the genome as determined by an analysis of linkage disequilibrium (LD) across the population (Temme *et al*., 2020); any SNPs exceeding this threshold were considered to be significantly associated with the trait of interest. Following this step, significant SNPs were grouped into significantly associated genomic regions, with all SNPs occurring within a previously identified haplotypic block being assumed to mark a single region (see Temme *et al*., 2020 for details). Suggestive associations were then identified as SNPs in the top 0.1% of all SNPs that co-localized with significant associations for one or more other traits. The relative effect size (RES) of associated SNPs/regions were then estimated as the percentage of the observed range of variation in a particular trait that is explained by each association, as follows:

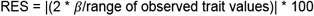

Here, *β* represents the effect of the minor allele on the trait value and the range is based on the distribution of trait values across all genotypes (Masalia *et al*., 2018). In cases of multi-SNP blocks, the RES value was estimated as the maximum value for all SNPs within that block. These values can also be summed to represent the total percentage variation explained by all significant associations for a particular trait.

## Data Statement

All raw data from the resequencing of the SAM population are stored in the Sequence Read Archive under Bioproject PRJNA353001. SNP set used and genome assembly as in Todesco *et al*. (2020). Raw phenotypic data and image files are available via Dryad at https://doi.org/10.5061/dryad.63xsj3v54.

## Acknowledgements

We thank Kelly Bettinger, Mike Boyd, Kevin Tarner, the rest of the greenhouse staff, and numerous members of the Burke and Donovan labs for help with various aspects of this research. Niki Padgett, Summerlin Courchaine, Kelly Bettinger, Nicole Reisinger, Nadia Krigger, and Jared Bennett provided invaluable assistance with the leaf image processing; Alex Bucksch provided valuable feedback on the automated image analyses; and members of the Burke and Donovan labs provided comments that greatly improved an earlier version of the manuscript. This work was supported by a grant from the NSF Plant Genome Program (IOS-1444522) to JMB as well as funding from the International Consortium on Sunflower Genomics.

## Supporting Information

***Figure S1:** Comparison of human measured, and neural network estimated vein lengths*

***Figure S2:** Network architecture used in-place of the U-Net structure*

***Figure S3**: Cross-validated training*

**Figure S4:** *Bivariate plots for all trait correlations*

**Figure S5:** *Principal component analysis (PCA) of all measured leaf traits*

**Figure S6:** *Scree plot*

**Figure S7.** *Linkage disequilibrium heatmaps for all significant SNPs found on each chromosome*

**Figure S8:** *Manhattan plots resulting from GWA analyses for all traits*

**Figure S9**: *Distribution of the number of genes per significant region*

**Figure S10**: Visualization of *haplotypic blocks across the sunflower genome*

**Table S1:** *All significant and suggestive regions underlying observed trait variation*

**Table S2:** *List of genes per region along with significant/suggestive trait associations*

**Methods S1:** *Neural Network Architecture, Training and Prediction*

