## Supplemental Figures for "Genomic regions associate with major axes of variation driven by gas exchange and leaf construction traits in cultivated sunflower (*Helianthus annuus* L.)"

***Figure S1:*** *Comparison of human measured and neural network estimated vein lengths on test-set images (N=85).*

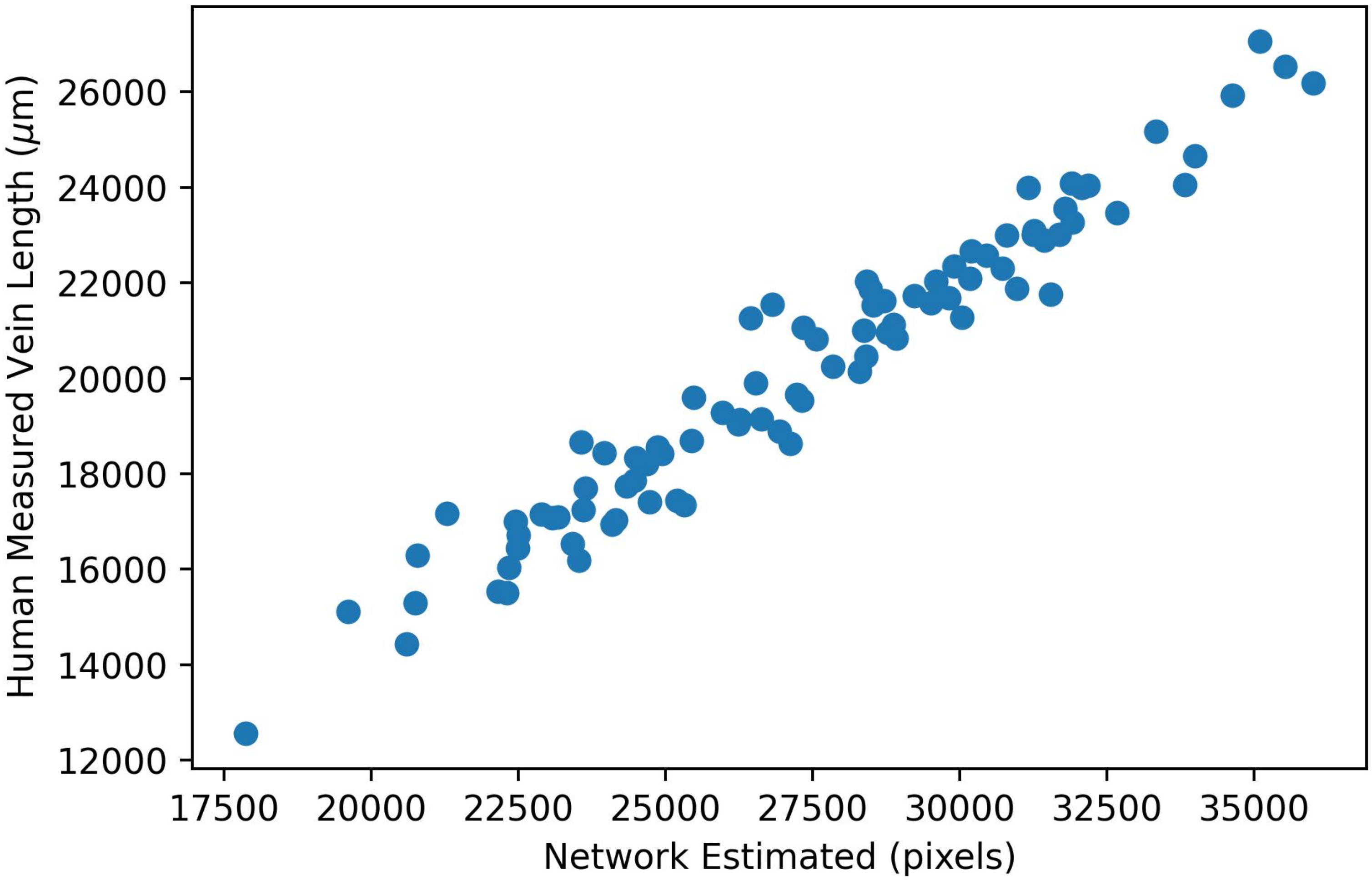
Human Measured Vein Length ($\mu$m)
26000
24000
22000
20000
18000
16000
14000
12000
17500
20000
22500
25000
27500
30000
32500
35000
Network Estimated (pixels)

***Figure S2:*** *Network architecture used in-place of the U-Net structure after the last max pool layer through the first up convolution. The 32x32x512 input tensor represents the output of the last max-pool layer. The 56x56x512 output tensor is the same as the output size of the first up convolution. + represents element-wise addition of the output tensors.*

32x32x512 pixels

| | Channels in | Channels out | Kernel | Stride | Padding | Dilation | Activation |
|---|---|---|---|---|---|---|---|
| Dilation 1 | 512 | 1024 | 3x3 | 1 | 0 | 1 | ELU |
| Dilation 2 | 1024 | 1024 | 3x3 | 1 | 2 | 2 | ELU |
| Dilation 4 | 1024 | 1024 | 3x3 | 1 | 4 | 4 | ELU |
| Dilation 8 | 1024 | 1024 | 3x3 | 1 | 8 | 8 | ELU |
| Dilation 16 | 1024 | 1024 | 3x3 | 1 | 16 | 16 | ELU |
| Dilation 32 | 1024 | 1024 | 3x3 | 1 | 32 | 32 | ELU |
| Transposed Convolution | 1024 | 1024 | 2x2 | 2 | 0 | 2 | ELU |
| Convolution | 1024 | 512 | 5x5 | 1 | 0 | 1 | ELU |

56x56x512 pixels

***Figure S3****: Cross-validated training (red) and validation (black) loss of model with final hyperparameters. Shading indicates standard error.*

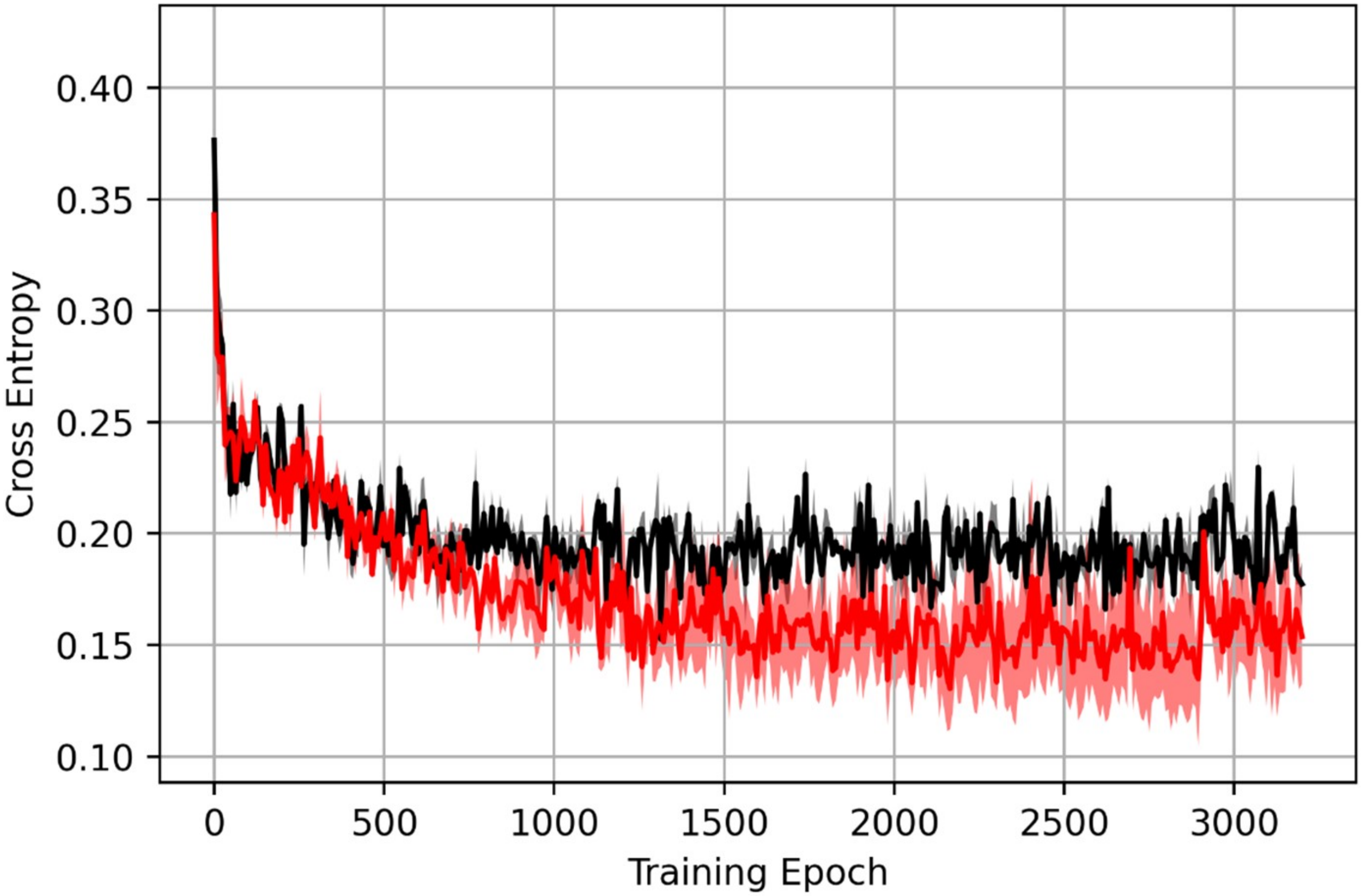

0.40
0.35
0.30
0.25
0.20
0.15
0.10
Cross Entropy
0
500
1000
1500
2000
2500
3000
Training Epoch

**Figure S4:** *Bivariate plots for all trait correlations presented in Figure 2. Points represent the estimated marginal means of genotypes. Blue line represents the fitted regression line. Units for all traits are as in Table 1.*

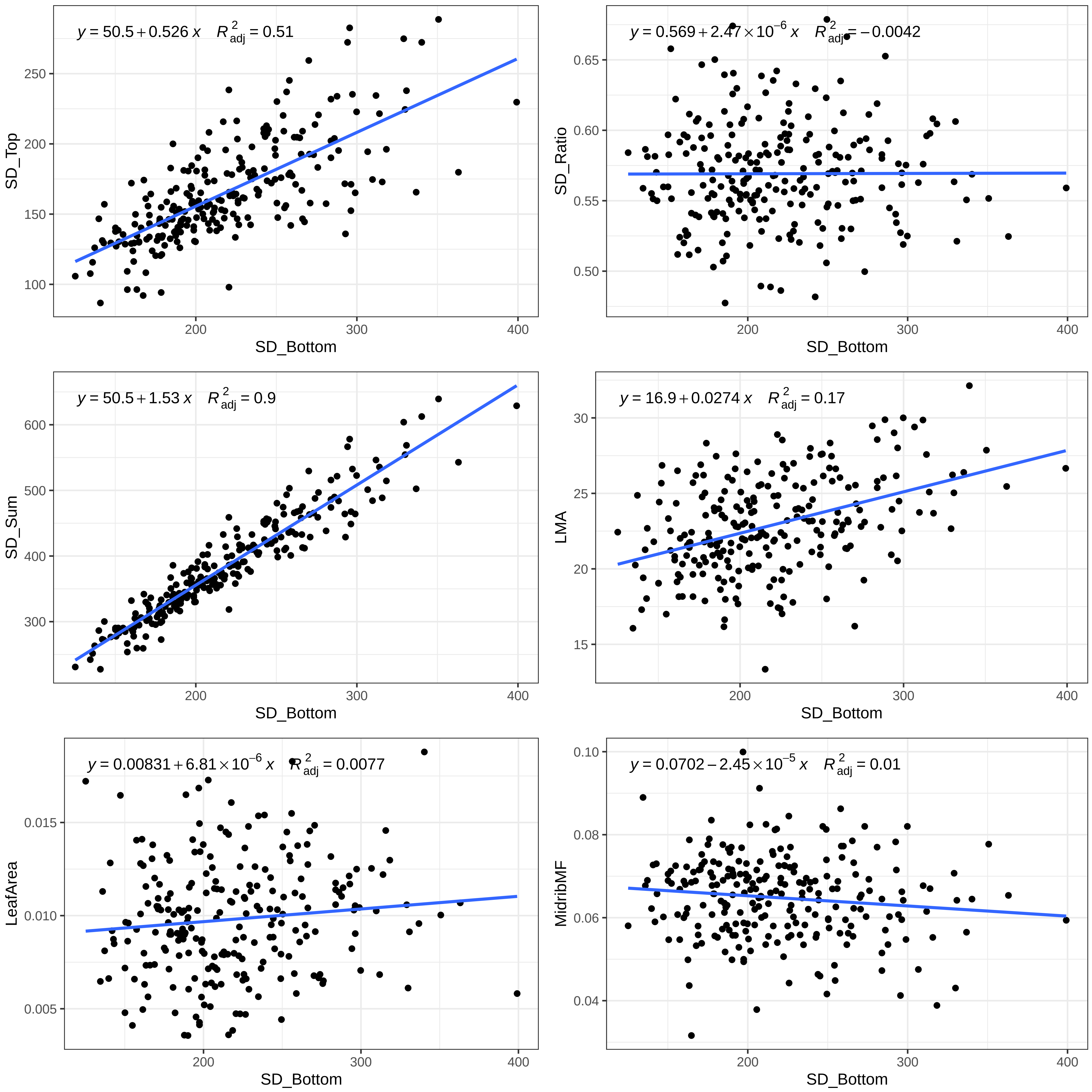

y = 50.5 + 0.526 x R2adj = 0.51
SD_Top
100
150
200
250
SD_Bottom
200
300
400
y = 0.569 + 2.47 × 10−6 x R2adj = −0.0042
SD_Ratio
0.50
0.55
0.60
0.65
SD_Bottom
200
300
400
y = 50.5 + 1.53 x R2adj = 0.9
SD_Sum
300
400
500
600
SD_Bottom
200
300
400
y = 16.9 + 0.0274 x R2adj = 0.17
LMA
15
20
25
30
SD_Bottom
200
300
400
y = 0.00831 + 6.81 × 10−6 x R2adj = 0.0077
LeafArea
0.005
0.010
0.015
SD_Bottom
200
300
400
y = 0.0702 − 2.45 × 10−5 x R2adj = 0.01
MidribMF
0.04
0.06
0.08
0.10
SD_Bottom
200
300
400

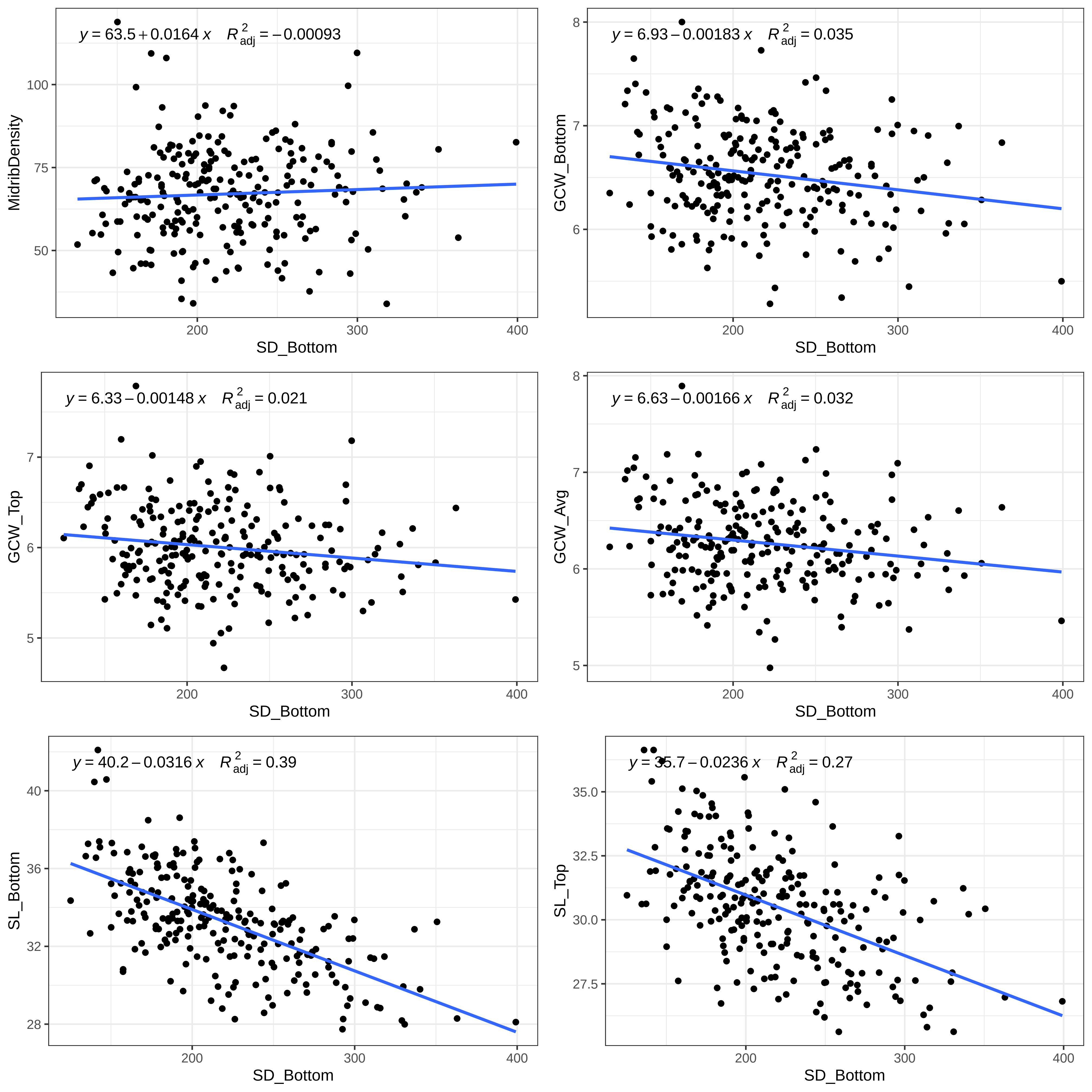

y = 63.5 + 0.0164 x R²adj = −0.00093
MidribDensity
SD_Bottom
y = 6.93 − 0.00183 x R²adj = 0.035
GCW_Bottom
SD_Bottom
y = 6.33 − 0.00148 x R²adj = 0.021
GCW_Top
SD_Bottom
y = 6.63 − 0.00166 x R²adj = 0.032
GCW_Avg
SD_Bottom
y = 40.2 − 0.0316 x R²adj = 0.39
SL_Bottom
SD_Bottom
y = 35.7 − 0.0236 x R²adj = 0.27
SL_Top
SD_Bottom

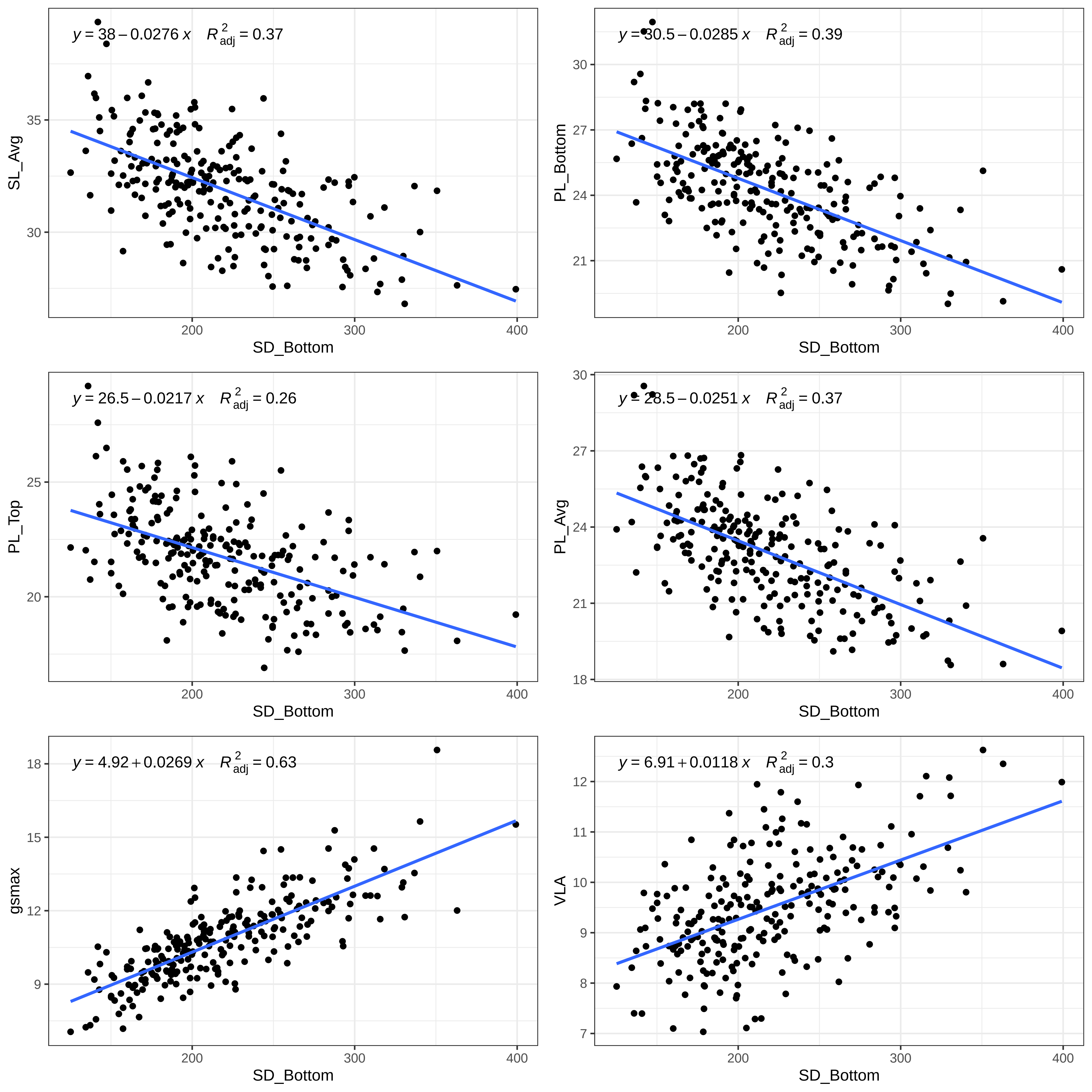

y = 38 − 0.0276 x   R²adj = 0.37
SL_Avg
SD_Bottom
y = 30.5 − 0.0285 x   R²adj = 0.39
PL_Bottom
SD_Bottom
y = 26.5 − 0.0217 x   R²adj = 0.26
PL_Top
SD_Bottom
y = 28.5 − 0.0251 x   R²adj = 0.37
PL_Avg
SD_Bottom
y = 4.92 + 0.0269 x   R²adj = 0.63
gsmax
SD_Bottom
y = 6.91 + 0.0118 x   R²adj = 0.3
VLA
SD_Bottom

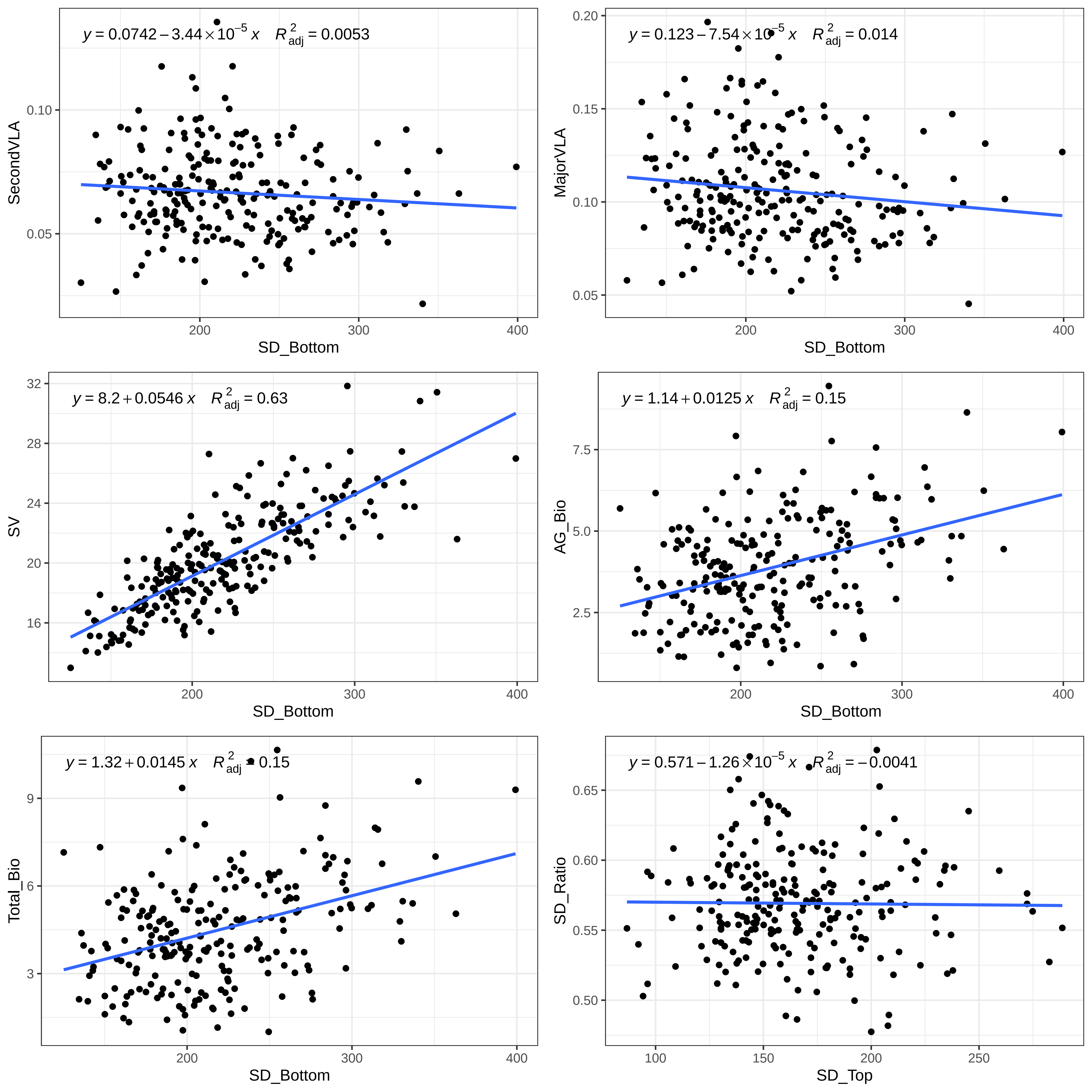

$y = 0.0742 - 3.44 \times 10^{-5}\,x \quad R^2_{adj} = 0.0053$
SecondVLA
SD_Bottom
$y = 0.123 - 7.54 \times 10^{-5}\,x \quad R^2_{adj} = 0.014$
MajorVLA
SD_Bottom
$y = 8.2 + 0.0546\,x \quad R^2_{adj} = 0.63$
SV
SD_Bottom
$y = 1.14 + 0.0125\,x \quad R^2_{adj} = 0.15$
AG_Bio
SD_Bottom
$y = 1.32 + 0.0145\,x \quad R^2_{adj} = 0.15$
Total_Bio
SD_Bottom
$y = 0.571 - 1.26 \times 10^{-5}\,x \quad R^2_{adj} = -0.0041$
SD_Ratio
SD_Top

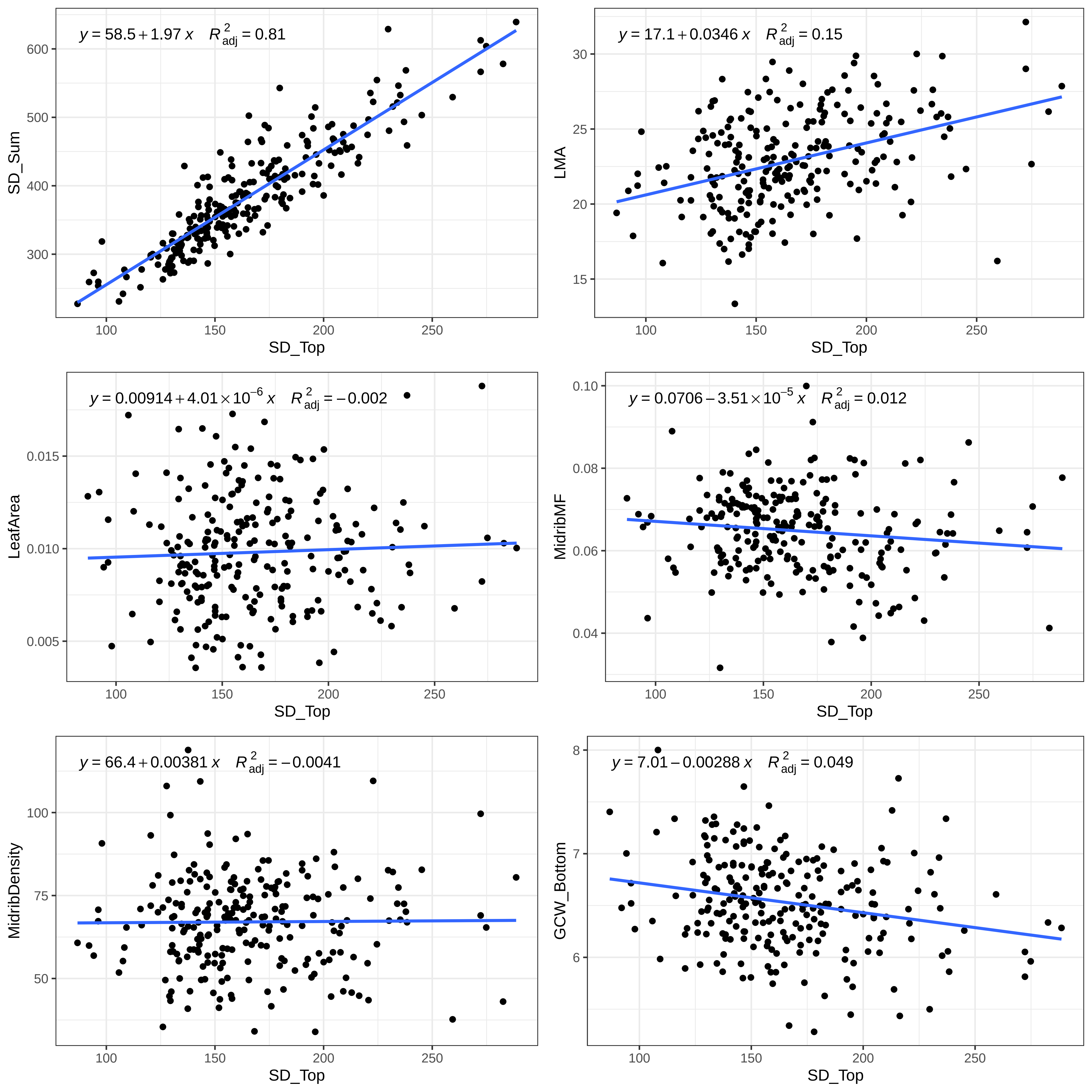
y = 58.5 + 1.97 x  R²adj = 0.81
SD_Sum
SD_Top
y = 17.1 + 0.0346 x  R²adj = 0.15
LMA
SD_Top
y = 0.00914 + 4.01 × 10⁻⁶ x  R²adj = −0.002
LeafArea
SD_Top
y = 0.0706 − 3.51 × 10⁻⁵ x  R²adj = 0.012
MidribMF
SD_Top
y = 66.4 + 0.00381 x  R²adj = −0.0041
MidribDensity
SD_Top
y = 7.01 − 0.00288 x  R²adj = 0.049
GCW_Bottom
SD_Top

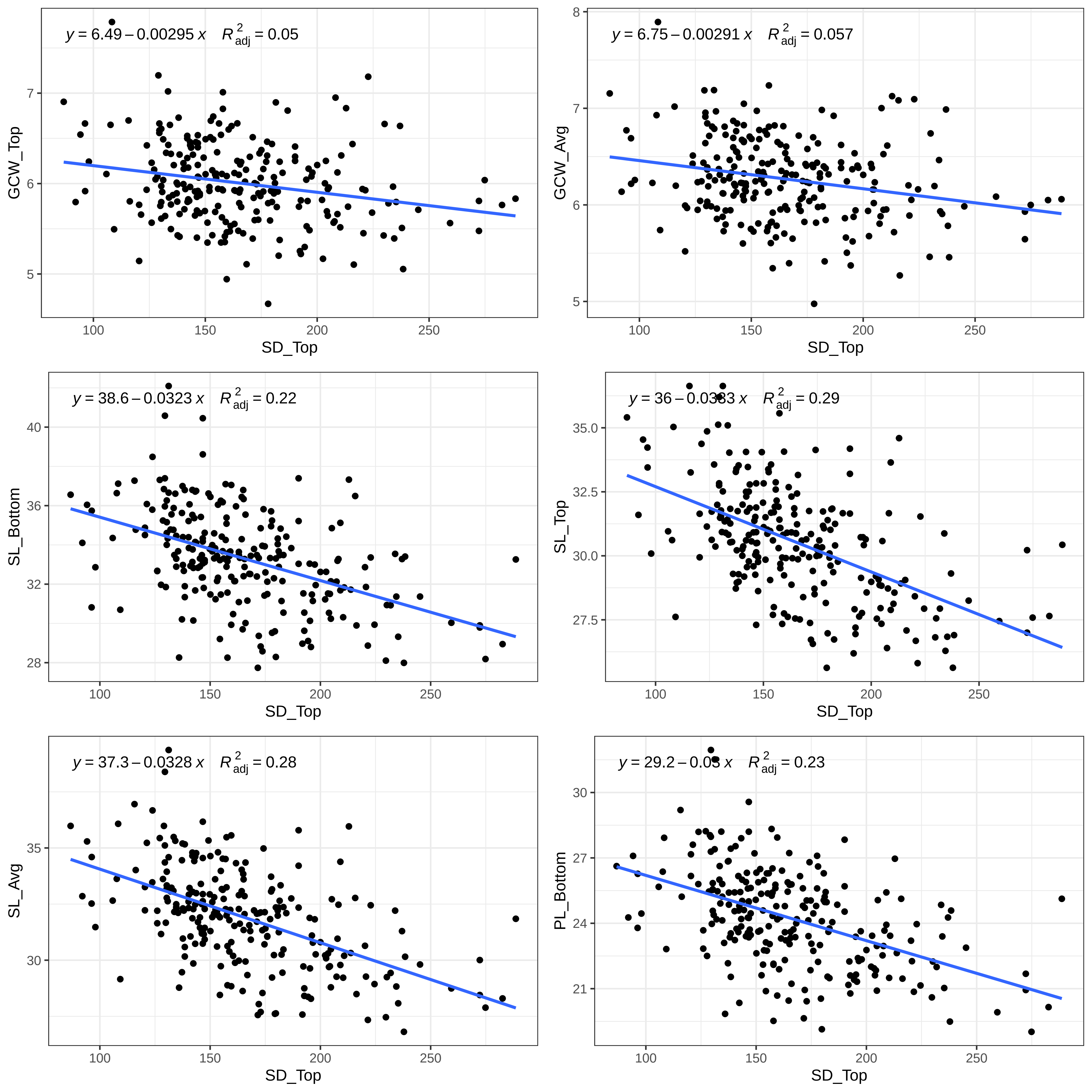
y = 6.49 – 0.00295 x R²adj = 0.05
GCW_Top
SD_Top
y = 6.75 – 0.00291 x R²adj = 0.057
GCW_Avg
SD_Top
y = 38.6 – 0.0323 x R²adj = 0.22
SL_Bottom
SD_Top
y = 36 – 0.0333 x R²adj = 0.29
SL_Top
SD_Top
y = 37.3 – 0.0328 x R²adj = 0.28
SL_Avg
SD_Top
y = 29.2 – 0.03 x R²adj = 0.23
PL_Bottom
SD_Top

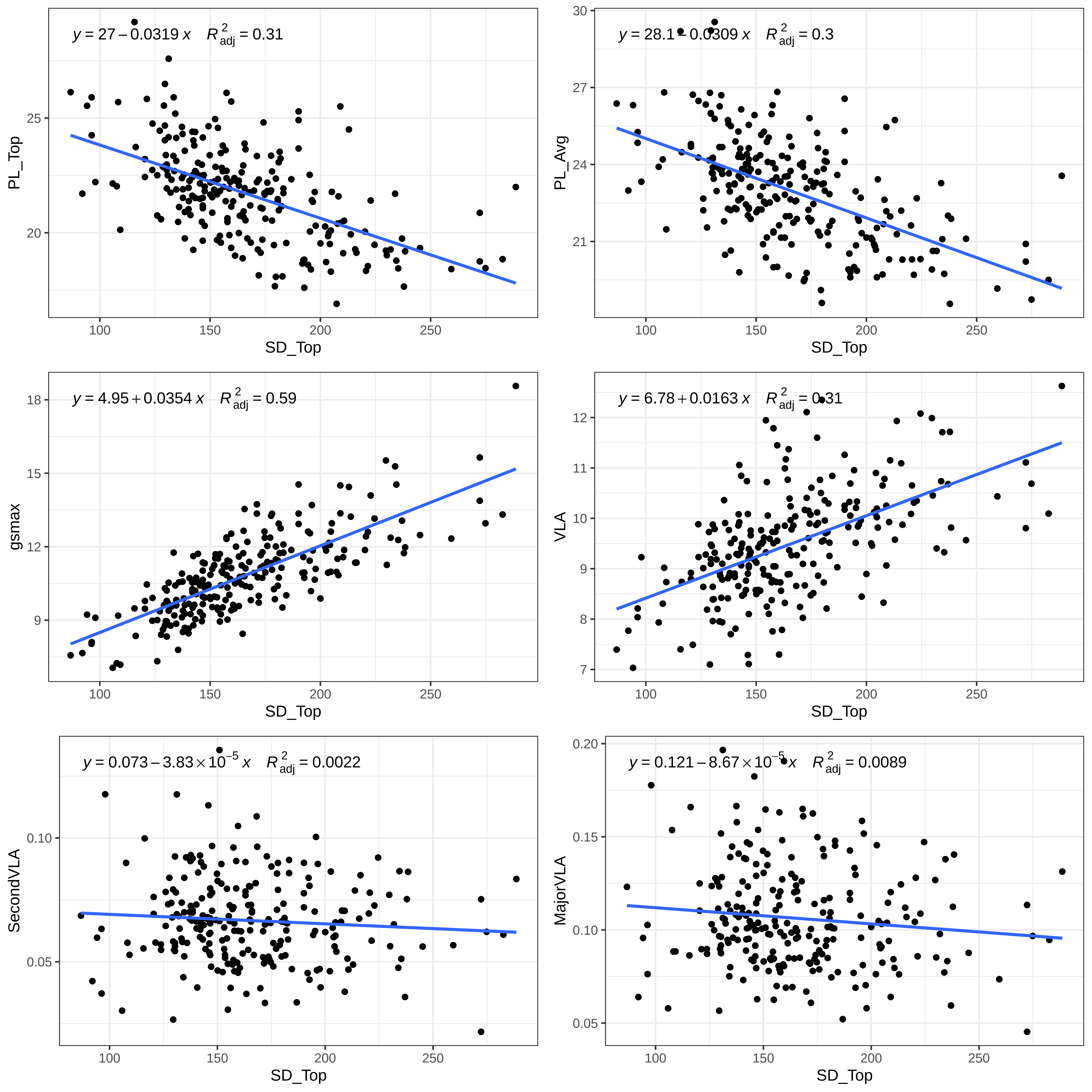
y = 27 − 0.0319 x R²adj = 0.31
PL_Top
SD_Top
y = 28.1 − 0.0309 x R²adj = 0.3
PL_Avg
SD_Top
y = 4.95 + 0.0354 x R²adj = 0.59
gsmax
SD_Top
y = 6.78 + 0.0163 x R²adj = 0.31
VLA
SD_Top
y = 0.073 − 3.83 × 10⁻⁵ x R²adj = 0.0022
SecondVLA
SD_Top
y = 0.121 − 8.67 × 10⁻⁵ x R²adj = 0.0089
MajorVLA
SD_Top

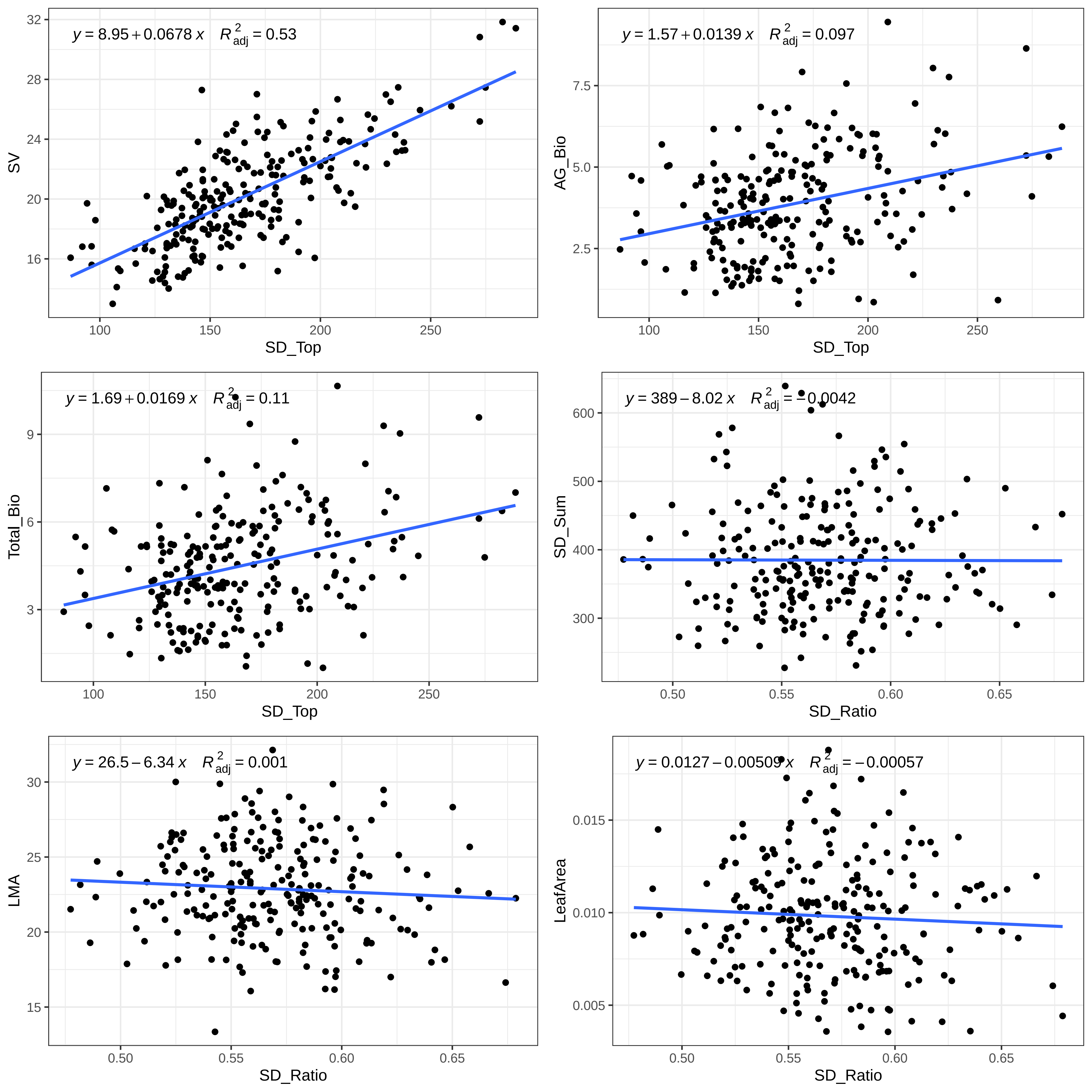

$y = 8.95 + 0.0678\,x \quad R^2_{adj} = 0.53$
SV
SD_Top
$y = 1.57 + 0.0139\,x \quad R^2_{adj} = 0.097$
AG_Bio
SD_Top
$y = 1.69 + 0.0169\,x \quad R^2_{adj} = 0.11$
Total_Bio
SD_Top
$y = 389 - 8.02\,x \quad R^2_{adj} = -0.0042$
SD_Sum
SD_Ratio
$y = 26.5 - 6.34\,x \quad R^2_{adj} = 0.001$
LMA
SD_Ratio
$y = 0.0127 - 0.00509\,x \quad R^2_{adj} = -0.00057$
LeafArea
SD_Ratio

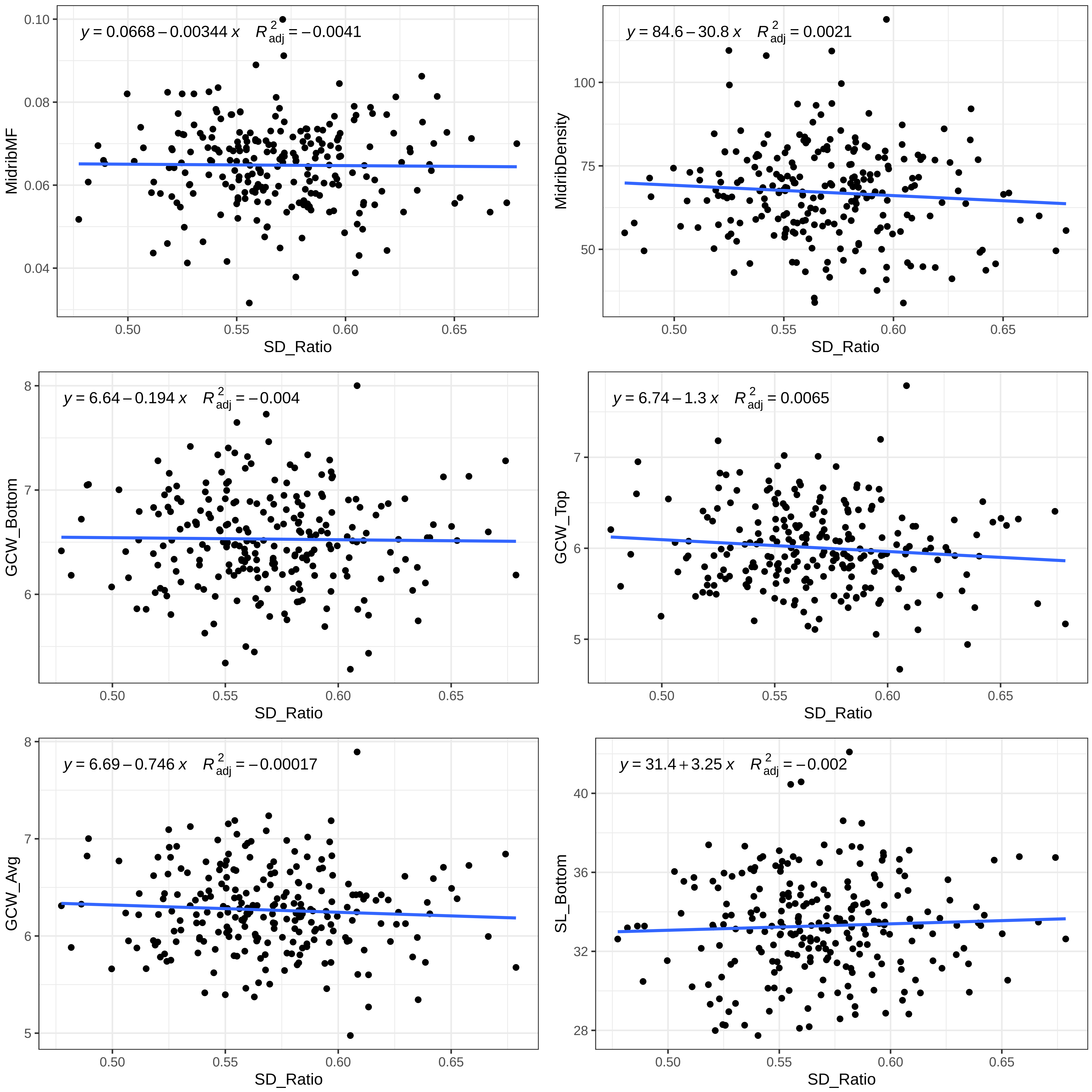

y = 0.0668 – 0.00344 x R²adj = –0.0041
MidribMF
SD_Ratio
y = 84.6 – 30.8 x R²adj = 0.0021
MidribDensity
SD_Ratio
y = 6.64 – 0.194 x R²adj = –0.004
GCW_Bottom
SD_Ratio
y = 6.74 – 1.3 x R²adj = 0.0065
GCW_Top
SD_Ratio
y = 6.69 – 0.746 x R²adj = –0.00017
GCW_Avg
SD_Ratio
y = 31.4 + 3.25 x R²adj = –0.002
SL_Bottom
SD_Ratio

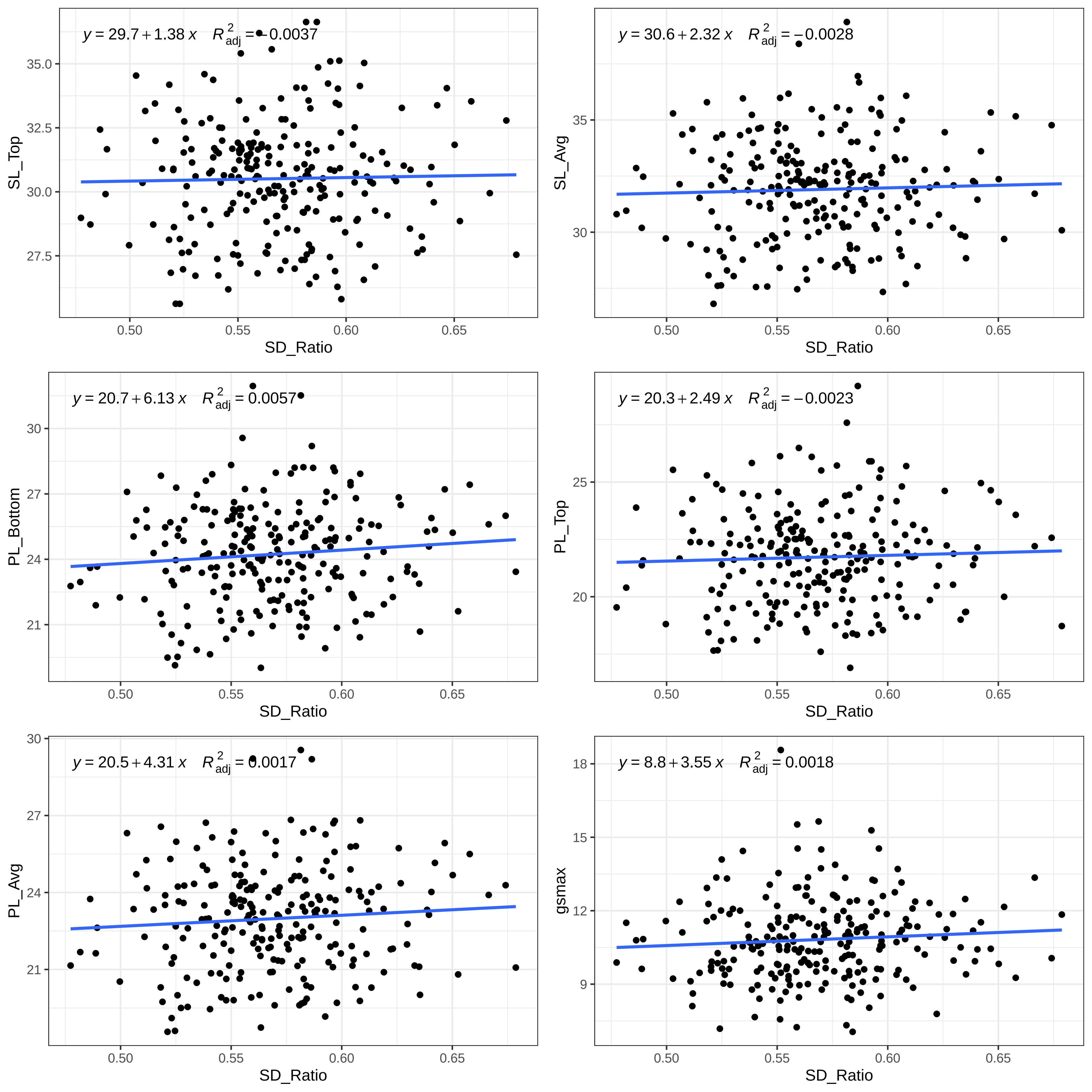

y = 29.7 + 1.38 x R²adj = −0.0037
SL_Top
SD_Ratio
y = 30.6 + 2.32 x R²adj = −0.0028
SL_Avg
SD_Ratio
y = 20.7 + 6.13 x R²adj = 0.0057
PL_Bottom
SD_Ratio
y = 20.3 + 2.49 x R²adj = −0.0023
PL_Top
SD_Ratio
y = 20.5 + 4.31 x R²adj = 0.0017
PL_Avg
SD_Ratio
y = 8.8 + 3.55 x R²adj = 0.0018
gsmax
SD_Ratio

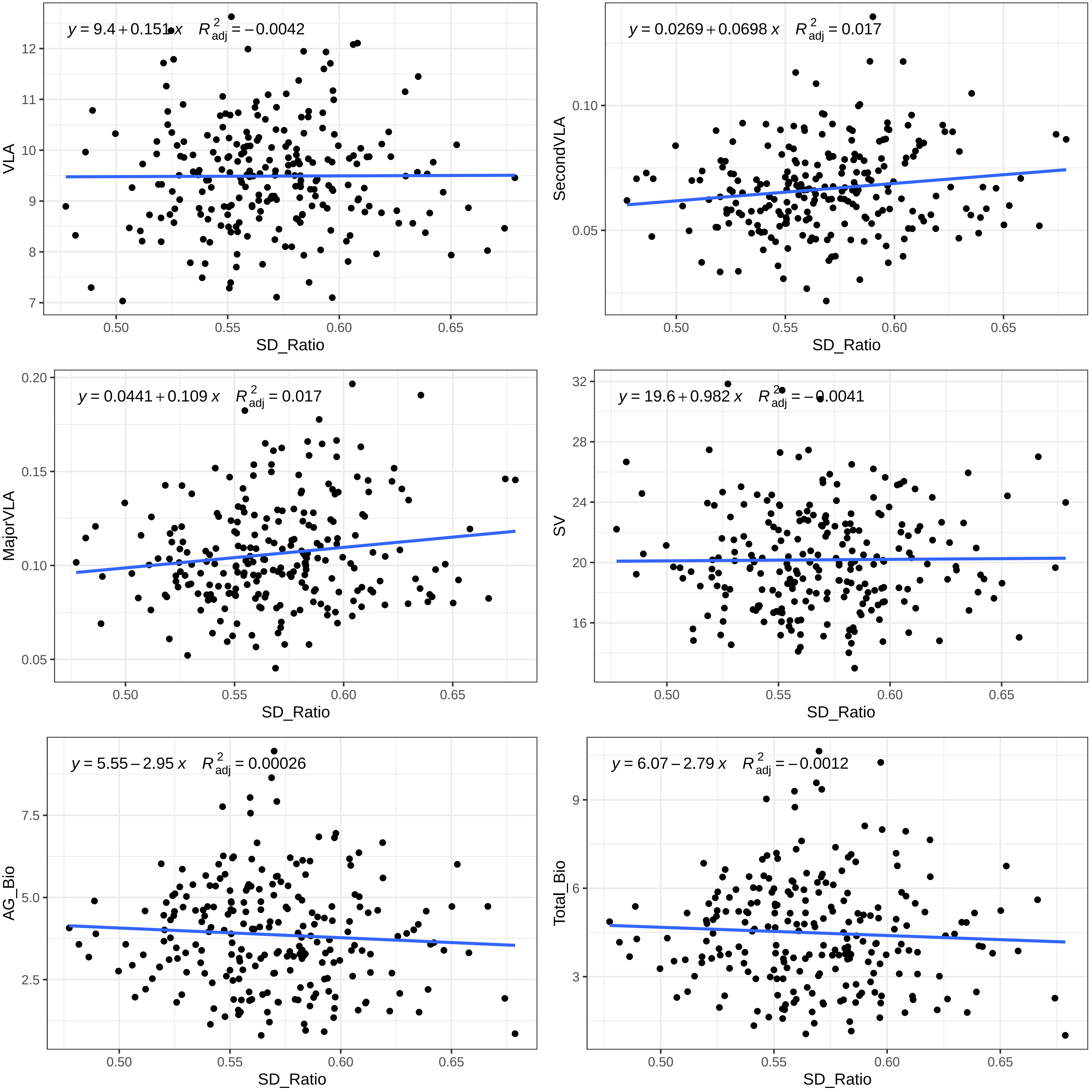

y = 9.4 + 0.151 x    R²adj = −0.0042
VLA
SD_Ratio
y = 0.0269 + 0.0698 x    R²adj = 0.017
SecondVLA
SD_Ratio
y = 0.0441 + 0.109 x    R²adj = 0.017
MajorVLA
SD_Ratio
y = 19.6 + 0.982 x    R²adj = −0.0041
SV
SD_Ratio
y = 5.55 − 2.95 x    R²adj = 0.00026
AG_Bio
SD_Ratio
y = 6.07 − 2.79 x    R²adj = −0.0012
Total_Bio
SD_Ratio

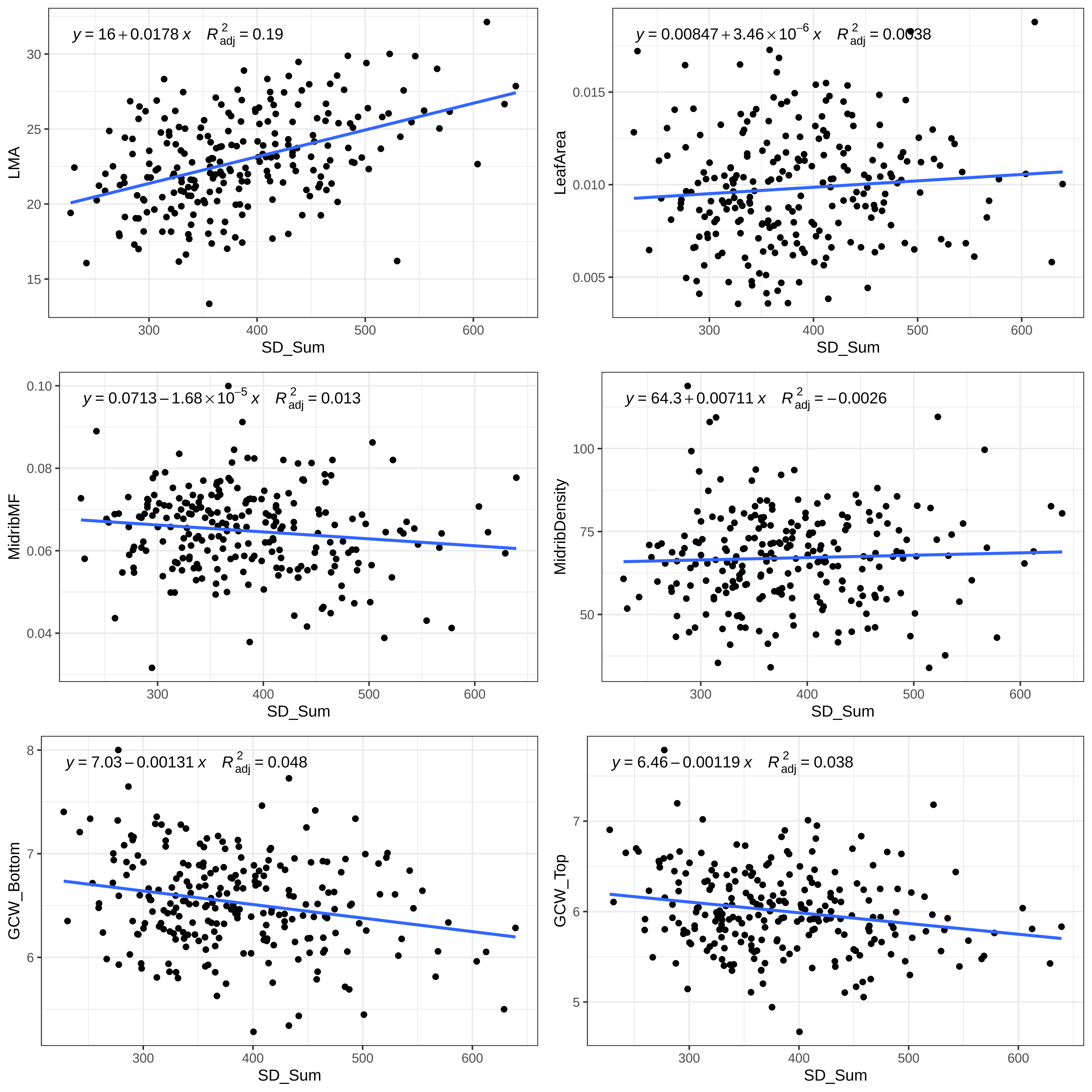
$y = 16 + 0.0178\,x \quad R^2_{adj} = 0.19$
LMA
SD_Sum
$y = 0.00847 + 3.46 \times 10^{-6}\,x \quad R^2_{adj} = 0.0038$
LeafArea
SD_Sum
$y = 0.0713 - 1.68 \times 10^{-5}\,x \quad R^2_{adj} = 0.013$
MidribMF
SD_Sum
$y = 64.3 + 0.00711\,x \quad R^2_{adj} = -0.0026$
MidribDensity
SD_Sum
$y = 7.03 - 0.00131\,x \quad R^2_{adj} = 0.048$
GCW_Bottom
SD_Sum
$y = 6.46 - 0.00119\,x \quad R^2_{adj} = 0.038$
GCW_Top
SD_Sum

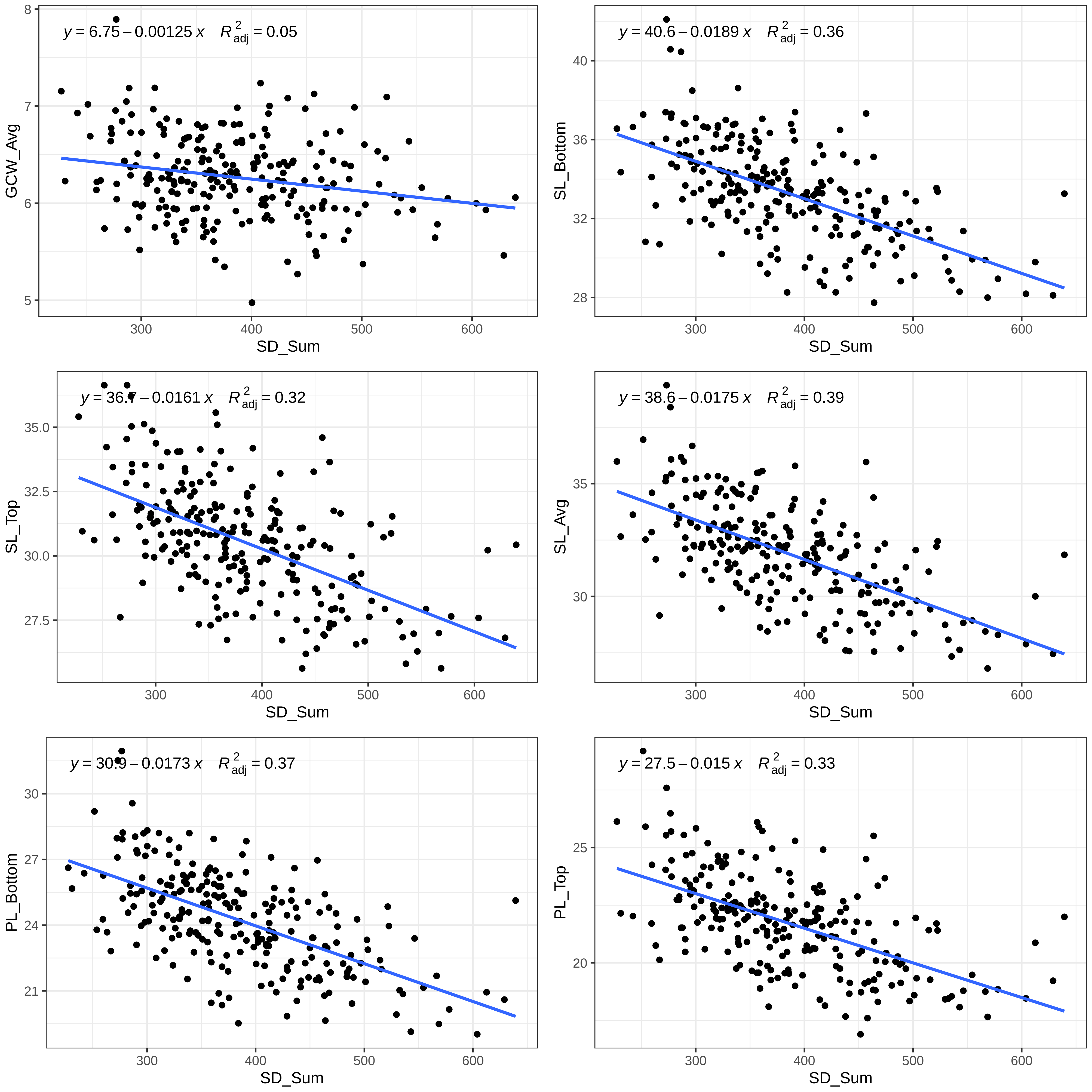
y = 6.75 – 0.00125 x   R²adj = 0.05
GCW_Avg
SD_Sum
y = 40.6 – 0.0189 x   R²adj = 0.36
SL_Bottom
SD_Sum
y = 36.7 – 0.0161 x   R²adj = 0.32
SL_Top
SD_Sum
y = 38.6 – 0.0175 x   R²adj = 0.39
SL_Avg
SD_Sum
y = 30.9 – 0.0173 x   R²adj = 0.37
PL_Bottom
SD_Sum
y = 27.5 – 0.015 x   R²adj = 0.33
PL_Top
SD_Sum

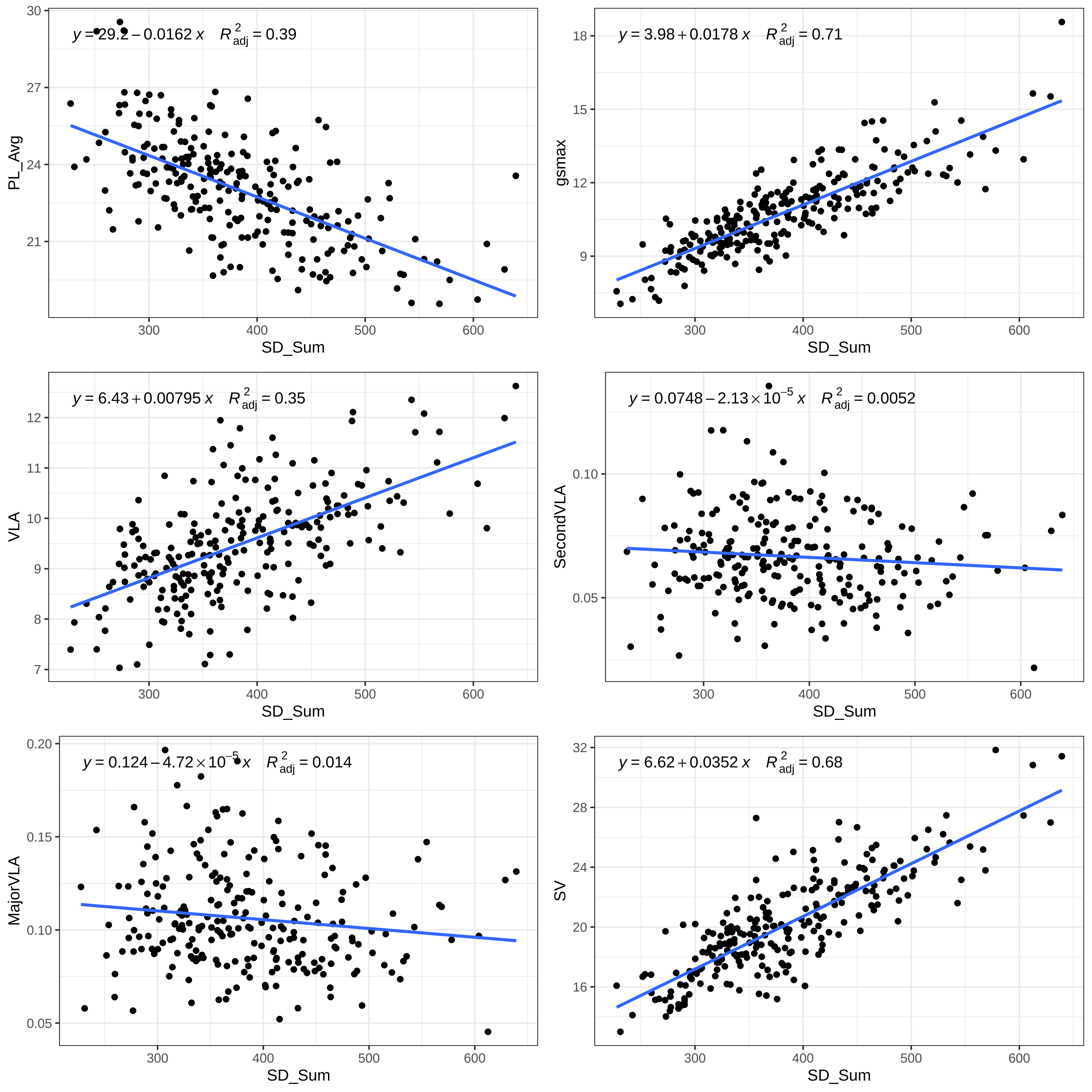

y = 29.2 − 0.0162 x   R²adj = 0.39
PL_Avg
SD_Sum
y = 3.98 + 0.0178 x   R²adj = 0.71
gsmax
SD_Sum
y = 6.43 + 0.00795 x   R²adj = 0.35
VLA
SD_Sum
y = 0.0748 − 2.13 × 10⁻⁵ x   R²adj = 0.0052
SecondVLA
SD_Sum
y = 0.124 − 4.72 × 10⁻⁵ x   R²adj = 0.014
MajorVLA
SD_Sum
y = 6.62 + 0.0352 x   R²adj = 0.68
SV
SD_Sum

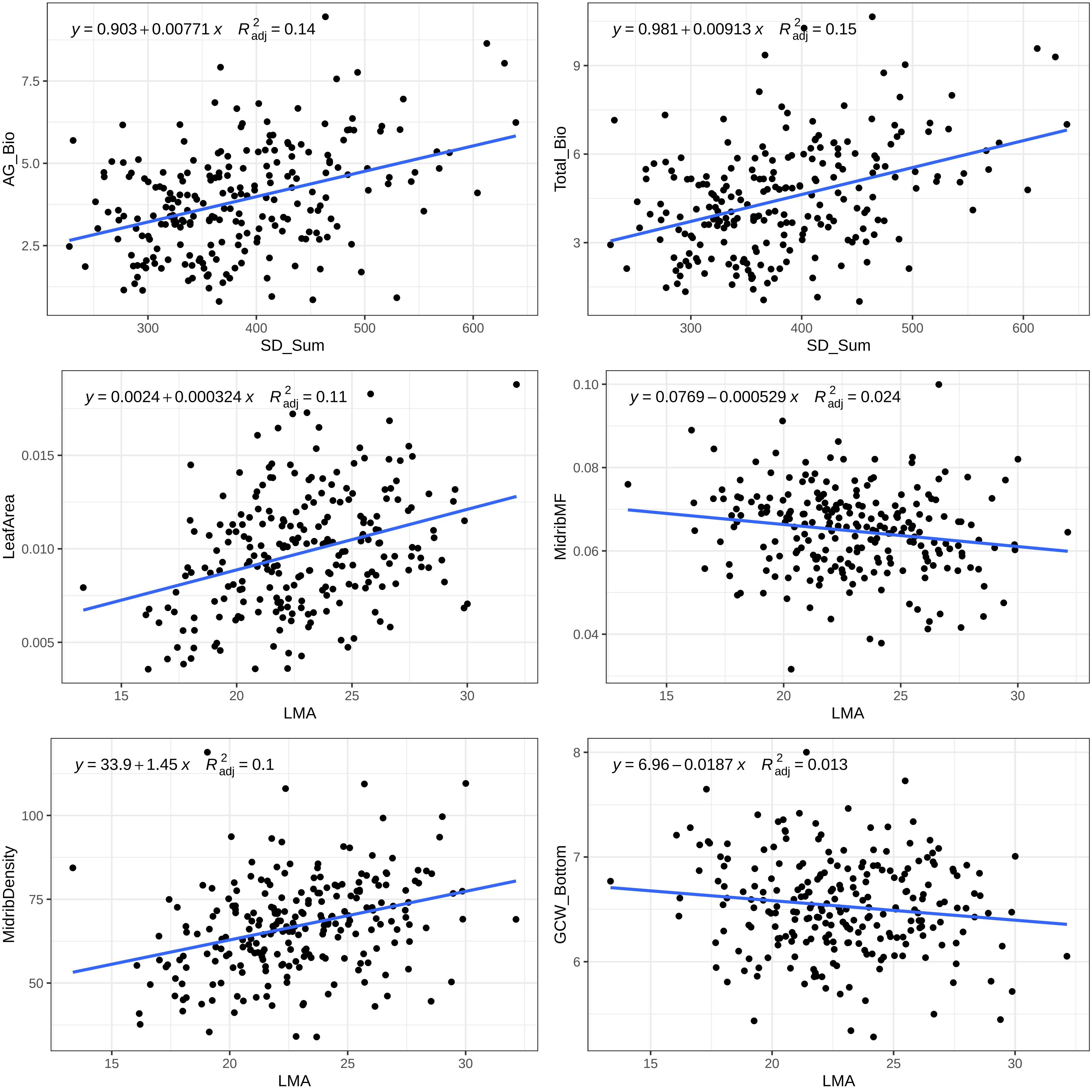

$y = 0.903 + 0.00771\,x \quad R^2_{adj} = 0.14$
AG_Bio
SD_Sum
$y = 0.981 + 0.00913\,x \quad R^2_{adj} = 0.15$
Total_Bio
SD_Sum
$y = 0.0024 + 0.000324\,x \quad R^2_{adj} = 0.11$
LeafArea
LMA
$y = 0.0769 - 0.000529\,x \quad R^2_{adj} = 0.024$
MidribMF
LMA
$y = 33.9 + 1.45\,x \quad R^2_{adj} = 0.1$
MidribDensity
LMA
$y = 6.96 - 0.0187\,x \quad R^2_{adj} = 0.013$
GCW_Bottom
LMA

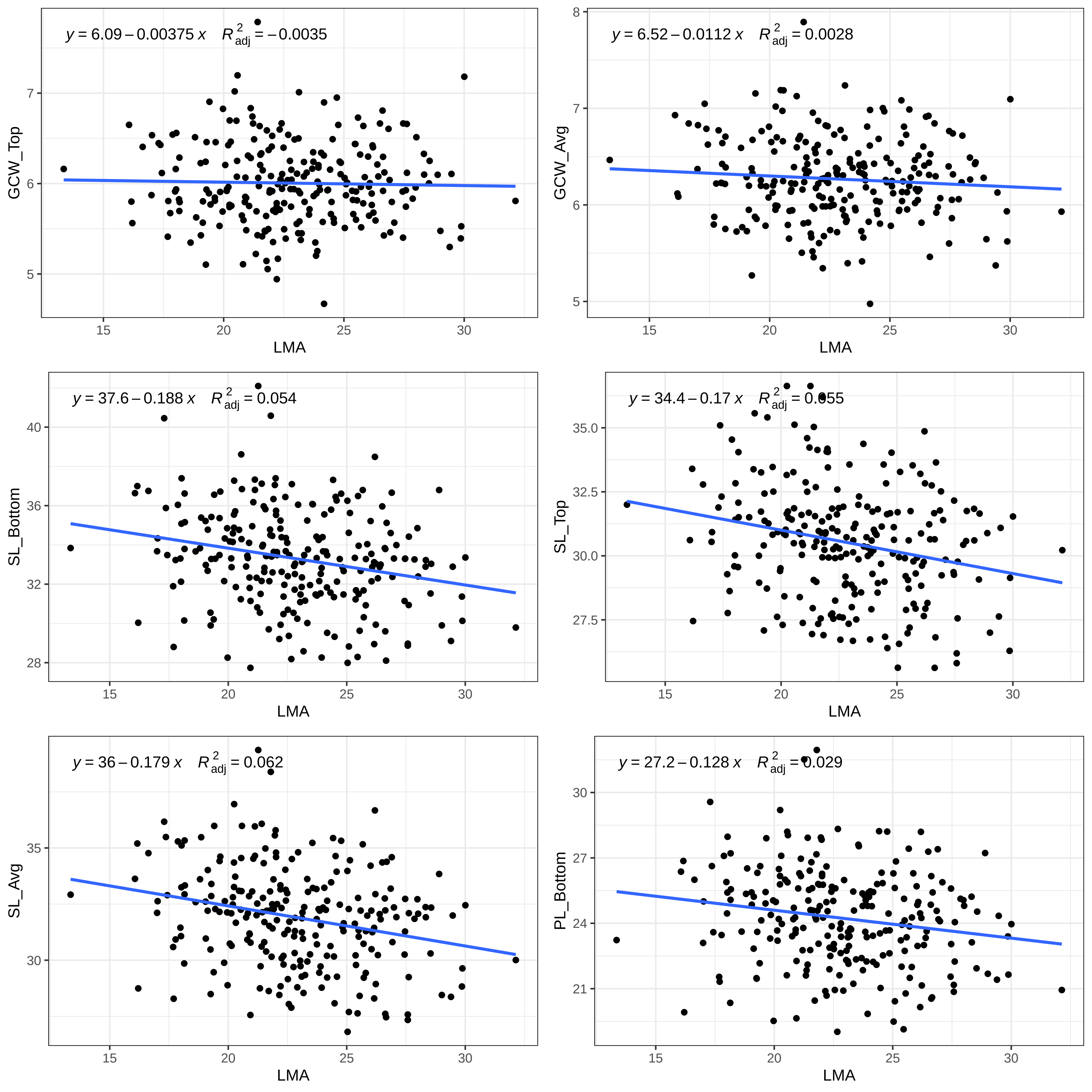

y = 6.09 – 0.00375 x  R²adj = – 0.0035
GCW_Top
LMA
y = 6.52 – 0.0112 x  R²adj = 0.0028
GCW_Avg
LMA
y = 37.6 – 0.188 x  R²adj = 0.054
SL_Bottom
LMA
y = 34.4 – 0.17 x  R²adj = 0.055
SL_Top
LMA
y = 36 – 0.179 x  R²adj = 0.062
SL_Avg
LMA
y = 27.2 – 0.128 x  R²adj = 0.029
PL_Bottom
LMA

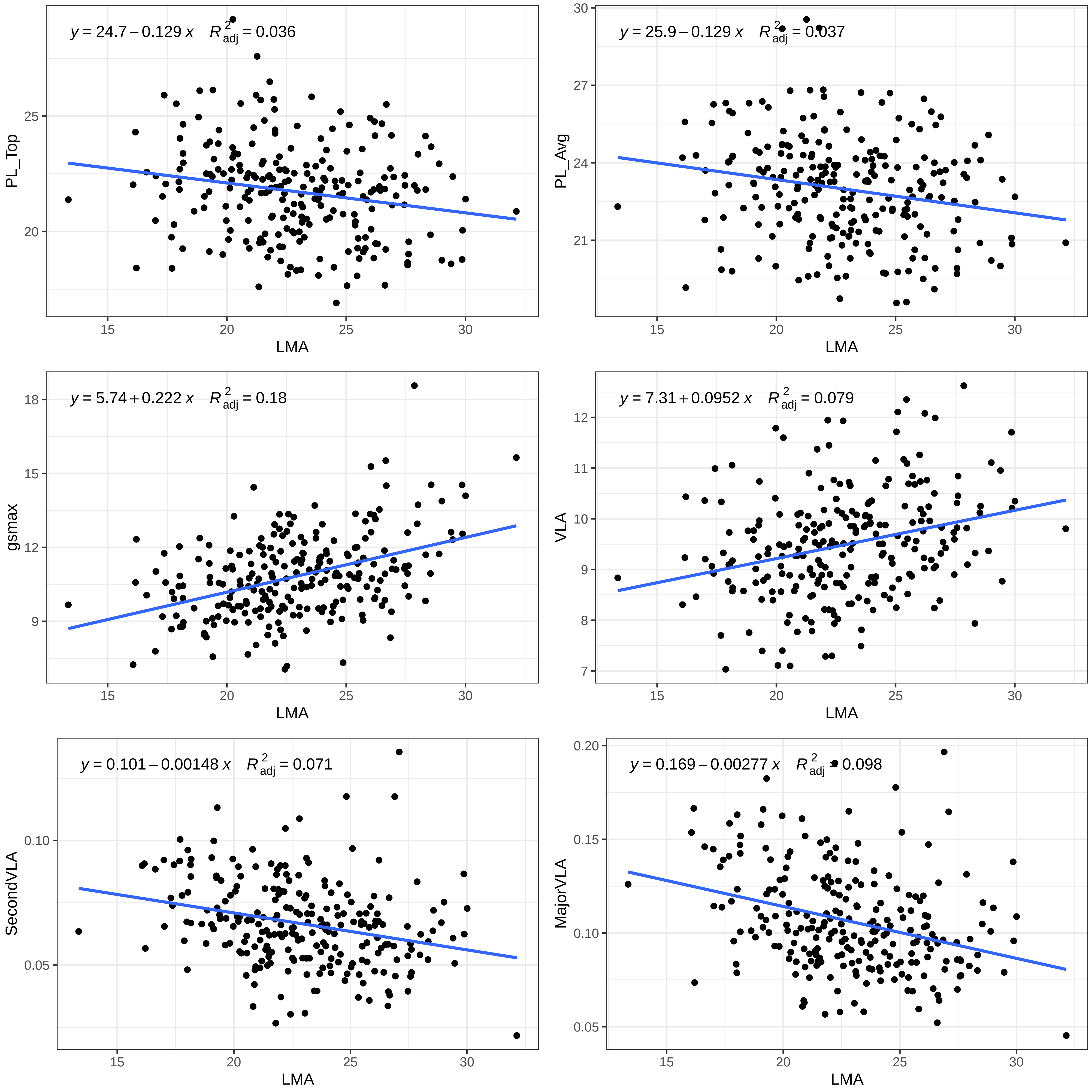

y = 24.7 − 0.129 x R2adj = 0.036
PL_Top
LMA
y = 25.9 − 0.129 x R2adj = 0.037
PL_Avg
LMA
y = 5.74 + 0.222 x R2adj = 0.18
gsmax
LMA
y = 7.31 + 0.0952 x R2adj = 0.079
VLA
LMA
y = 0.101 − 0.00148 x R2adj = 0.071
SecondVLA
LMA
y = 0.169 − 0.00277 x R2adj = 0.098
MajorVLA
LMA

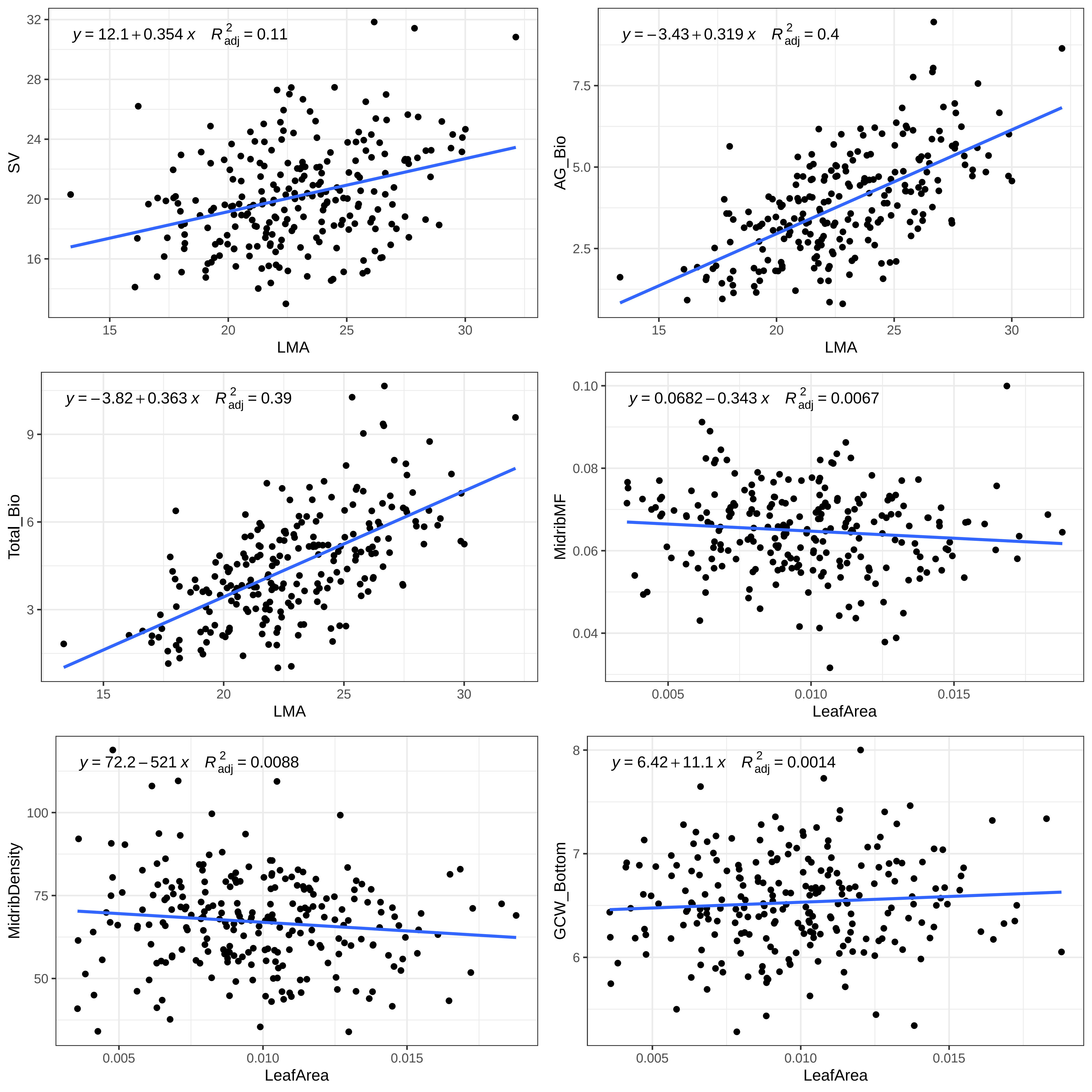

y = 12.1 + 0.354 x  R²adj = 0.11
SV
LMA
y = −3.43 + 0.319 x  R²adj = 0.4
AG_Bio
LMA
y = −3.82 + 0.363 x  R²adj = 0.39
Total_Bio
LMA
y = 0.0682 − 0.343 x  R²adj = 0.0067
MidribMF
LeafArea
y = 72.2 − 521 x  R²adj = 0.0088
MidribDensity
LeafArea
y = 6.42 + 11.1 x  R²adj = 0.0014
GCW_Bottom
LeafArea

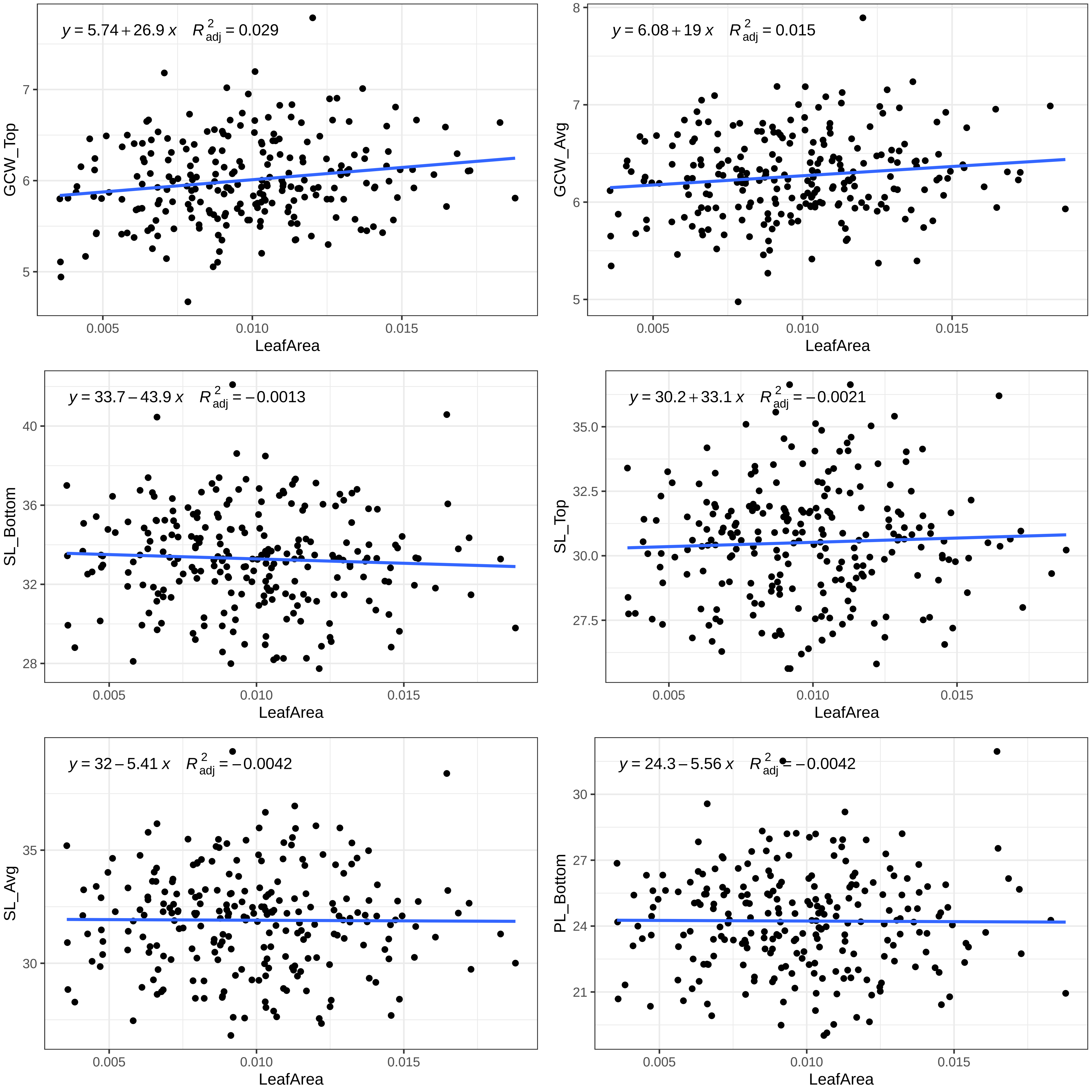

y = 5.74 + 26.9 x    R²adj = 0.029
GCW_Top
7
6
5
0.005
0.010
0.015
LeafArea
y = 6.08 + 19 x    R²adj = 0.015
GCW_Avg
8
7
6
5
0.005
0.010
0.015
LeafArea
y = 33.7 − 43.9 x    R²adj = −0.0013
SL_Bottom
40
36
32
28
0.005
0.010
0.015
LeafArea
y = 30.2 + 33.1 x    R²adj = −0.0021
SL_Top
35.0
32.5
30.0
27.5
0.005
0.010
0.015
LeafArea
y = 32 − 5.41 x    R²adj = −0.0042
SL_Avg
35
30
0.005
0.010
0.015
LeafArea
y = 24.3 − 5.56 x    R²adj = −0.0042
PL_Bottom
30
27
24
21
0.005
0.010
0.015
LeafArea

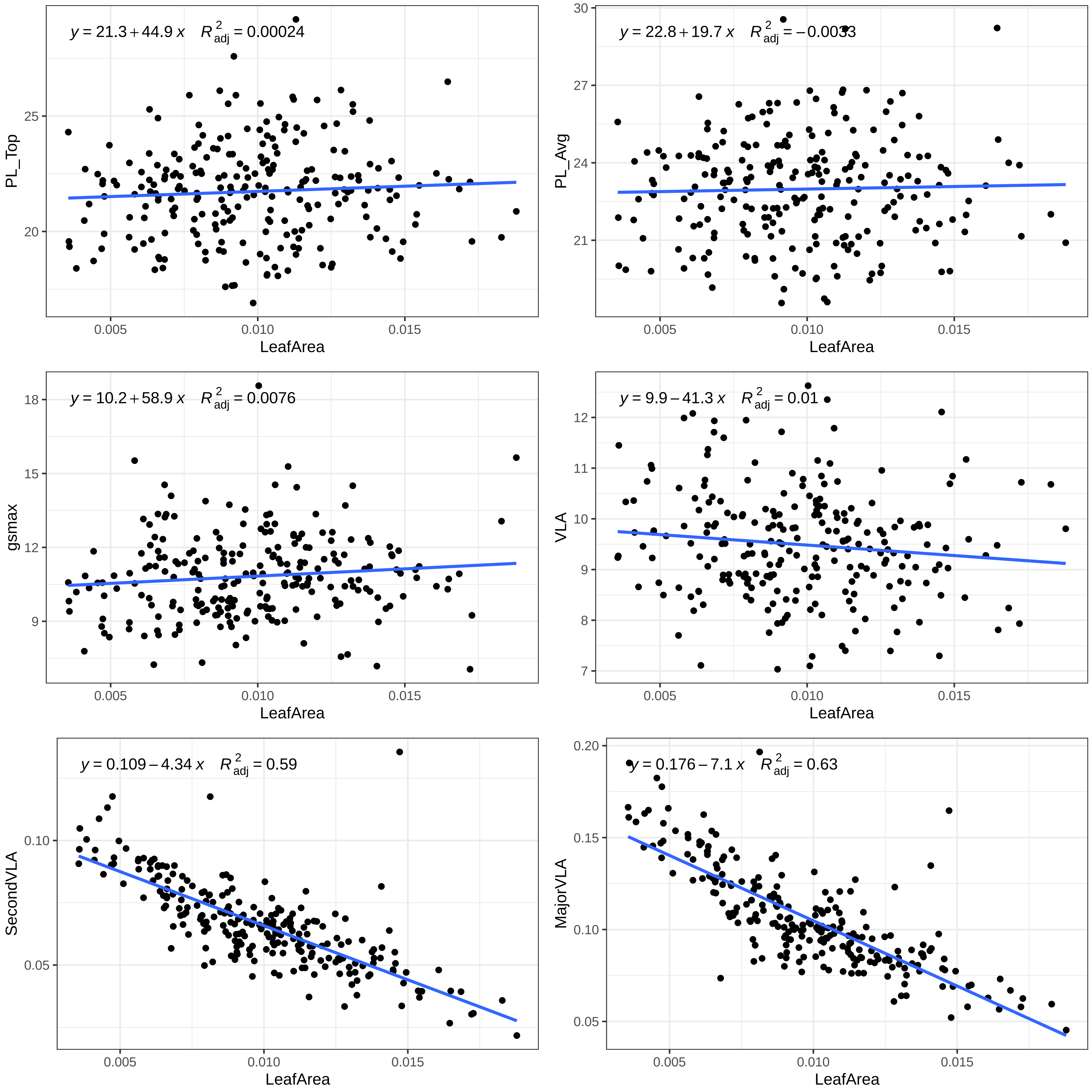

y = 21.3 + 44.9 x R²adj = 0.00024
PL_Top
LeafArea
y = 22.8 + 19.7 x R²adj = −0.0033
PL_Avg
LeafArea
y = 10.2 + 58.9 x R²adj = 0.0076
gsmax
LeafArea
y = 9.9 − 41.3 x R²adj = 0.01
VLA
LeafArea
y = 0.109 − 4.34 x R²adj = 0.59
SecondVLA
LeafArea
y = 0.176 − 7.1 x R²adj = 0.63
MajorVLA
LeafArea

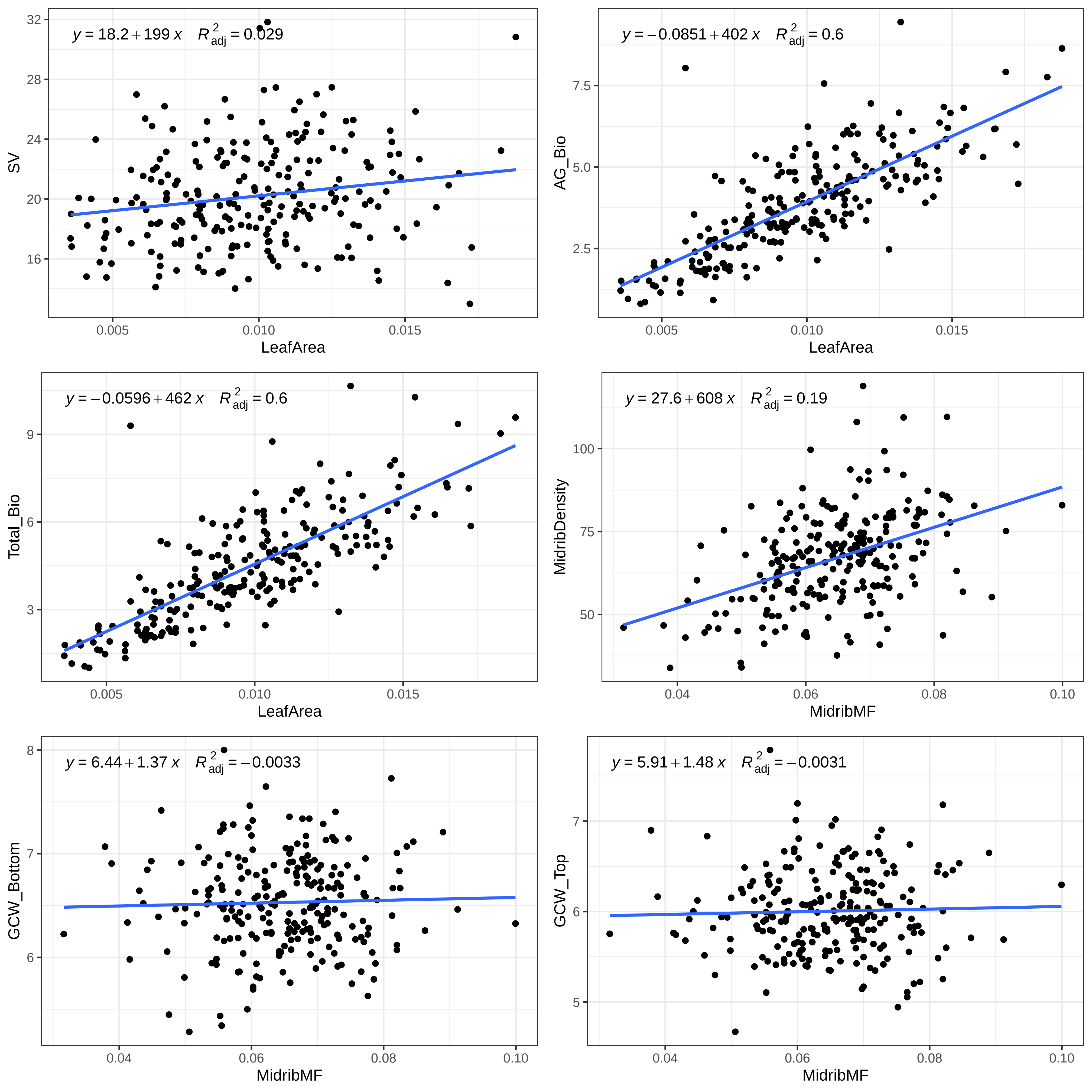

y = 18.2 + 199 x R²adj = 0.029
SV
32
28
24
20
16
0.005
0.010
0.015
LeafArea
y = −0.0851 + 402 x R²adj = 0.6
AG_Bio
7.5
5.0
2.5
0.005
0.010
0.015
LeafArea
y = −0.0596 + 462 x R²adj = 0.6
Total_Bio
9
6
3
0.005
0.010
0.015
LeafArea
y = 27.6 + 608 x R²adj = 0.19
MidribDensity
100
75
50
0.04
0.06
0.08
0.10
MidribMF
y = 6.44 + 1.37 x R²adj = −0.0033
GCW_Bottom
8
7
6
0.04
0.06
0.08
0.10
MidribMF
y = 5.91 + 1.48 x R²adj = −0.0031
GCW_Top
7
6
5
0.04
0.06
0.08
0.10
MidribMF

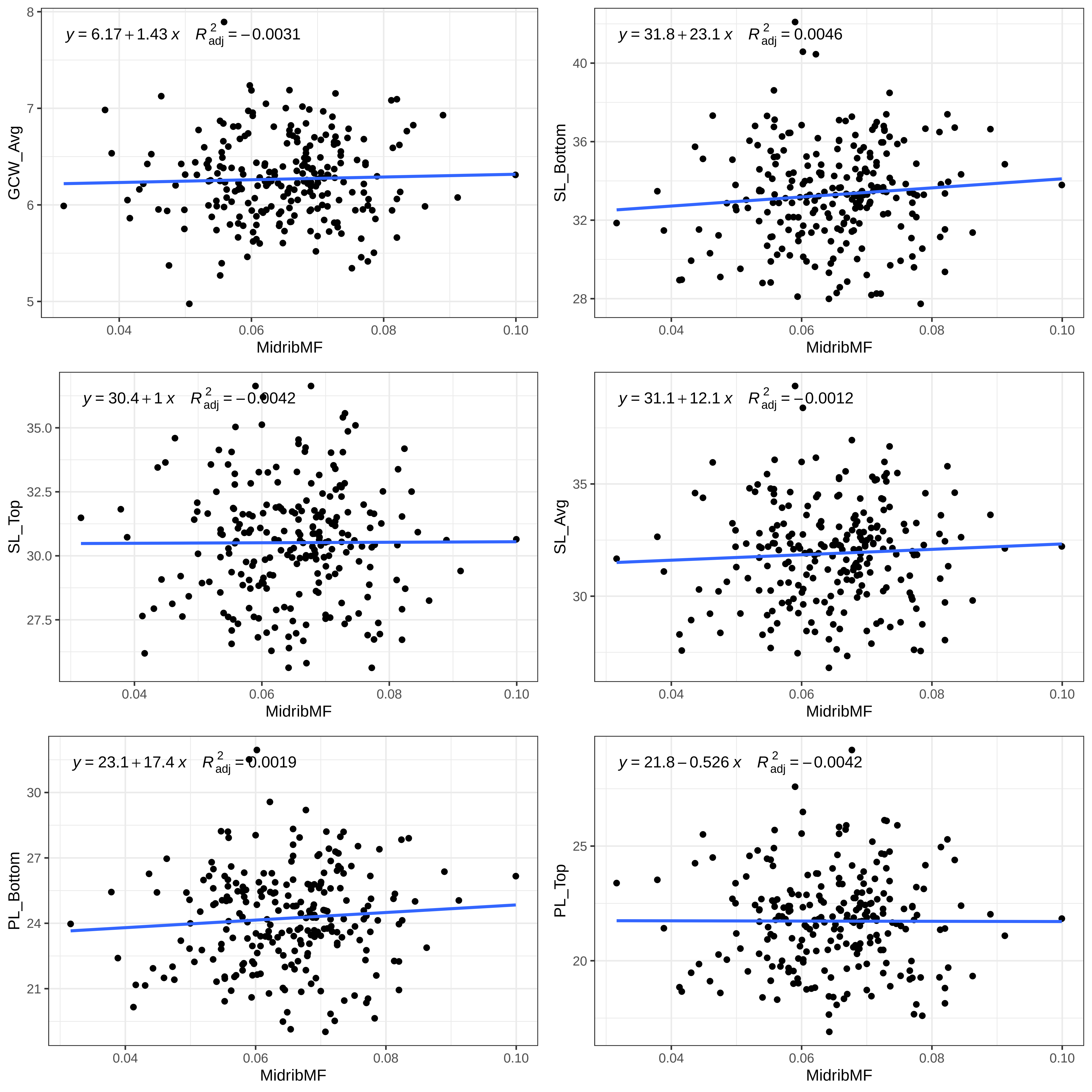

y = 6.17 + 1.43 x R2adj = −0.0031
GCW_Avg
MidribMF
y = 31.8 + 23.1 x R2adj = 0.0046
SL_Bottom
MidribMF
y = 30.4 + 1 x R2adj = −0.0042
SL_Top
MidribMF
y = 31.1 + 12.1 x R2adj = −0.0012
SL_Avg
MidribMF
y = 23.1 + 17.4 x R2adj = 0.0019
PL_Bottom
MidribMF
y = 21.8 − 0.526 x R2adj = −0.0042
PL_Top
MidribMF

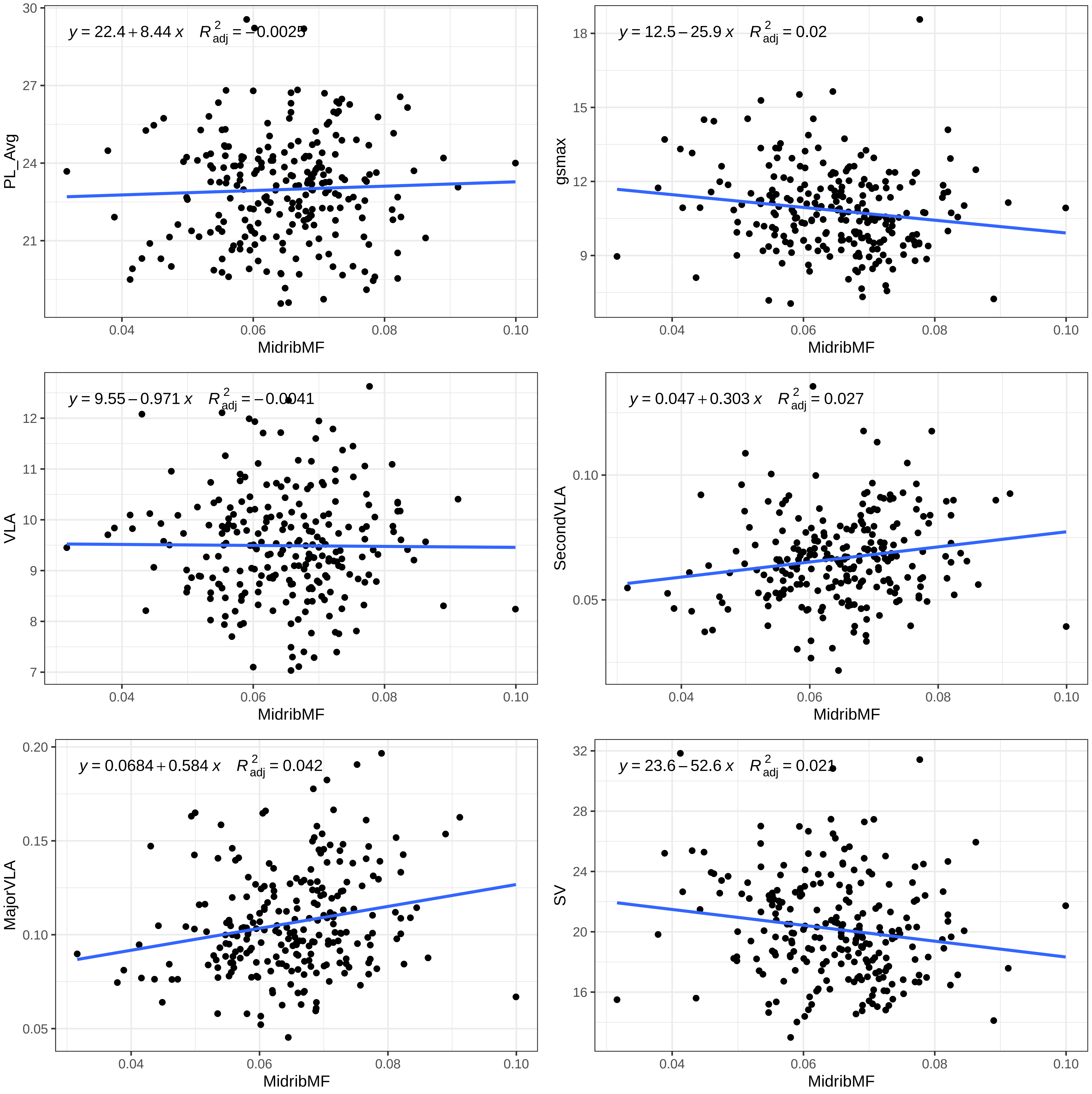

y = 22.4 + 8.44 x  R2adj = −0.0025
PL_Avg
21
24
27
30
MidribMF
0.04
0.06
0.08
0.10
y = 12.5 − 25.9 x  R2adj = 0.02
gsmax
9
12
15
18
MidribMF
0.04
0.06
0.08
0.10
y = 9.55 − 0.971 x  R2adj = −0.0041
VLA
7
8
9
10
11
12
MidribMF
0.04
0.06
0.08
0.10
y = 0.047 + 0.303 x  R2adj = 0.027
SecondVLA
0.05
0.10
MidribMF
0.04
0.06
0.08
0.10
y = 0.0684 + 0.584 x  R2adj = 0.042
MajorVLA
0.05
0.10
0.15
0.20
MidribMF
0.04
0.06
0.08
0.10
y = 23.6 − 52.6 x  R2adj = 0.021
SV
16
20
24
28
32
MidribMF
0.04
0.06
0.08
0.10

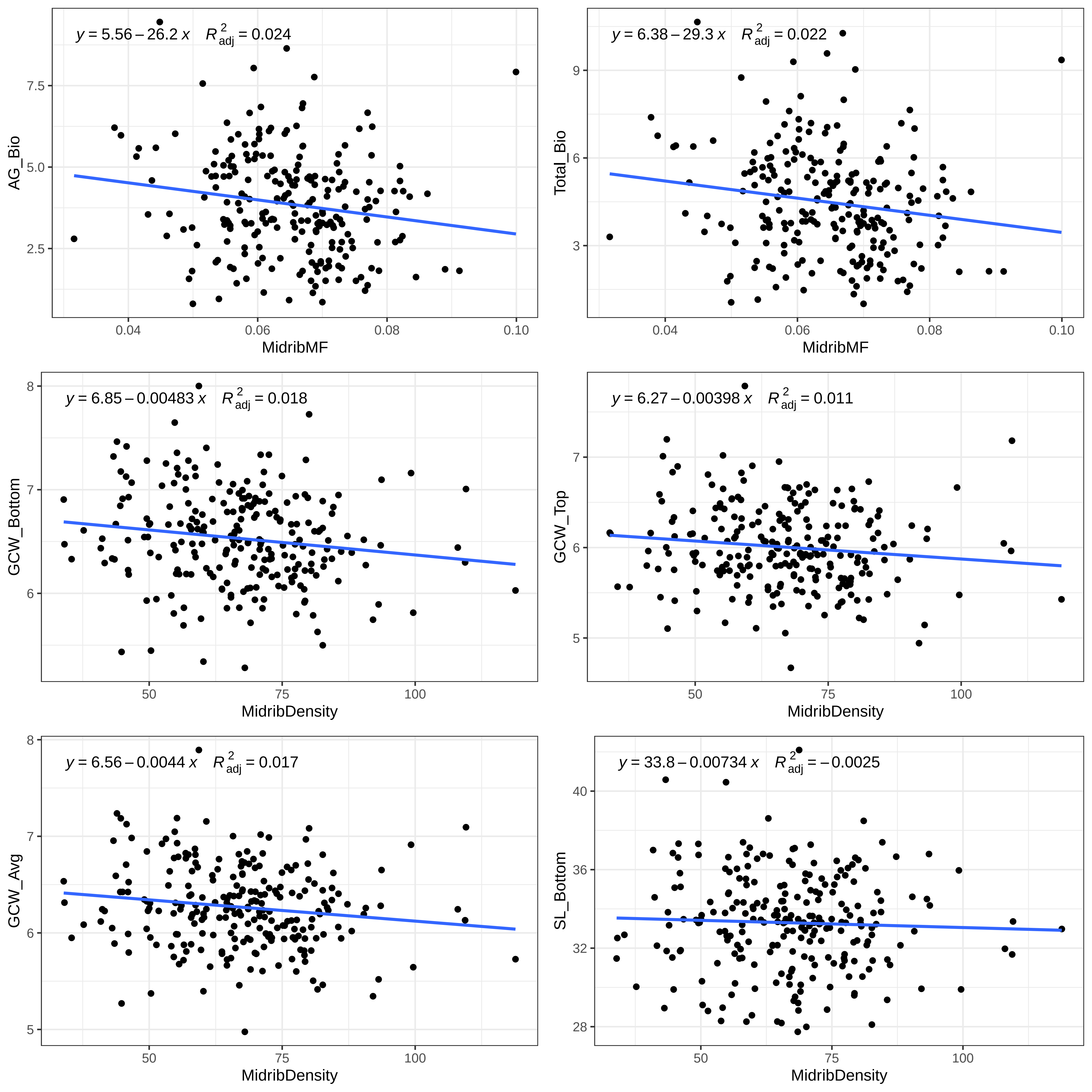

y = 5.56 – 26.2 x   R2adj = 0.024
AG_Bio
MidribMF
y = 6.38 – 29.3 x   R2adj = 0.022
Total_Bio
MidribMF
y = 6.85 – 0.00483 x   R2adj = 0.018
GCW_Bottom
MidribDensity
y = 6.27 – 0.00398 x   R2adj = 0.011
GCW_Top
MidribDensity
y = 6.56 – 0.0044 x   R2adj = 0.017
GCW_Avg
MidribDensity
y = 33.8 – 0.00734 x   R2adj = – 0.0025
SL_Bottom
MidribDensity

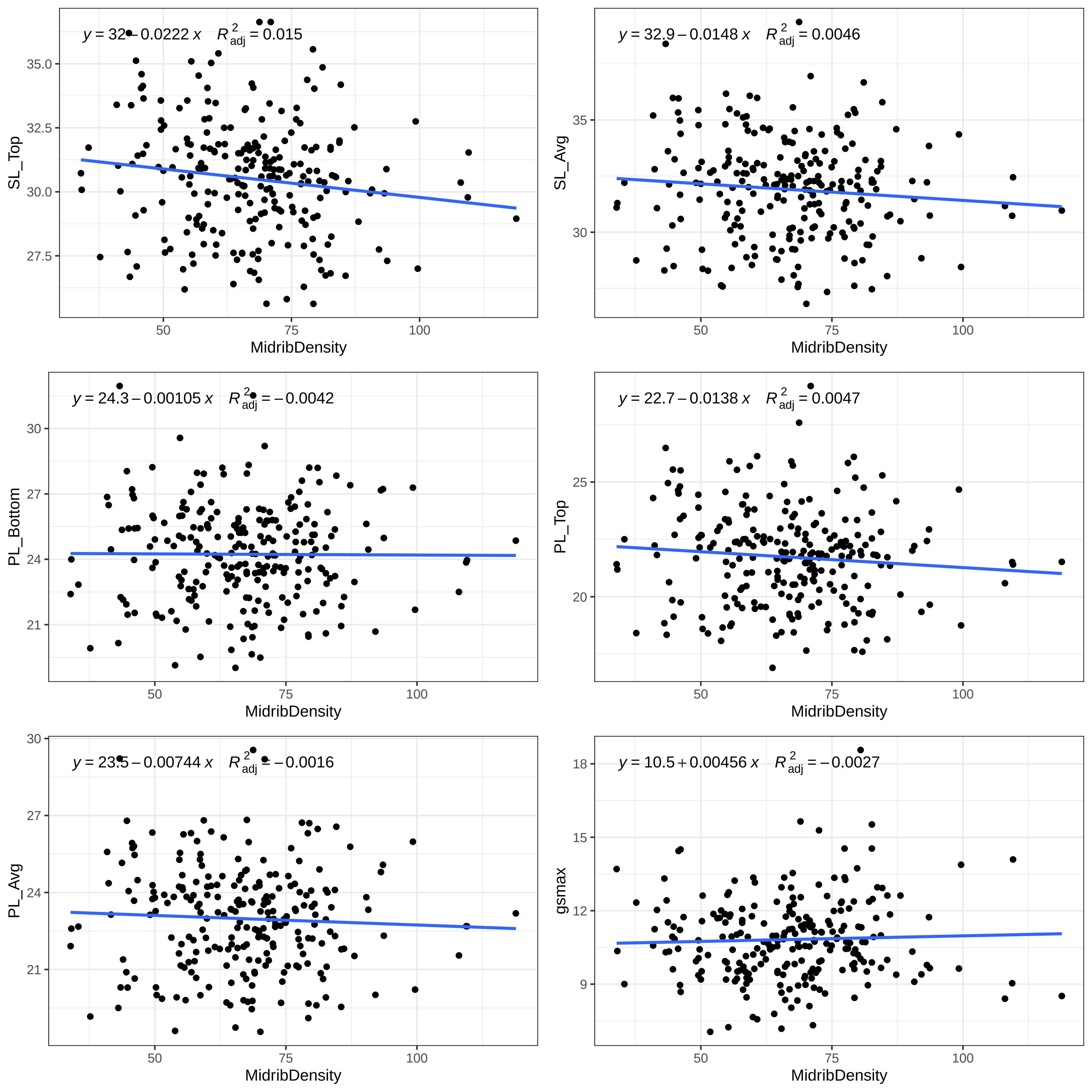
y = 32 − 0.0222 x  R²adj = 0.015
SL_Top
MidribDensity
y = 32.9 − 0.0148 x  R²adj = 0.0046
SL_Avg
MidribDensity
y = 24.3 − 0.00105 x  R²adj = −0.0042
PL_Bottom
MidribDensity
y = 22.7 − 0.0138 x  R²adj = 0.0047
PL_Top
MidribDensity
y = 23.5 − 0.00744 x  R²adj = −0.0016
PL_Avg
MidribDensity
y = 10.5 + 0.00456 x  R²adj = −0.0027
gsmax
MidribDensity

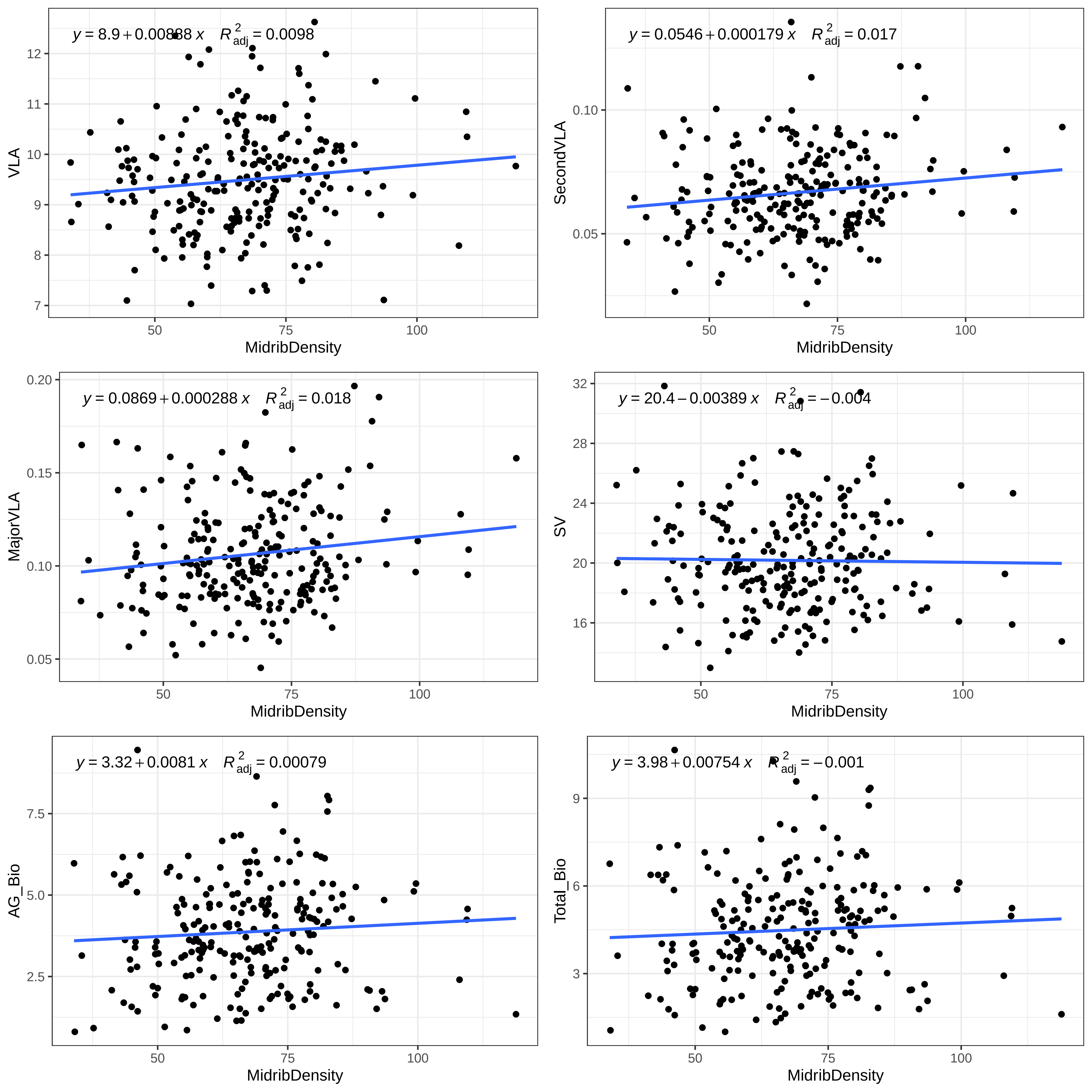

y = 8.9 + 0.00888 x  R²adj = 0.0098
VLA
MidribDensity
y = 0.0546 + 0.000179 x  R²adj = 0.017
SecondVLA
MidribDensity
y = 0.0869 + 0.000288 x  R²adj = 0.018
MajorVLA
MidribDensity
y = 20.4 − 0.00389 x  R²adj = −0.004
SV
MidribDensity
y = 3.32 + 0.0081 x  R²adj = 0.00079
AG_Bio
MidribDensity
y = 3.98 + 0.00754 x  R²adj = −0.001
Total_Bio
MidribDensity

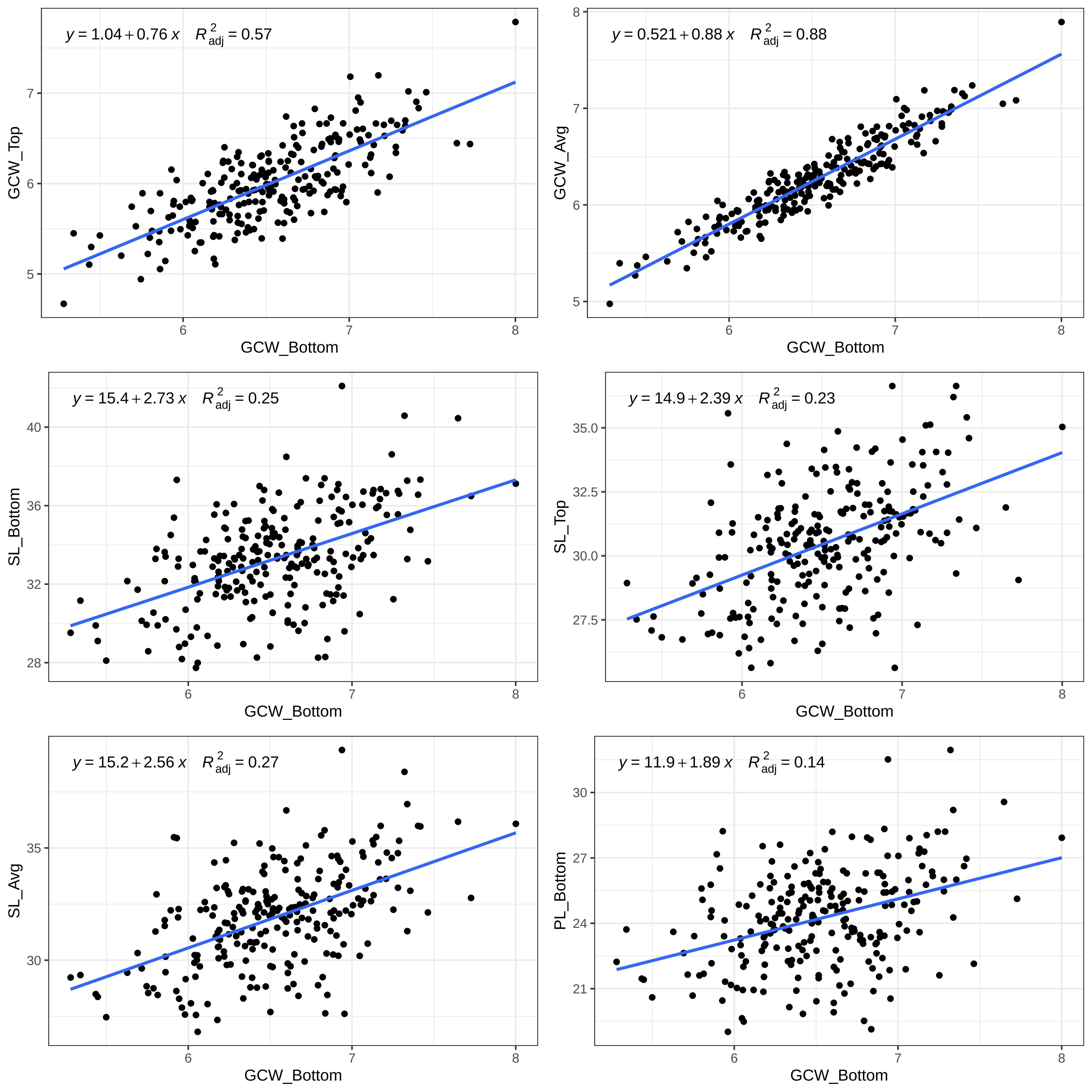

$y = 1.04 + 0.76\,x \quad R^2_{adj} = 0.57$
GCW_Top
GCW_Bottom
$y = 0.521 + 0.88\,x \quad R^2_{adj} = 0.88$
GCW_Avg
GCW_Bottom
$y = 15.4 + 2.73\,x \quad R^2_{adj} = 0.25$
SL_Bottom
GCW_Bottom
$y = 14.9 + 2.39\,x \quad R^2_{adj} = 0.23$
SL_Top
GCW_Bottom
$y = 15.2 + 2.56\,x \quad R^2_{adj} = 0.27$
SL_Avg
GCW_Bottom
$y = 11.9 + 1.89\,x \quad R^2_{adj} = 0.14$
PL_Bottom
GCW_Bottom

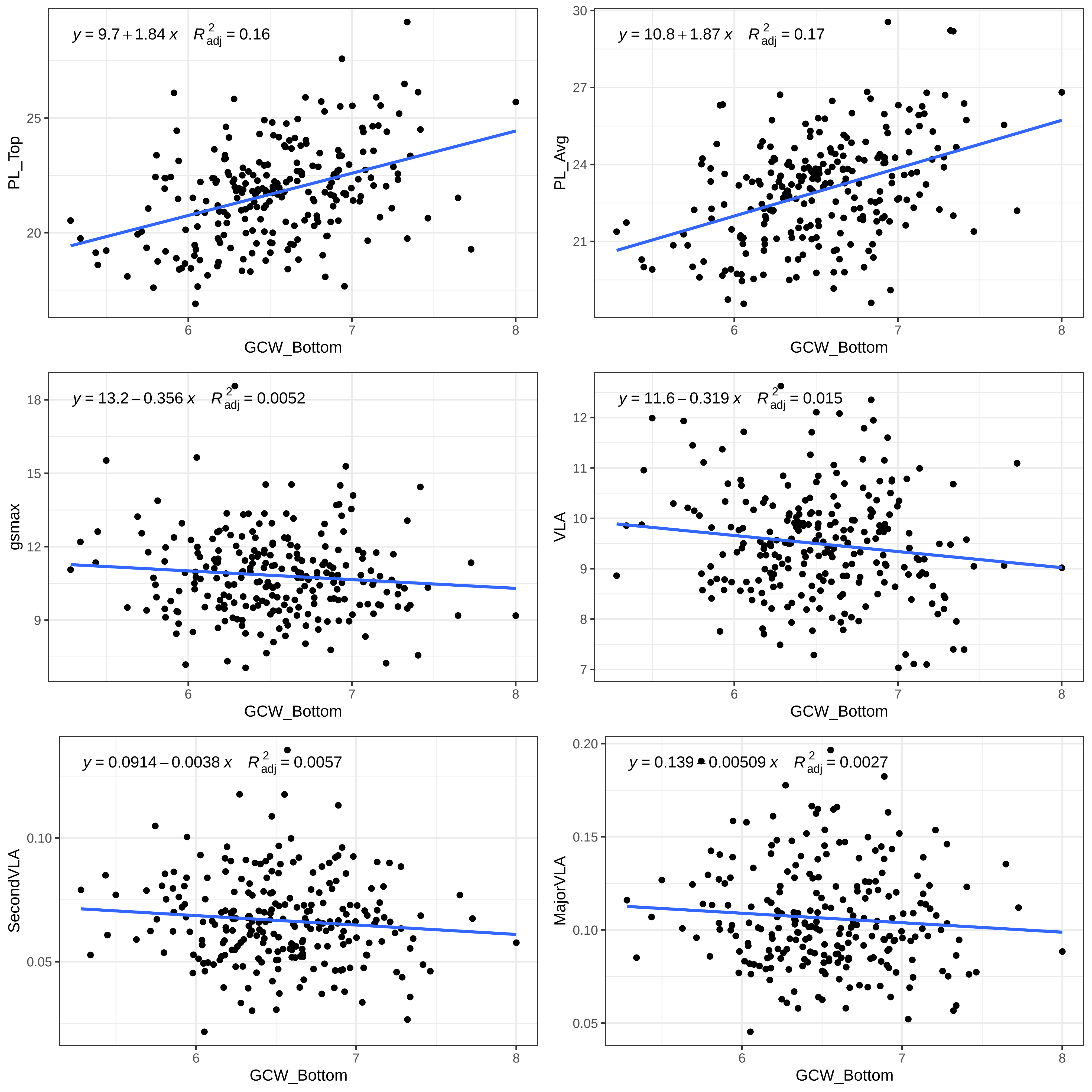

y = 9.7 + 1.84 x R2adj = 0.16
PL_Top
GCW_Bottom
y = 10.8 + 1.87 x R2adj = 0.17
PL_Avg
GCW_Bottom
y = 13.2 – 0.356 x R2adj = 0.0052
gsmax
GCW_Bottom
y = 11.6 – 0.319 x R2adj = 0.015
VLA
GCW_Bottom
y = 0.0914 – 0.0038 x R2adj = 0.0057
SecondVLA
GCW_Bottom
y = 0.139 – 0.00509 x R2adj = 0.0027
MajorVLA
GCW_Bottom

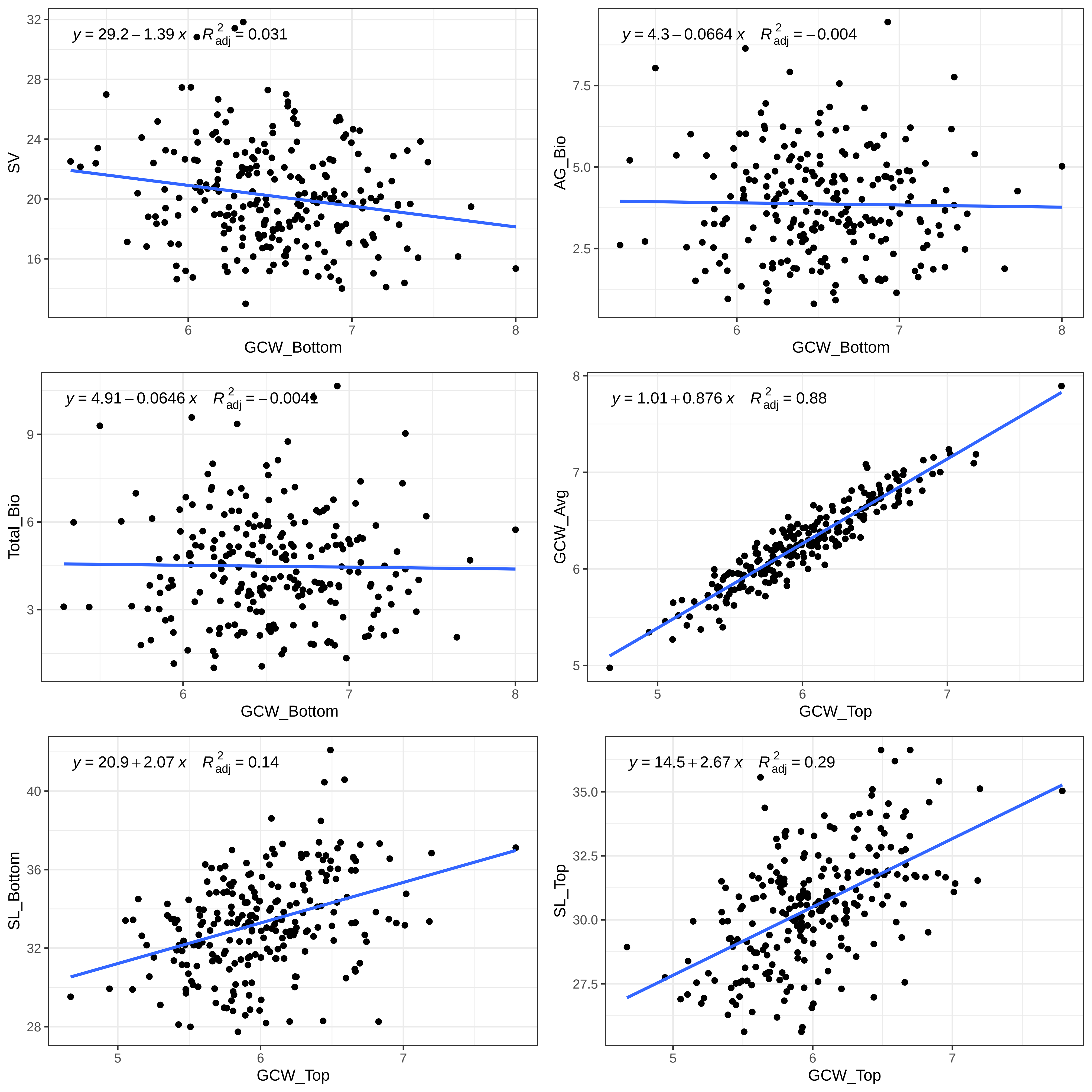

y = 29.2 − 1.39 x R2adj = 0.031
SV
GCW_Bottom
y = 4.3 − 0.0664 x R2adj = −0.004
AG_Bio
GCW_Bottom
y = 4.91 − 0.0646 x R2adj = −0.0041
Total_Bio
GCW_Bottom
y = 1.01 + 0.876 x R2adj = 0.88
GCW_Avg
GCW_Top
y = 20.9 + 2.07 x R2adj = 0.14
SL_Bottom
GCW_Top
y = 14.5 + 2.67 x R2adj = 0.29
SL_Top
GCW_Top

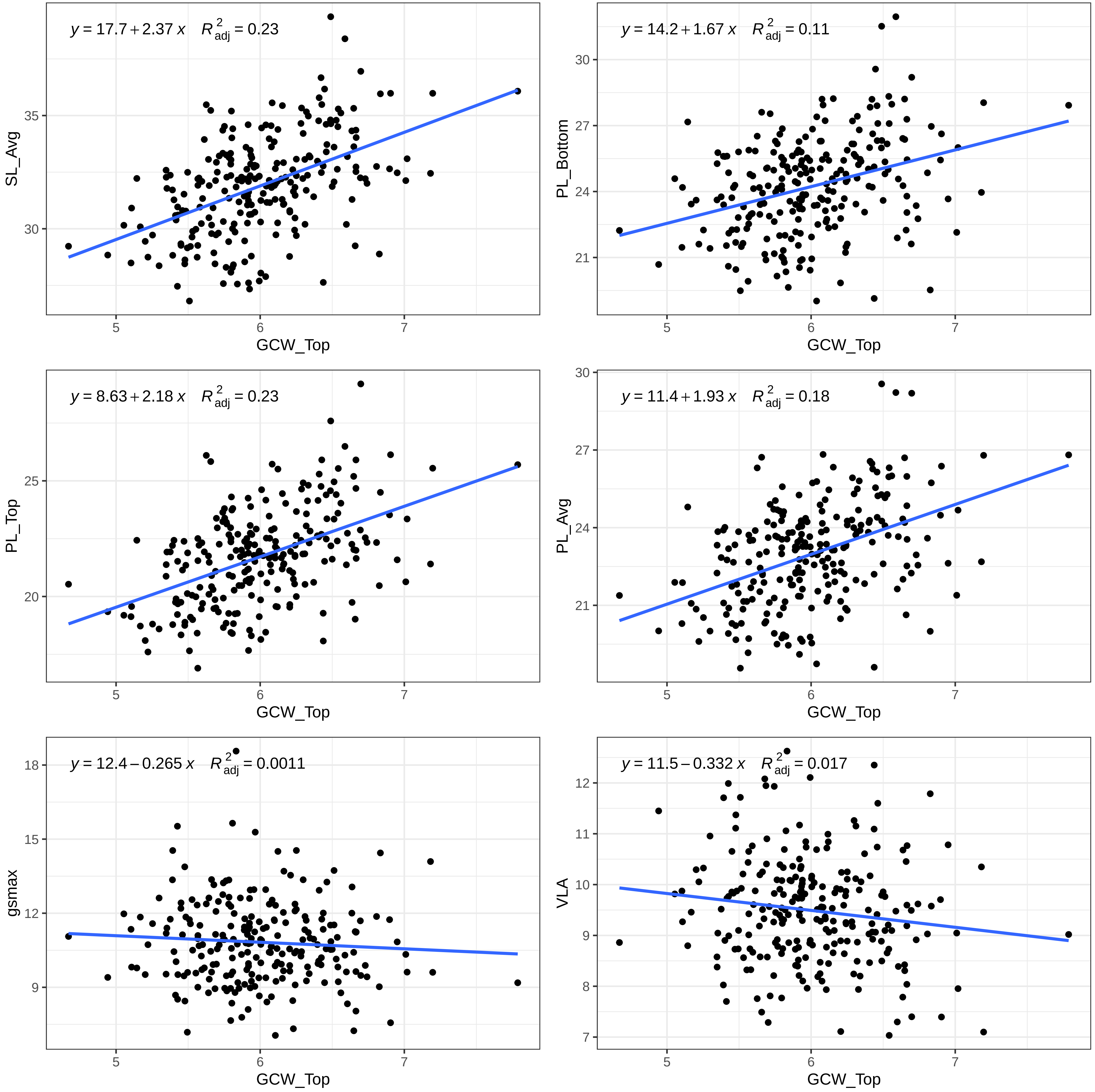

y = 17.7 + 2.37 x R adj 2 = 0.23
SL_Avg
30
35
GCW_Top
5
6
7
y = 14.2 + 1.67 x R adj 2 = 0.11
PL_Bottom
21
24
27
30
GCW_Top
y = 8.63 + 2.18 x R adj 2 = 0.23
PL_Top
20
25
GCW_Top
y = 11.4 + 1.93 x R adj 2 = 0.18
PL_Avg
21
24
27
30
GCW_Top
y = 12.4 − 0.265 x R adj 2 = 0.0011
gsmax
9
12
15
18
GCW_Top
y = 11.5 − 0.332 x R adj 2 = 0.017
VLA
7
8
9
10
11
12
GCW_Top

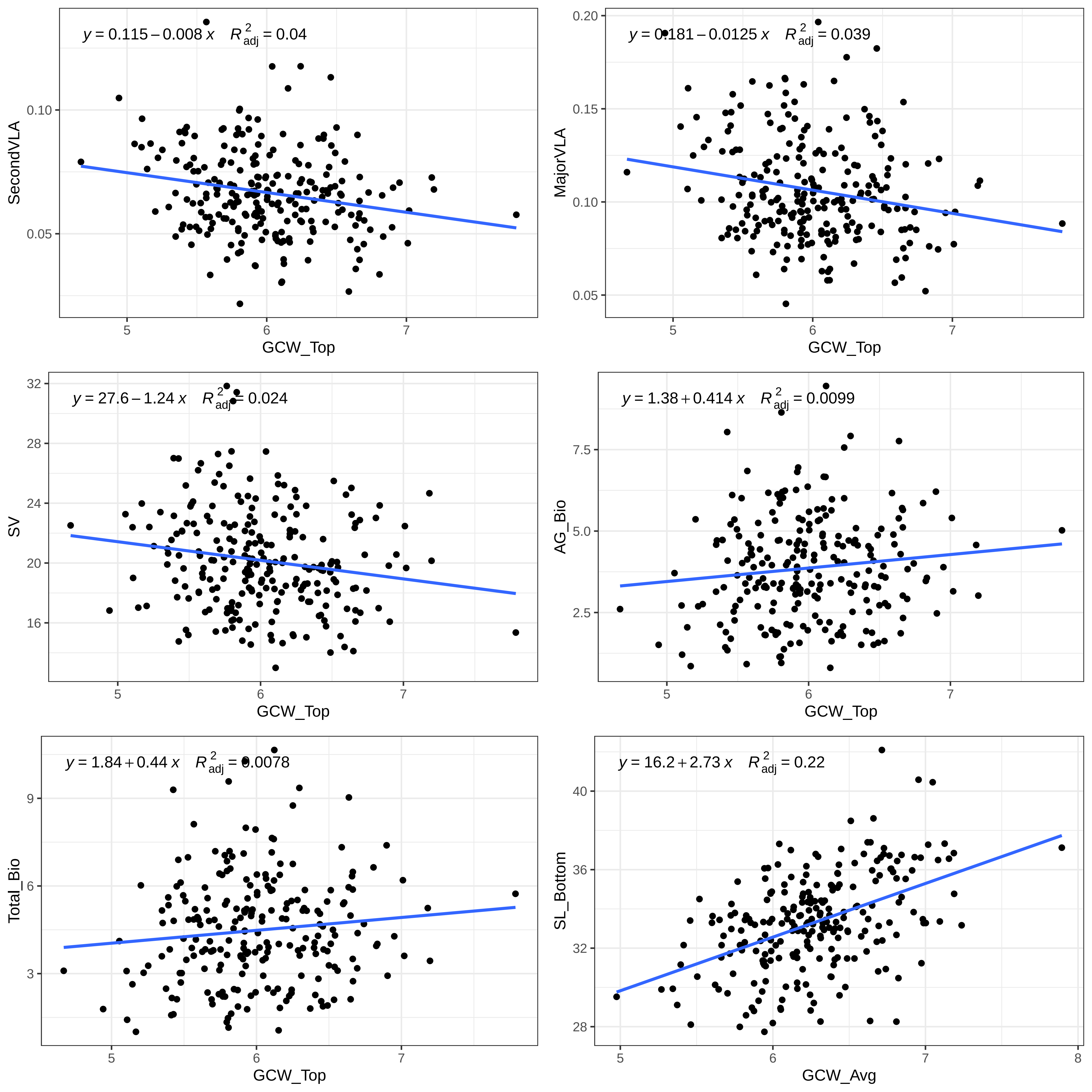

y = 0.115 − 0.008 x  R²adj = 0.04
SecondVLA
0.10
0.05
5
6
7
GCW_Top
y = 0.181 − 0.0125 x  R²adj = 0.039
MajorVLA
0.20
0.15
0.10
0.05
5
6
7
GCW_Top
y = 27.6 − 1.24 x  R²adj = 0.024
SV
32
28
24
20
16
5
6
7
GCW_Top
y = 1.38 + 0.414 x  R²adj = 0.0099
AG_Bio
7.5
5.0
2.5
5
6
7
GCW_Top
y = 1.84 + 0.44 x  R²adj = 0.0078
Total_Bio
9
6
3
5
6
7
GCW_Top
y = 16.2 + 2.73 x  R²adj = 0.22
SL_Bottom
40
36
32
28
5
6
7
8
GCW_Avg

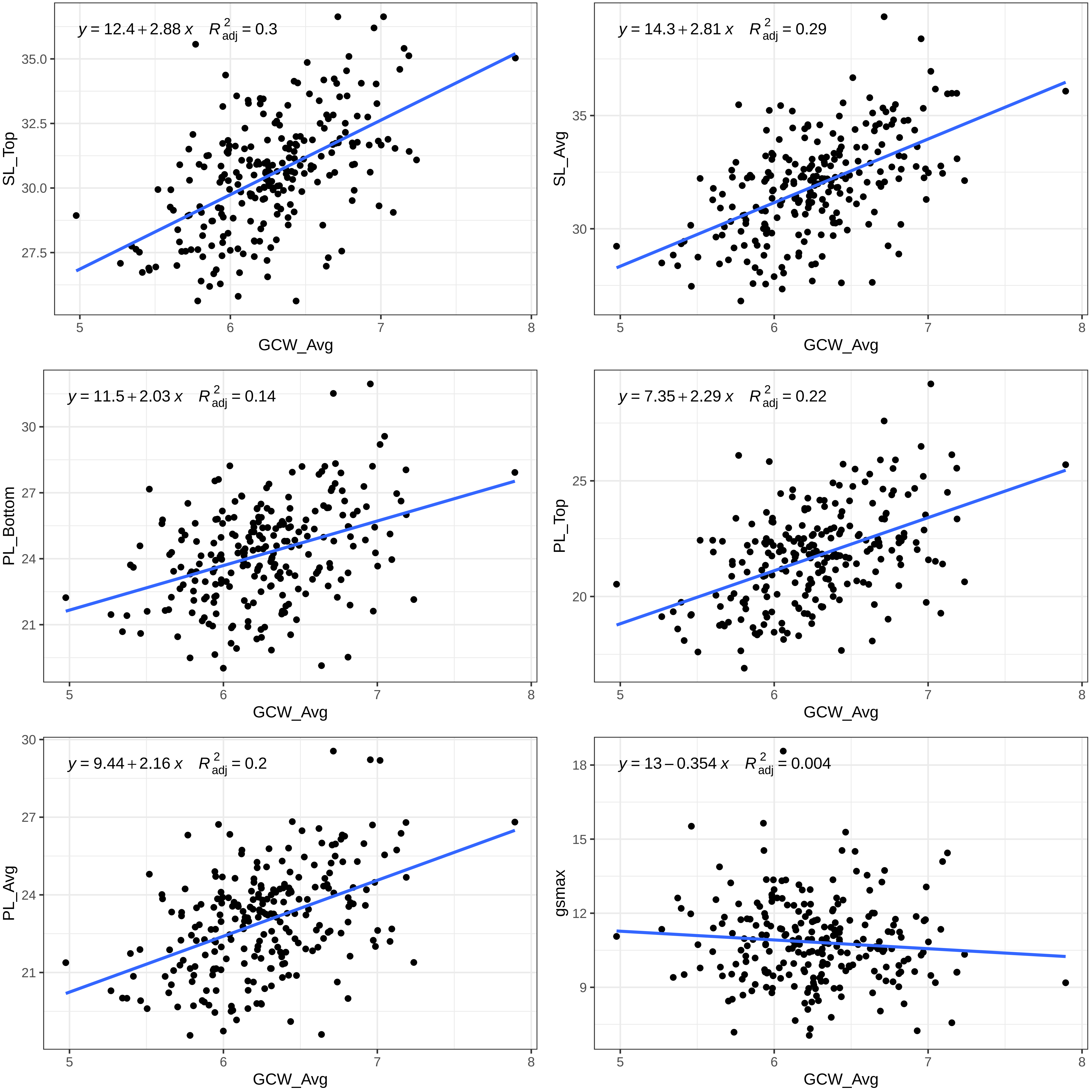

y = 12.4 + 2.88 x   R²adj = 0.3
SL_Top
GCW_Avg
y = 14.3 + 2.81 x   R²adj = 0.29
SL_Avg
GCW_Avg
y = 11.5 + 2.03 x   R²adj = 0.14
PL_Bottom
GCW_Avg
y = 7.35 + 2.29 x   R²adj = 0.22
PL_Top
GCW_Avg
y = 9.44 + 2.16 x   R²adj = 0.2
PL_Avg
GCW_Avg
y = 13 − 0.354 x   R²adj = 0.004
gsmax
GCW_Avg

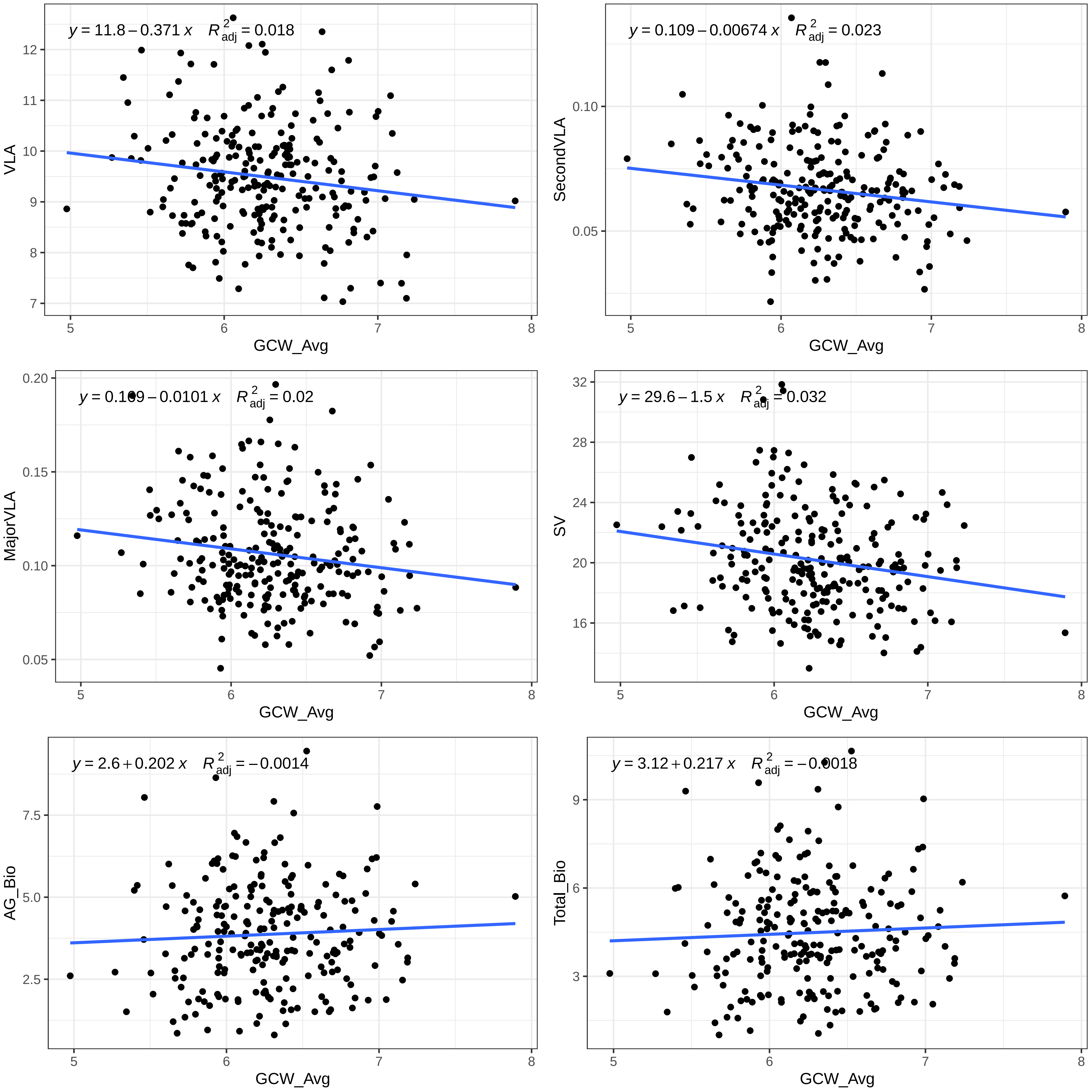

y = 11.8 − 0.371 x   R²adj = 0.018
VLA
GCW_Avg
y = 0.109 − 0.00674 x   R²adj = 0.023
SecondVLA
GCW_Avg
y = 0.109 − 0.0101 x   R²adj = 0.02
MajorVLA
GCW_Avg
y = 29.6 − 1.5 x   R²adj = 0.032
SV
GCW_Avg
y = 2.6 + 0.202 x   R²adj = −0.0014
AG_Bio
GCW_Avg
y = 3.12 + 0.217 x   R²adj = −0.0018
Total_Bio
GCW_Avg

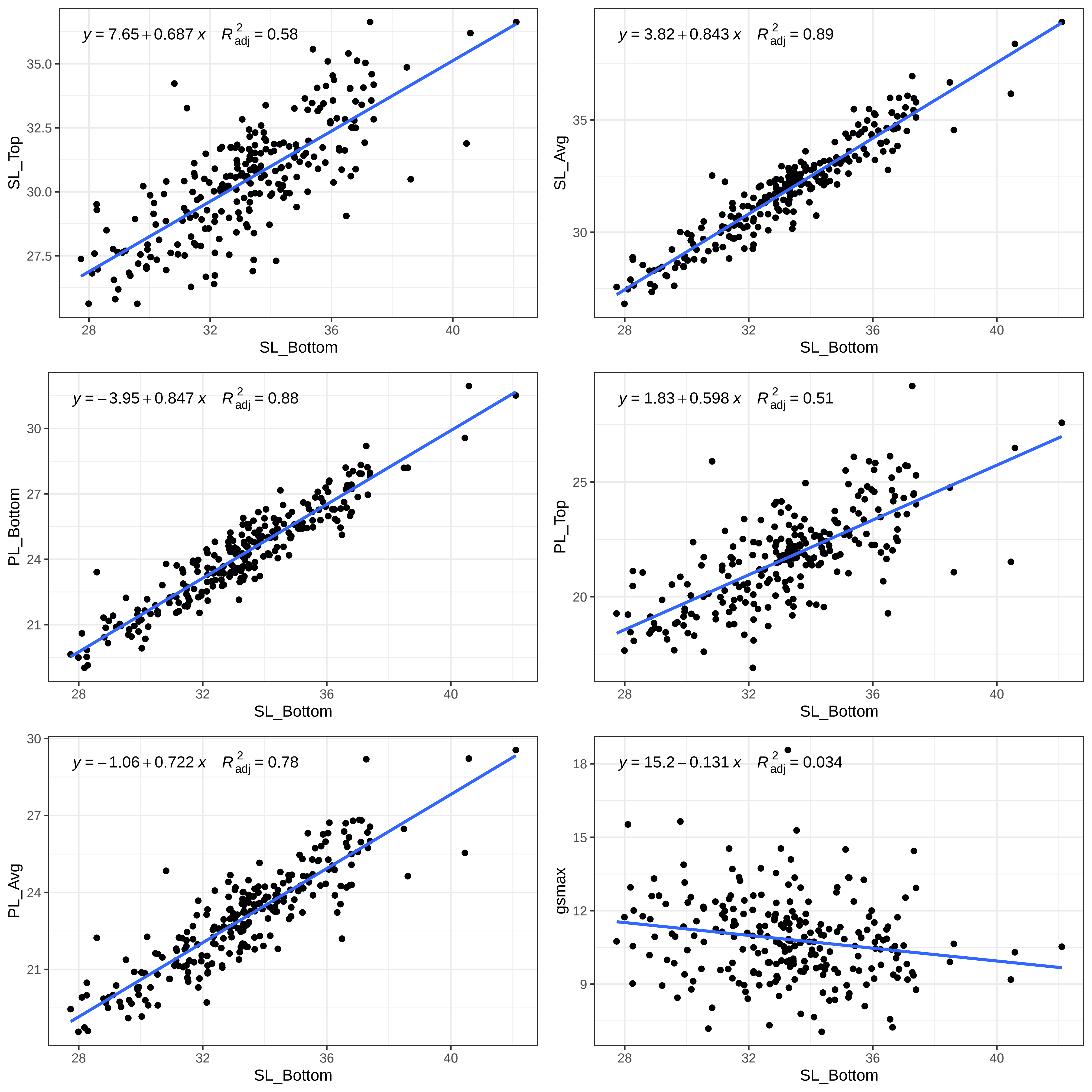

y = 7.65 + 0.687 x    R²adj = 0.58
SL_Top
SL_Bottom
y = 3.82 + 0.843 x    R²adj = 0.89
SL_Avg
SL_Bottom
y = −3.95 + 0.847 x    R²adj = 0.88
PL_Bottom
SL_Bottom
y = 1.83 + 0.598 x    R²adj = 0.51
PL_Top
SL_Bottom
y = −1.06 + 0.722 x    R²adj = 0.78
PL_Avg
SL_Bottom
y = 15.2 − 0.131 x    R²adj = 0.034
gsmax
SL_Bottom

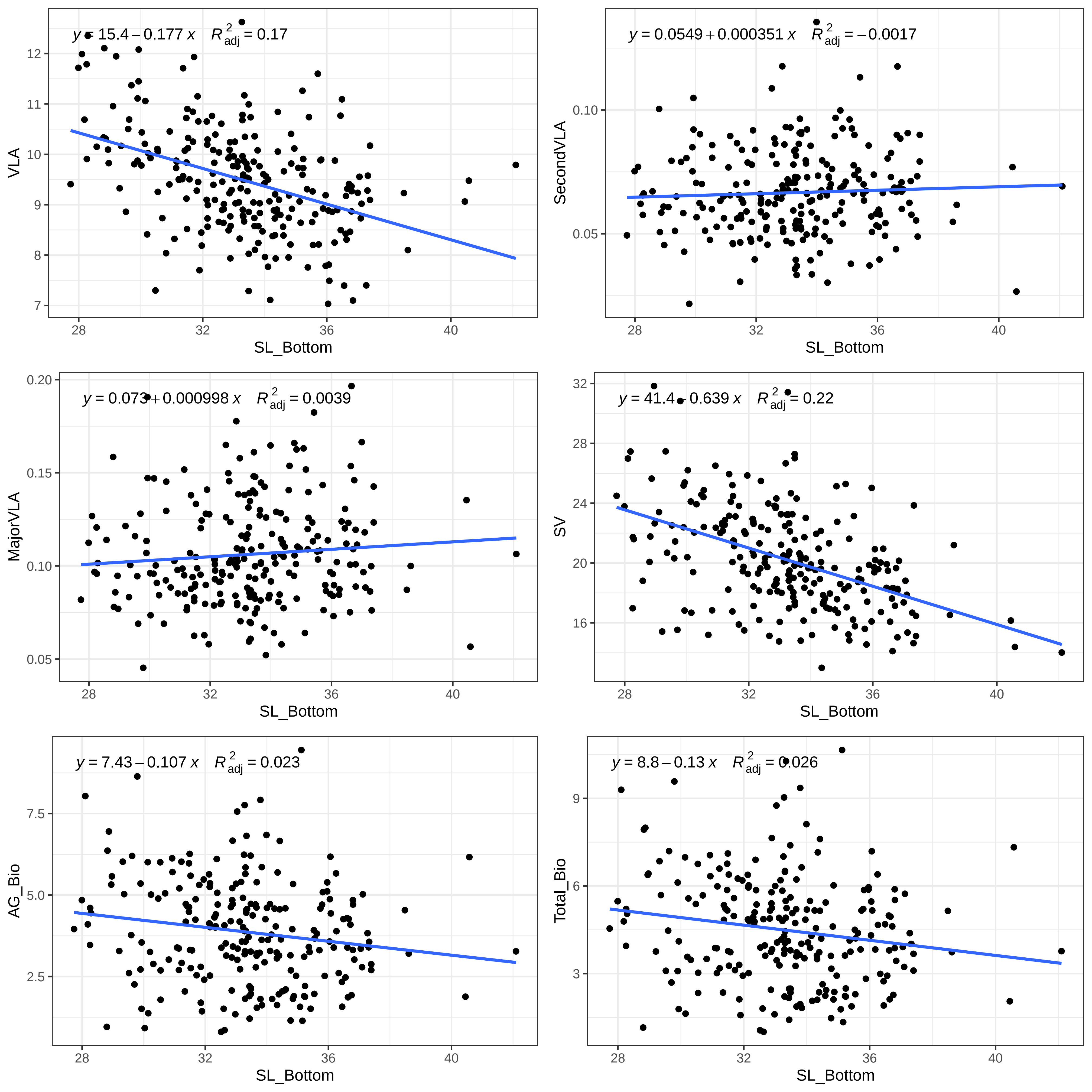

y = 15.4 − 0.177 x   R²adj = 0.17
VLA
SL_Bottom
y = 0.0549 + 0.000351 x   R²adj = −0.0017
SecondVLA
SL_Bottom
y = 0.073 + 0.000998 x   R²adj = 0.0039
MajorVLA
SL_Bottom
y = 41.4 − 0.639 x   R²adj = 0.22
SV
SL_Bottom
y = 7.43 − 0.107 x   R²adj = 0.023
AG_Bio
SL_Bottom
y = 8.8 − 0.13 x   R²adj = 0.026
Total_Bio
SL_Bottom

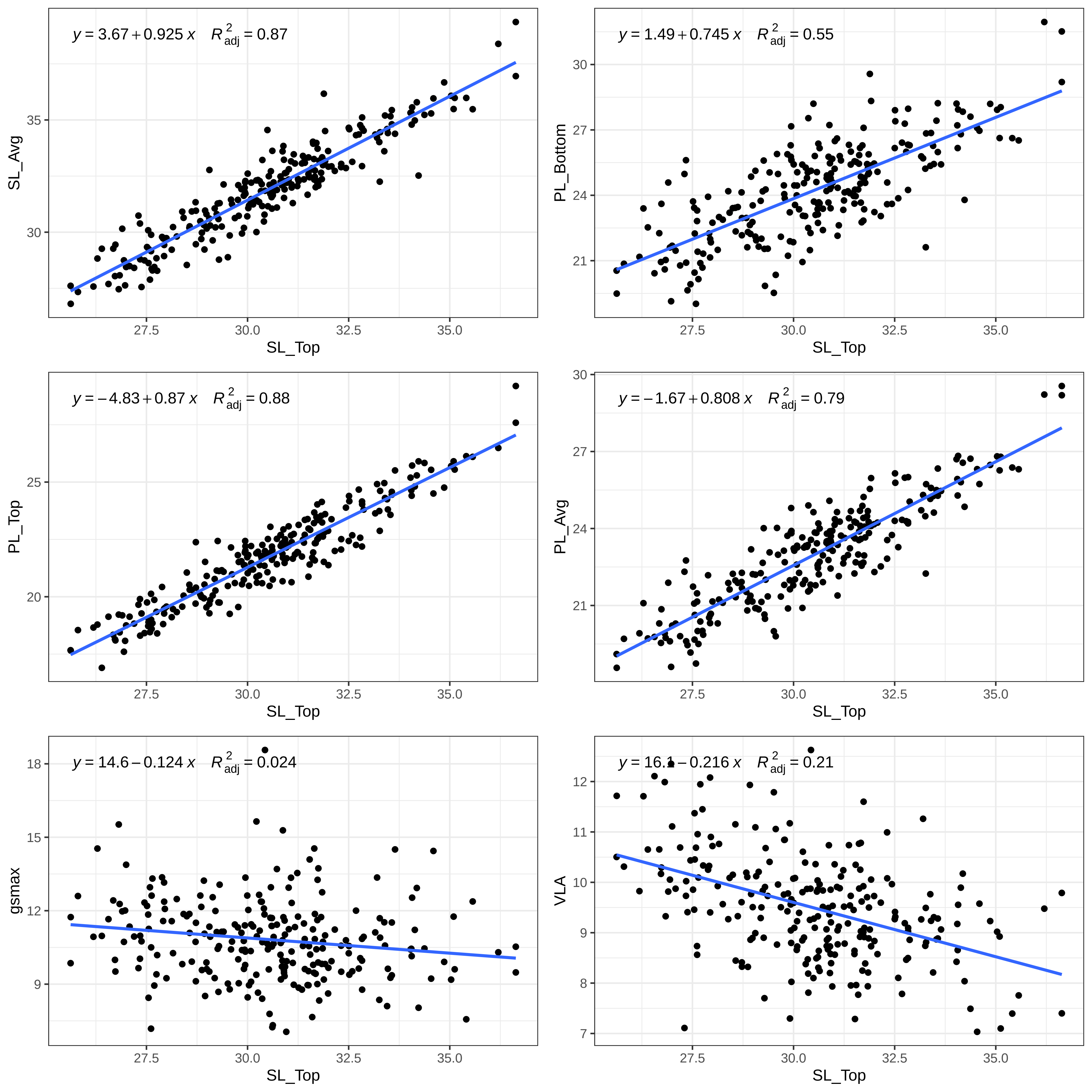

y = 3.67 + 0.925 x R2adj = 0.87
SL_Avg
SL_Top
y = 1.49 + 0.745 x R2adj = 0.55
PL_Bottom
SL_Top
y = −4.83 + 0.87 x R2adj = 0.88
PL_Top
SL_Top
y = −1.67 + 0.808 x R2adj = 0.79
PL_Avg
SL_Top
y = 14.6 − 0.124 x R2adj = 0.024
gsmax
SL_Top
y = 16.1 − 0.216 x R2adj = 0.21
VLA
SL_Top

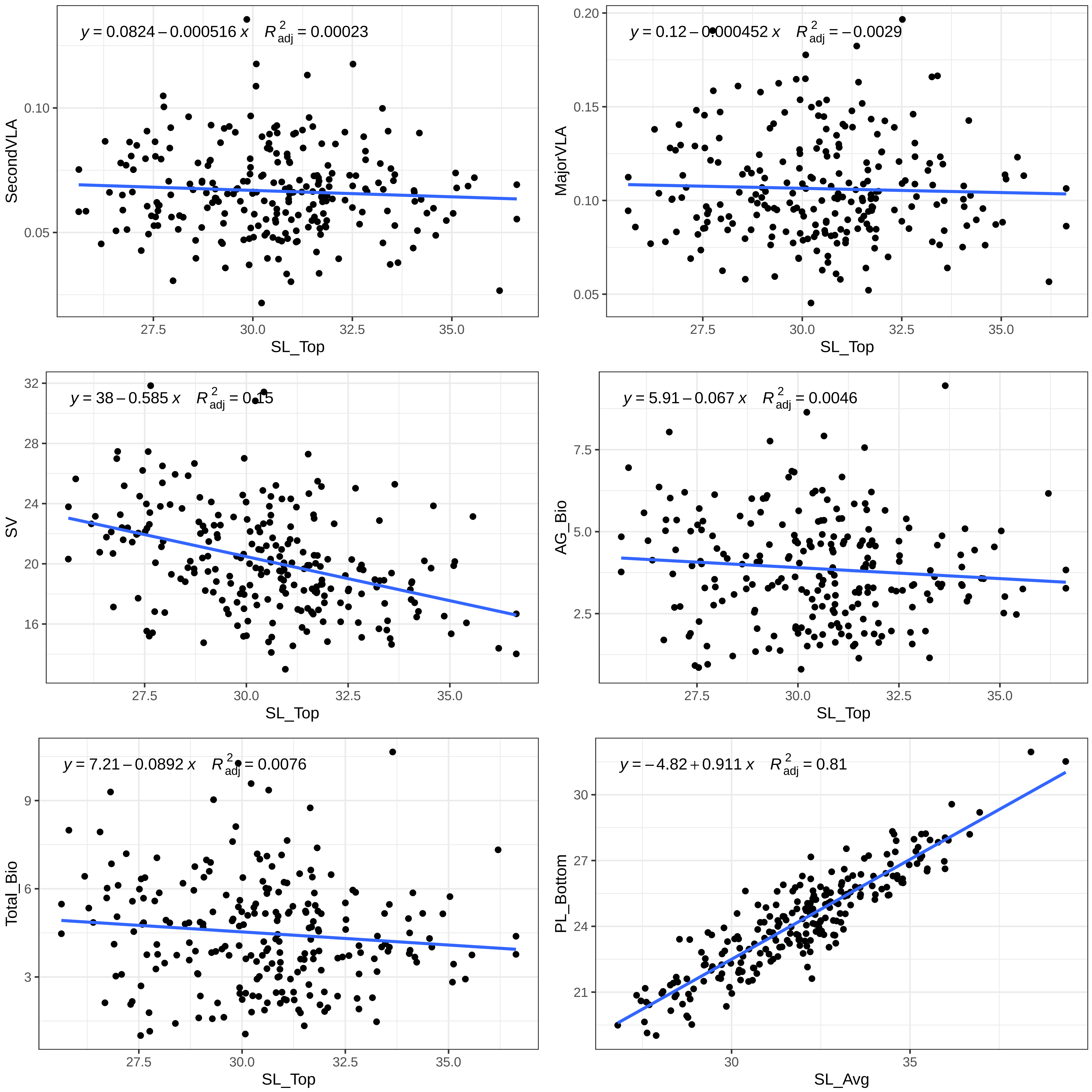
y = 0.0824 − 0.000516 x   R²adj = 0.00023
SecondVLA
SL_Top
y = 0.12 − 0.000452 x   R²adj = −0.0029
MajorVLA
SL_Top
y = 38 − 0.585 x   R²adj = 0.15
SV
SL_Top
y = 5.91 − 0.067 x   R²adj = 0.0046
AG_Bio
SL_Top
y = 7.21 − 0.0892 x   R²adj = 0.0076
Total_Bio
SL_Top
y = −4.82 + 0.911 x   R²adj = 0.81
PL_Bottom
SL_Avg

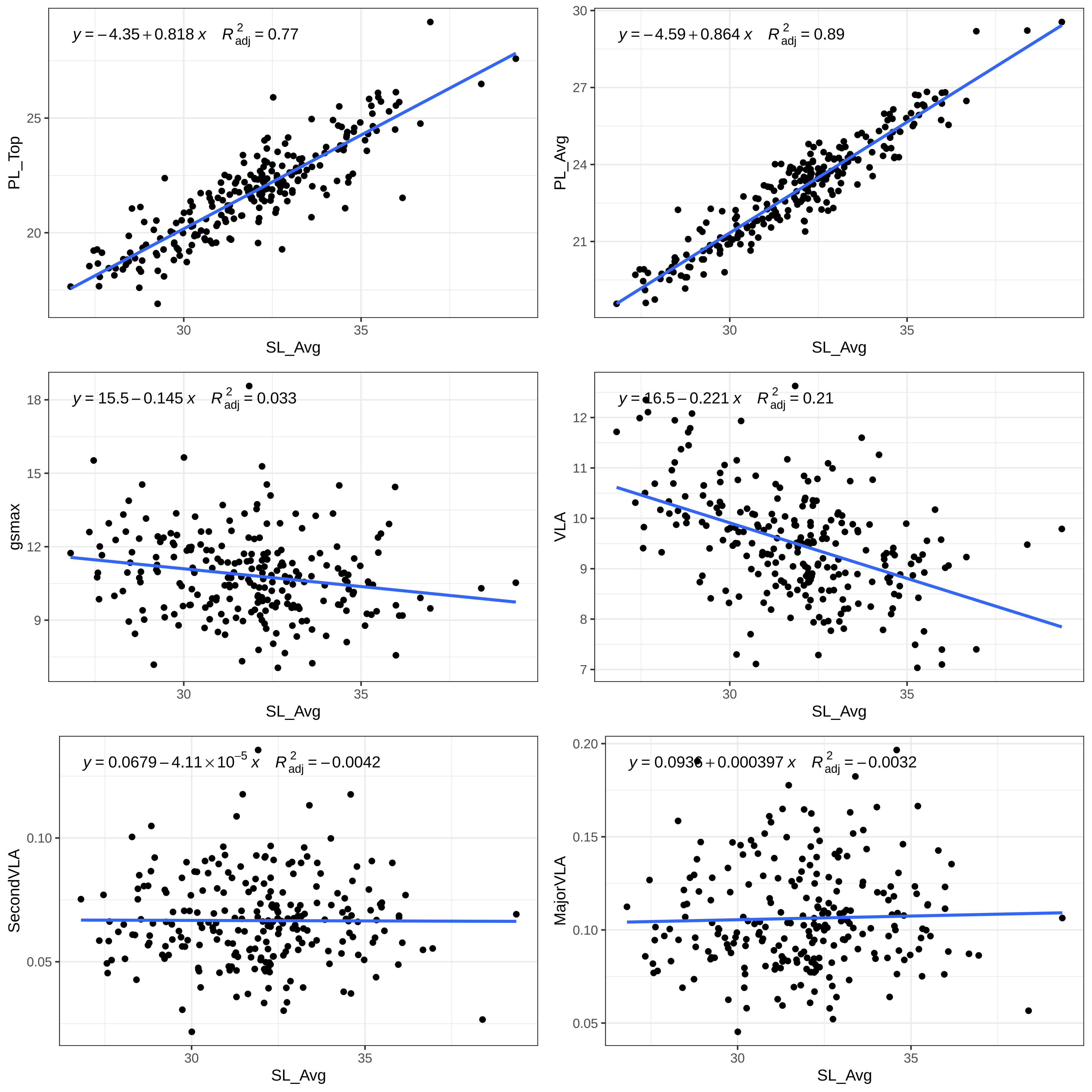

$y = -4.35 + 0.818\,x \quad R^2_{adj} = 0.77$
PL_Top
SL_Avg
$y = -4.59 + 0.864\,x \quad R^2_{adj} = 0.89$
PL_Avg
SL_Avg
$y = 15.5 - 0.145\,x \quad R^2_{adj} = 0.033$
gsmax
SL_Avg
$y = 16.5 - 0.221\,x \quad R^2_{adj} = 0.21$
VLA
SL_Avg
$y = 0.0679 - 4.11 \times 10^{-5}\,x \quad R^2_{adj} = -0.0042$
SecondVLA
SL_Avg
$y = 0.0936 + 0.000397\,x \quad R^2_{adj} = -0.0032$
MajorVLA
SL_Avg

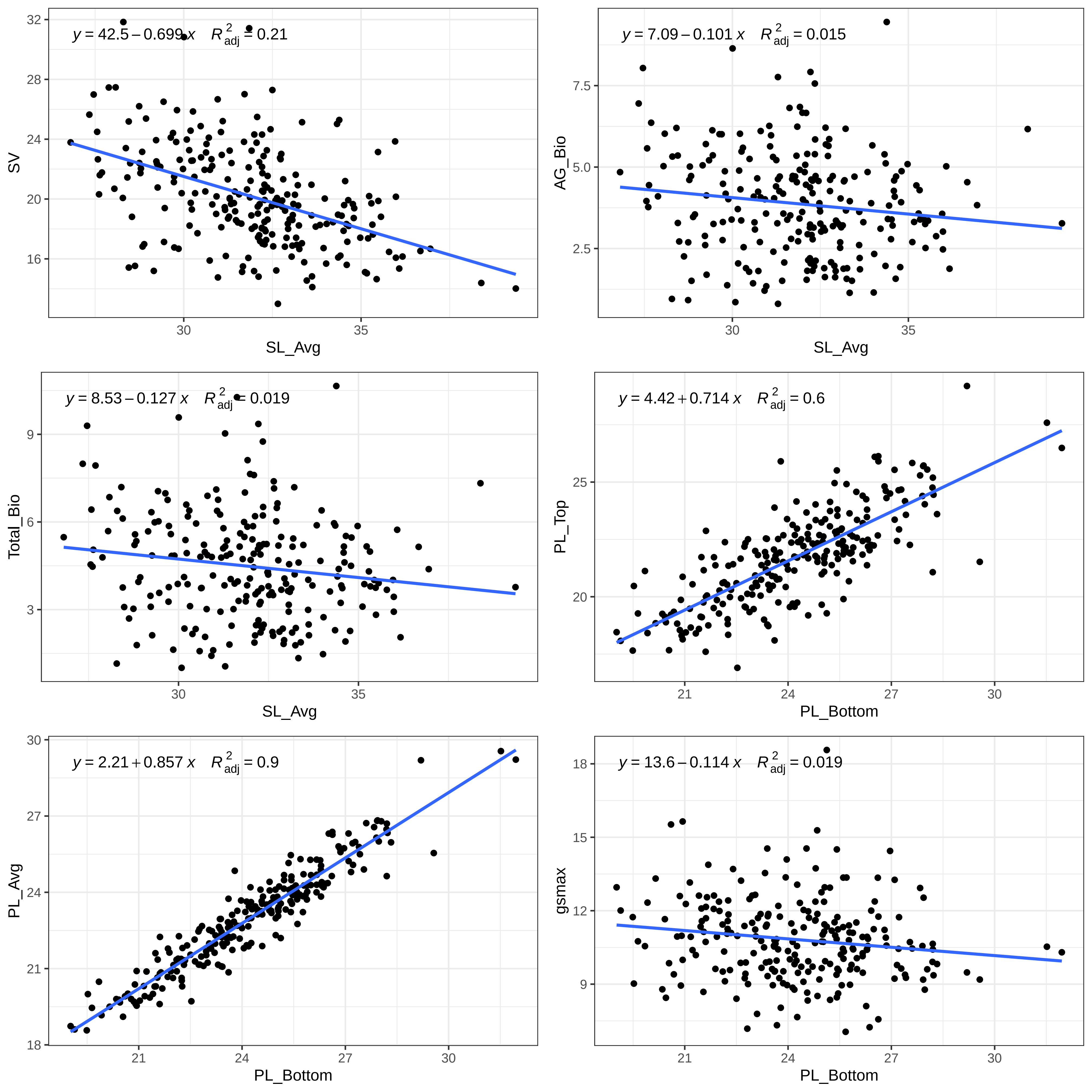

y = 42.5 − 0.699 x  R²adj = 0.21
SV
SL_Avg
y = 7.09 − 0.101 x  R²adj = 0.015
AG_Bio
SL_Avg
y = 8.53 − 0.127 x  R²adj = 0.019
Total_Bio
SL_Avg
y = 4.42 + 0.714 x  R²adj = 0.6
PL_Top
PL_Bottom
y = 2.21 + 0.857 x  R²adj = 0.9
PL_Avg
PL_Bottom
y = 13.6 − 0.114 x  R²adj = 0.019
gsmax
PL_Bottom

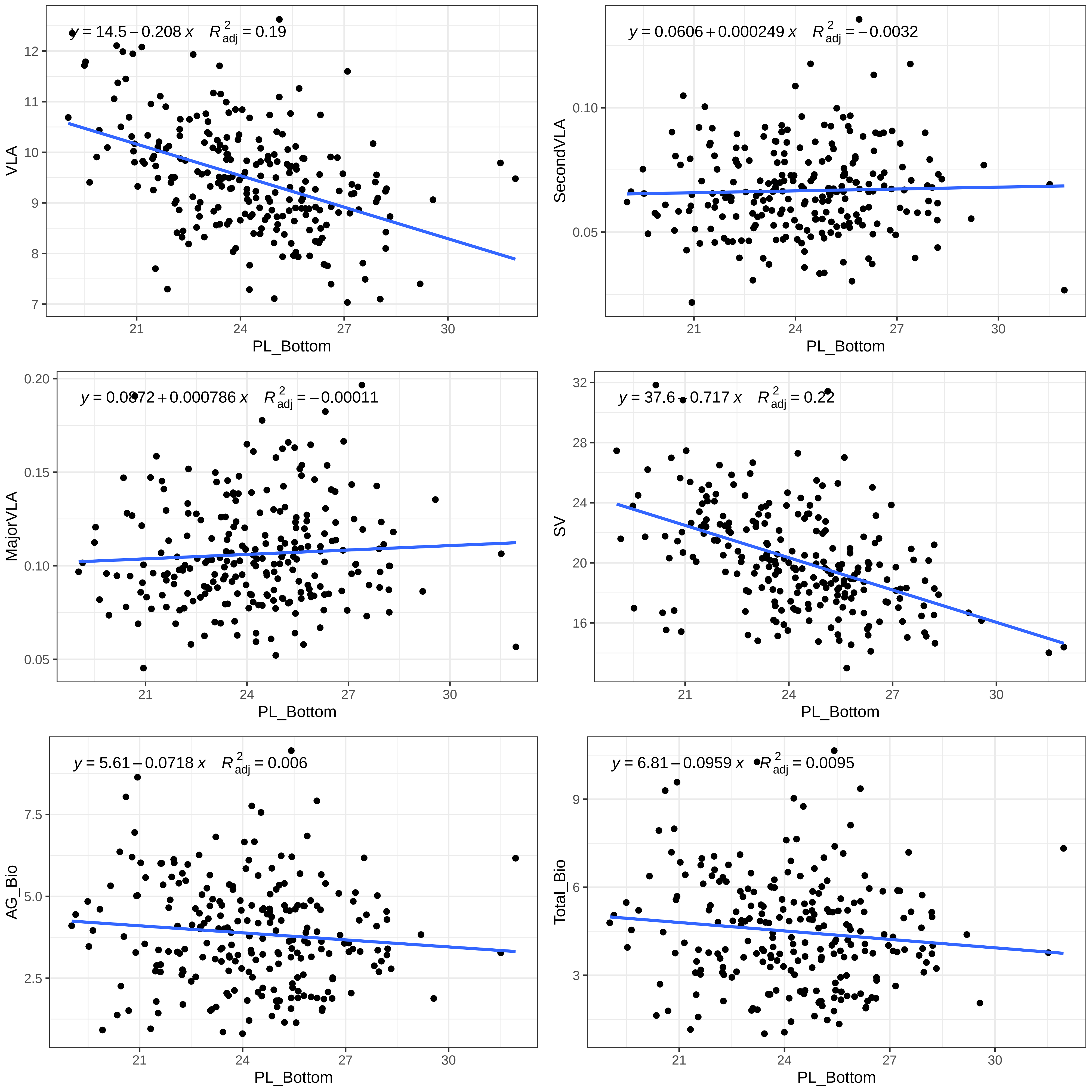

y = 14.5 − 0.208 x  R²adj = 0.19
VLA
PL_Bottom
y = 0.0606 + 0.000249 x  R²adj = −0.0032
SecondVLA
PL_Bottom
y = 0.0872 + 0.000786 x  R²adj = −0.00011
MajorVLA
PL_Bottom
y = 37.6 − 0.717 x  R²adj = 0.22
SV
PL_Bottom
y = 5.61 − 0.0718 x  R²adj = 0.006
AG_Bio
PL_Bottom
y = 6.81 − 0.0959 x  R²adj = 0.0095
Total_Bio
PL_Bottom

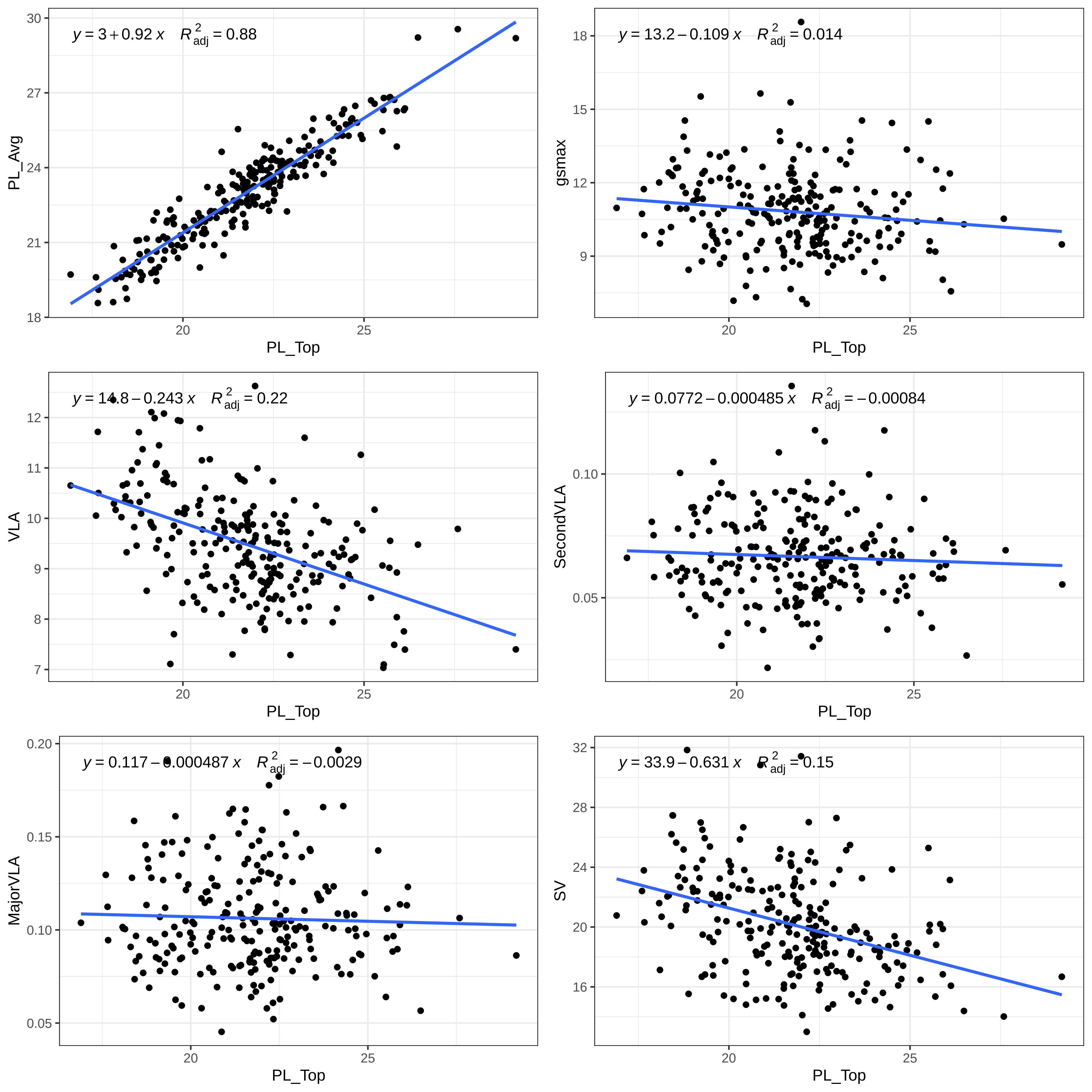
y = 3 + 0.92 x    R²adj = 0.88
PL_Avg
PL_Top
y = 13.2 − 0.109 x    R²adj = 0.014
gsmax
PL_Top
y = 14.8 − 0.243 x    R²adj = 0.22
VLA
PL_Top
y = 0.0772 − 0.000485 x    R²adj = −0.00084
SecondVLA
PL_Top
y = 0.117 − 0.000487 x    R²adj = −0.0029
MajorVLA
PL_Top
y = 33.9 − 0.631 x    R²adj = 0.15
SV
PL_Top

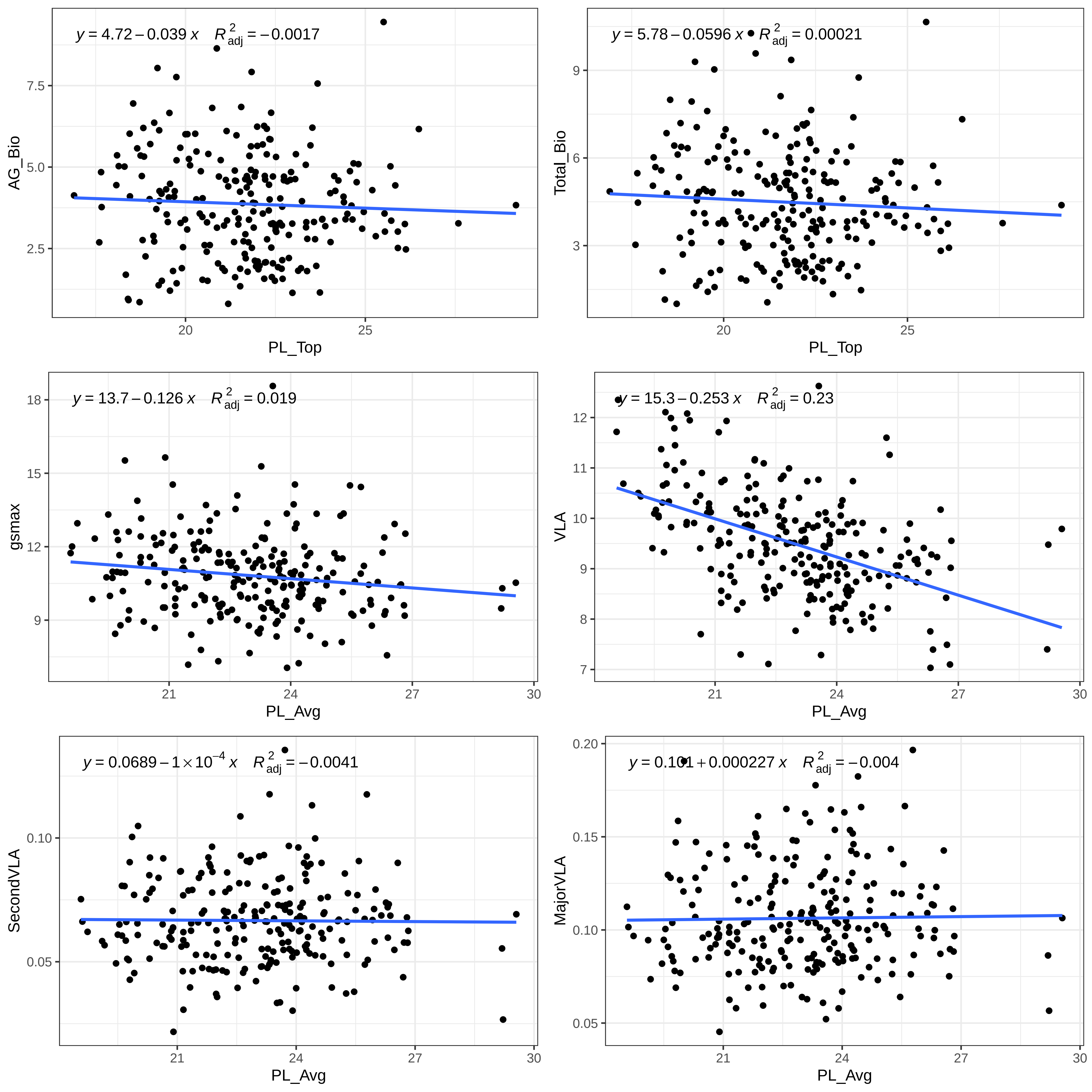

y = 4.72 − 0.039 x   R²adj = −0.0017
AG_Bio
7.5
5.0
2.5
20
25
PL_Top
y = 5.78 − 0.0596 x   R²adj = 0.00021
Total_Bio
9
6
3
20
25
PL_Top
y = 13.7 − 0.126 x   R²adj = 0.019
gsmax
18
15
12
9
21
24
27
30
PL_Avg
y = 15.3 − 0.253 x   R²adj = 0.23
VLA
12
11
10
9
8
7
21
24
27
30
PL_Avg
y = 0.0689 − 1 × 10⁻⁴ x   R²adj = −0.0041
SecondVLA
0.10
0.05
21
24
27
30
PL_Avg
y = 0.101 + 0.000227 x   R²adj = −0.004
MajorVLA
0.20
0.15
0.10
0.05
21
24
27
30
PL_Avg

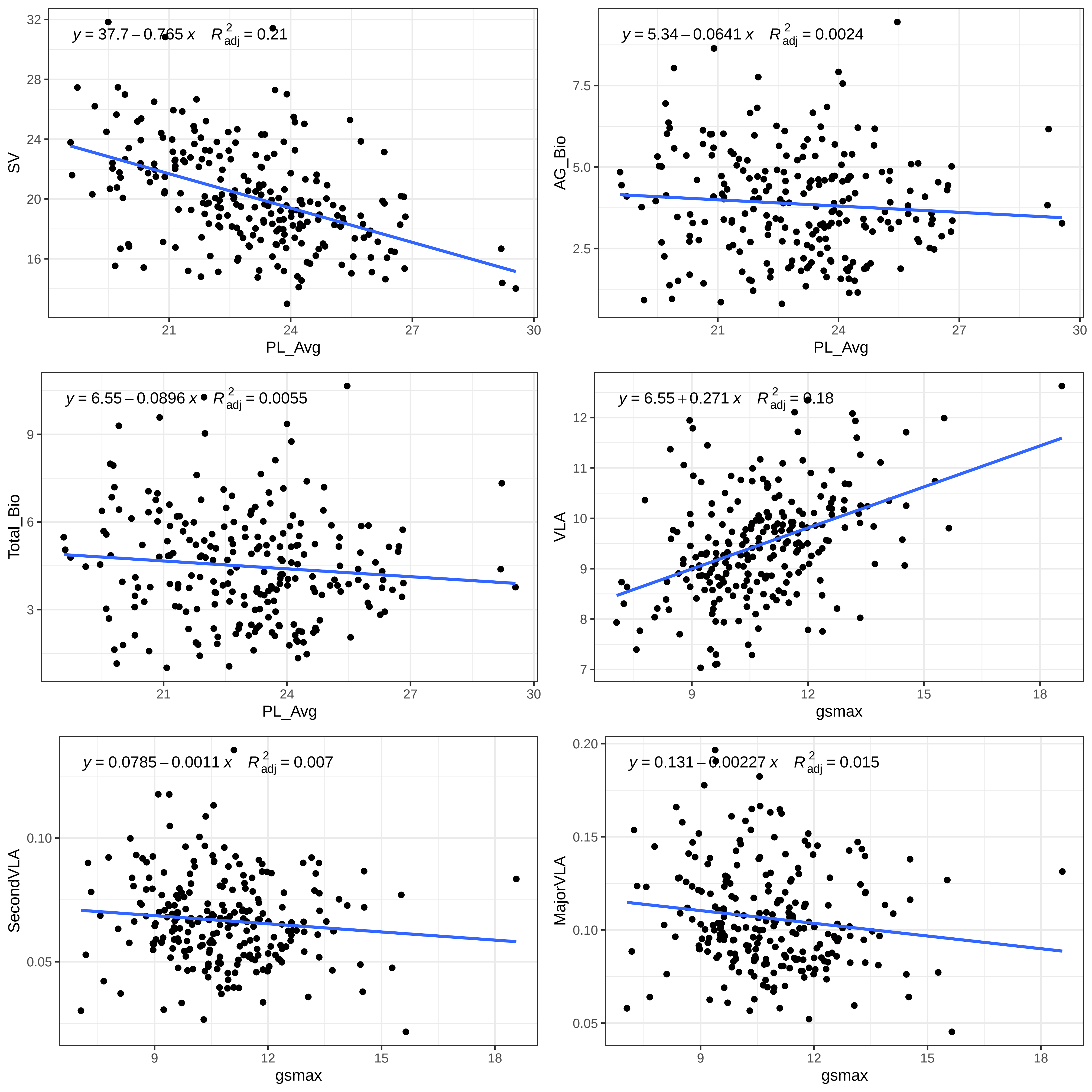

y = 37.7 − 0.765 x   R2adj = 0.21
SV
PL_Avg
y = 5.34 − 0.0641 x   R2adj = 0.0024
AG_Bio
PL_Avg
y = 6.55 − 0.0896 x   R2adj = 0.0055
Total_Bio
PL_Avg
y = 6.55 + 0.271 x   R2adj = 0.18
VLA
gsmax
y = 0.0785 − 0.0011 x   R2adj = 0.007
SecondVLA
gsmax
y = 0.131 − 0.00227 x   R2adj = 0.015
MajorVLA
gsmax

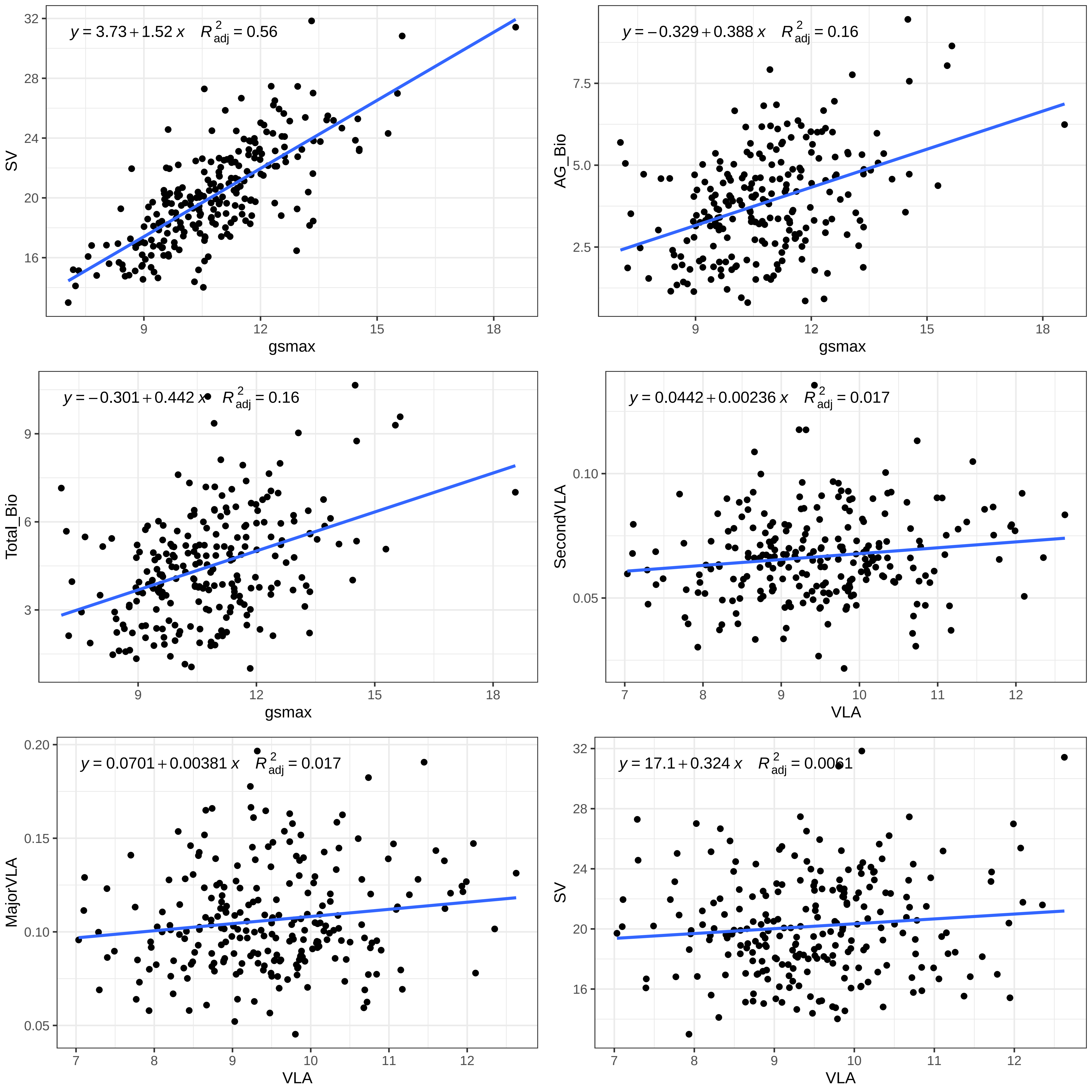

y = 3.73 + 1.52 x    R2adj = 0.56
SV
gsmax
y = −0.329 + 0.388 x    R2adj = 0.16
AG_Bio
gsmax
y = −0.301 + 0.442 x    R2adj = 0.16
Total_Bio
gsmax
y = 0.0442 + 0.00236 x    R2adj = 0.017
SecondVLA
VLA
y = 0.0701 + 0.00381 x    R2adj = 0.017
MajorVLA
VLA
y = 17.1 + 0.324 x    R2adj = 0.0061
SV
VLA

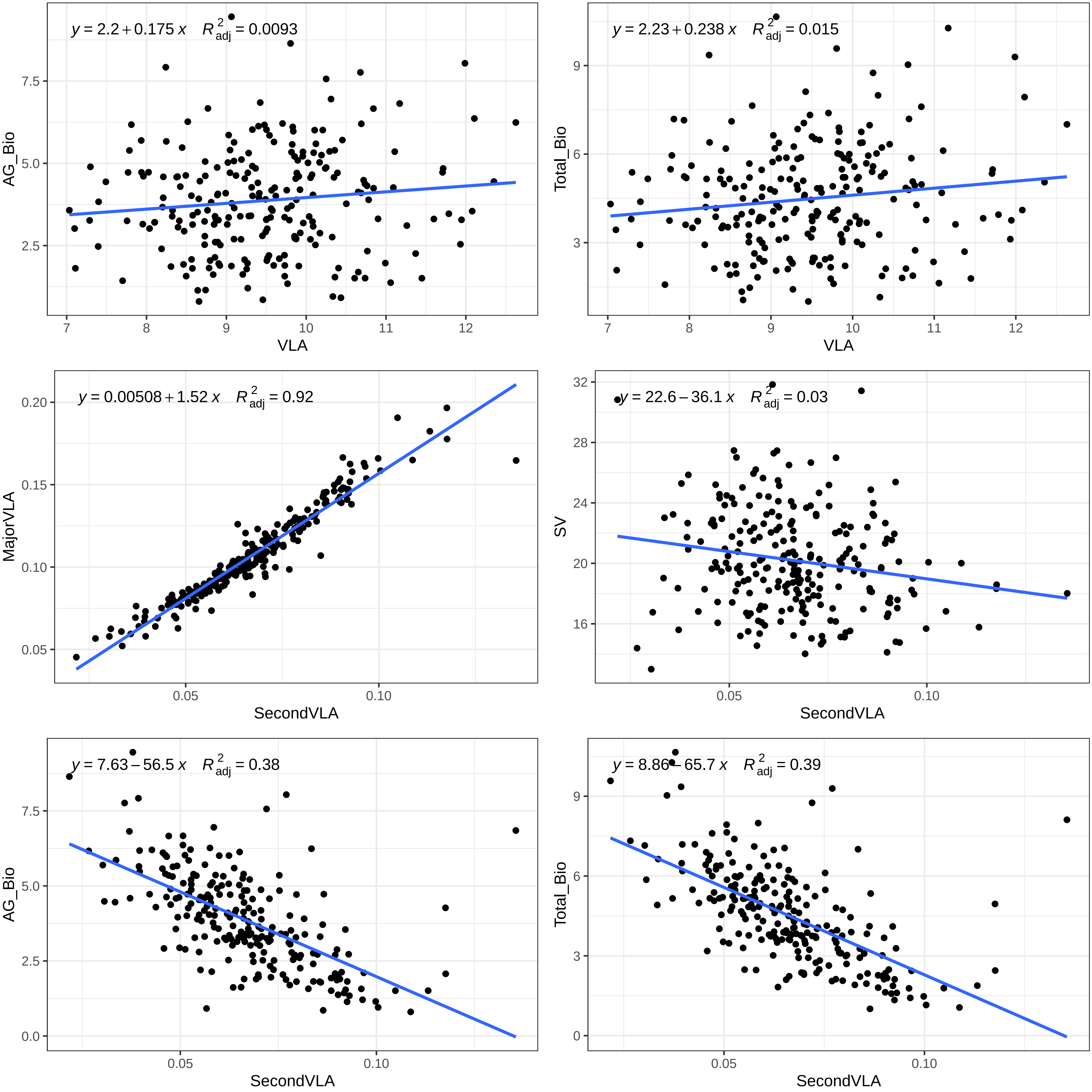

y = 2.2 + 0.175 x R²adj = 0.0093
AG_Bio
VLA
y = 2.23 + 0.238 x R²adj = 0.015
Total_Bio
VLA
y = 0.00508 + 1.52 x R²adj = 0.92
MajorVLA
SecondVLA
y = 22.6 – 36.1 x R²adj = 0.03
SV
SecondVLA
y = 7.63 – 56.5 x R²adj = 0.38
AG_Bio
SecondVLA
y = 8.86 – 65.7 x R²adj = 0.39
Total_Bio
SecondVLA

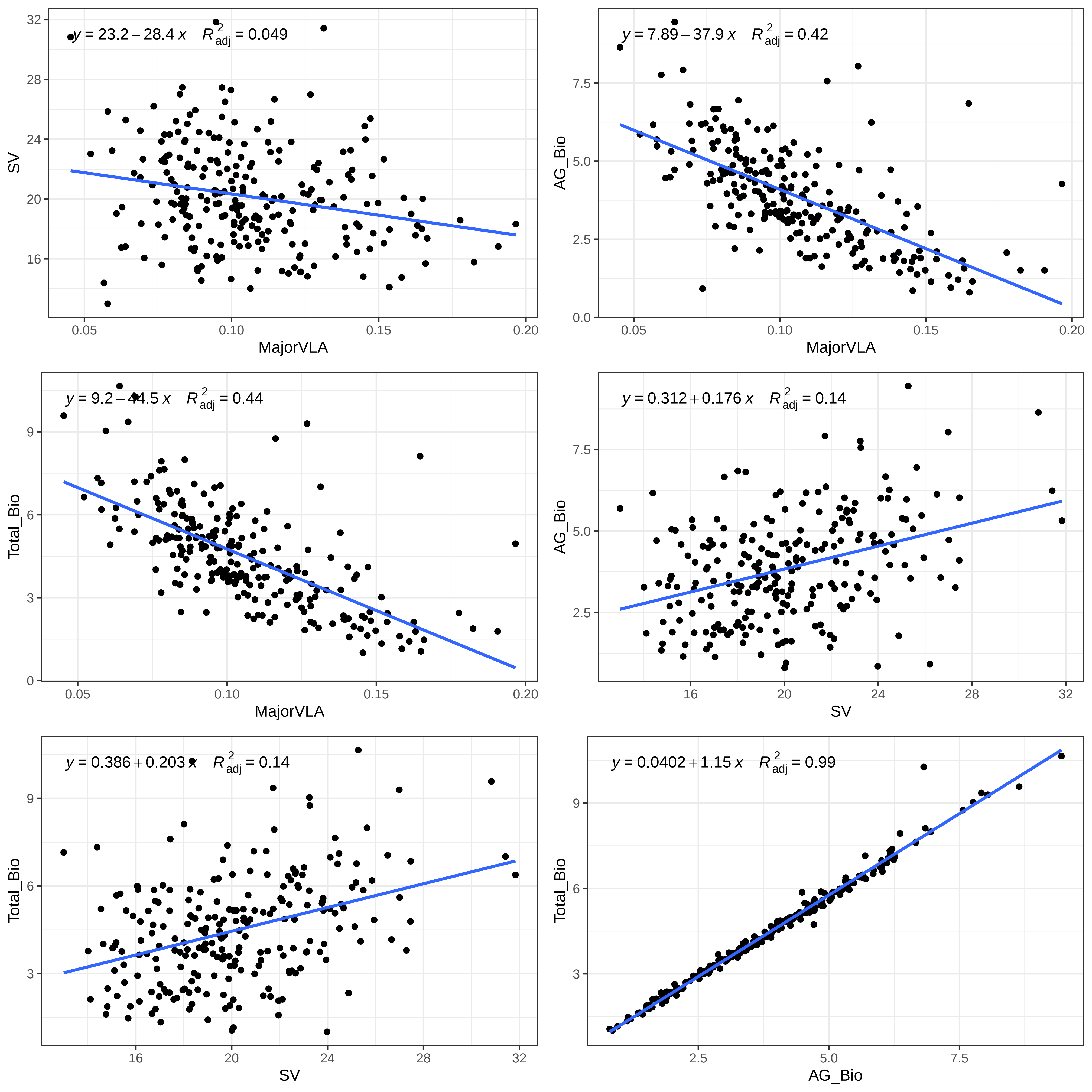

y = 23.2 − 28.4 x   R2adj = 0.049
SV
MajorVLA
y = 7.89 − 37.9 x   R2adj = 0.42
AG_Bio
MajorVLA
y = 9.2 − 44.5 x   R2adj = 0.44
Total_Bio
MajorVLA
y = 0.312 + 0.176 x   R2adj = 0.14
AG_Bio
SV
y = 0.386 + 0.203 x   R2adj = 0.14
Total_Bio
SV
y = 0.0402 + 1.15 x   R2adj = 0.99
Total_Bio
AG_Bio

**Figure S5:** *Principal component analysis (PCA) of all measured leaf traits using estimated marginal means for each trait. For stomatal density, length, pore length, and guard cell width, trait values are included separately for the top and bottom of the leaf.*

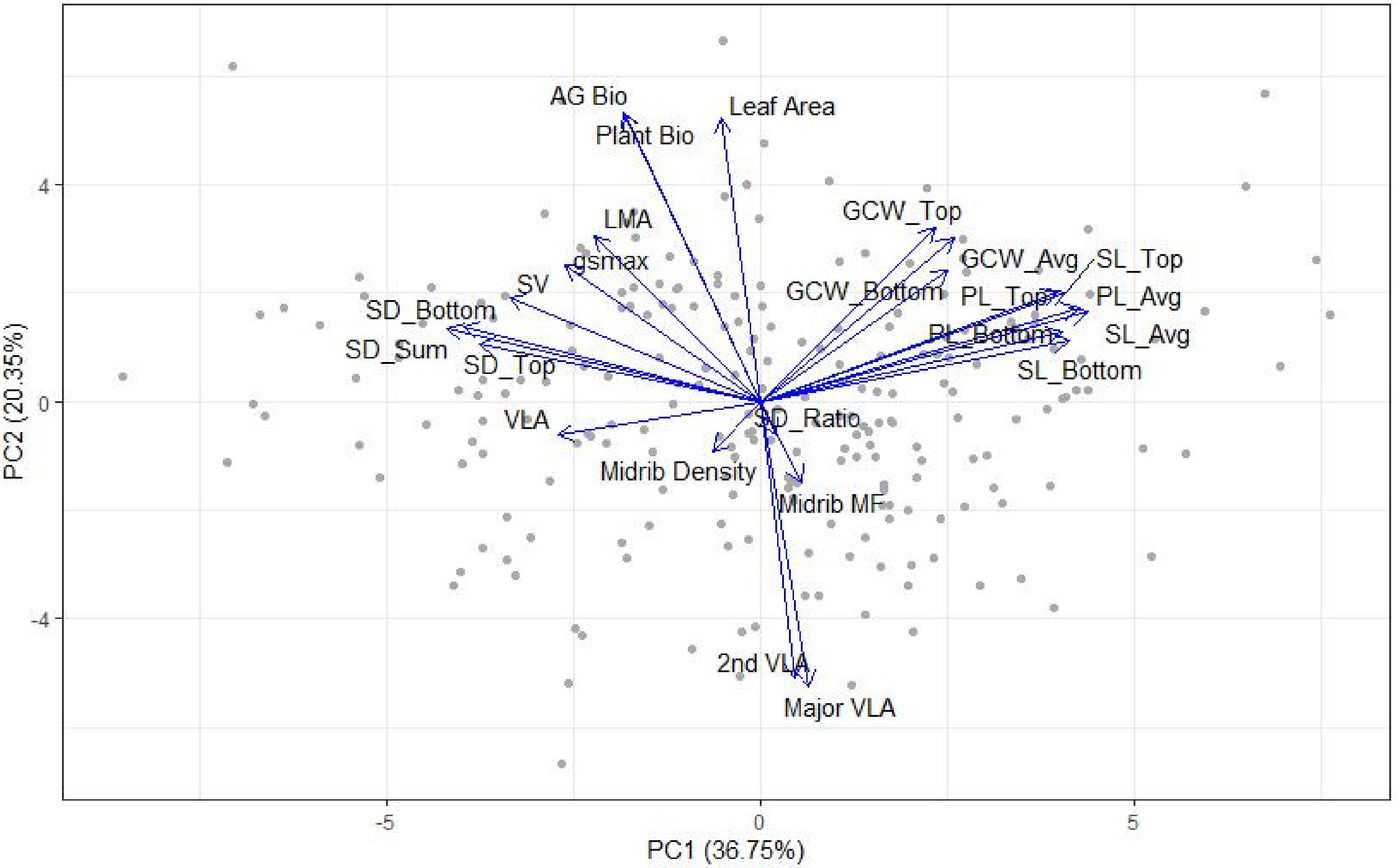
AG Bio
Leaf Area
Plant Bio
LMA
gsmax
SV
GCW_Top
GCW_Avg
SL_Top
GCW_Bottom
PL_Top
PL_Avg
PL_Bottom
SL_Avg
SL_Bottom
SD_Bottom
SD_Sum
SD_Top
VLA
SD_Ratio
Midrib Density
Midrib MF
2nd VLA
Major VLA
4
0
-4
-5
0
5
PC1 (36.75%)
PC2 (20.35%)

**Figure S6:** *Scree plot showing principal components vs. percent variance explained.*

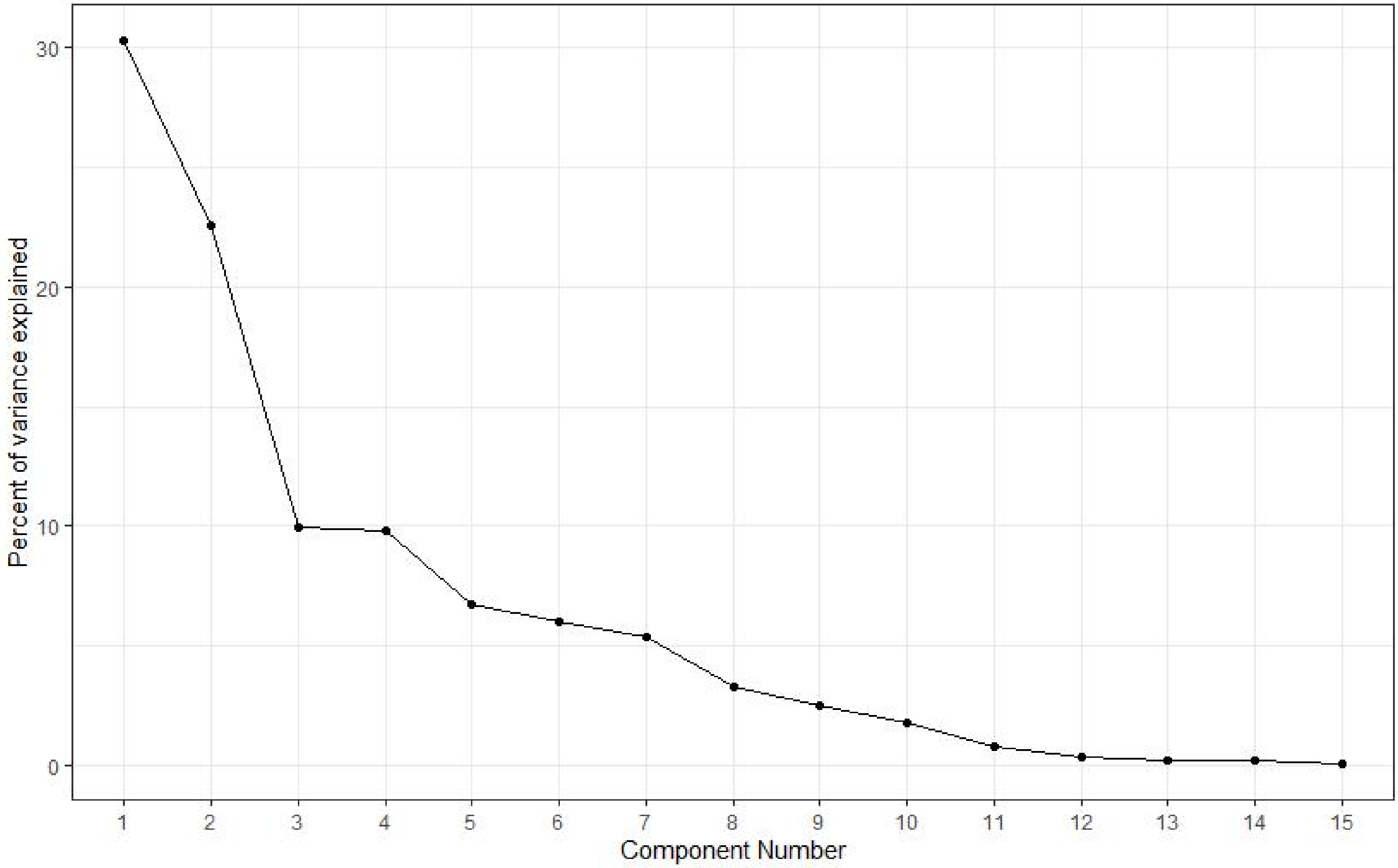

Percent of variance explained
30
20
10
Component Number

**Figure S7.** *Linkage disequilibrium heatmaps for all significant SNPs found on each chromosome. Linkage disequilibrium was estimated as* $R^2$ *between all significant SNPs per chromosome. Colored blocks represent haplotypic blocks that the SNPs belong to based on the full haplotype map (Figure S10). Block membership is shown for both an analysis based on the full, genome-wide collection of SNPs ("genome") as well as a re-analysis based on only significant SNPs ("significant"). Colors are arbitrary. For full methodological details, see Temme et al. (2020).*

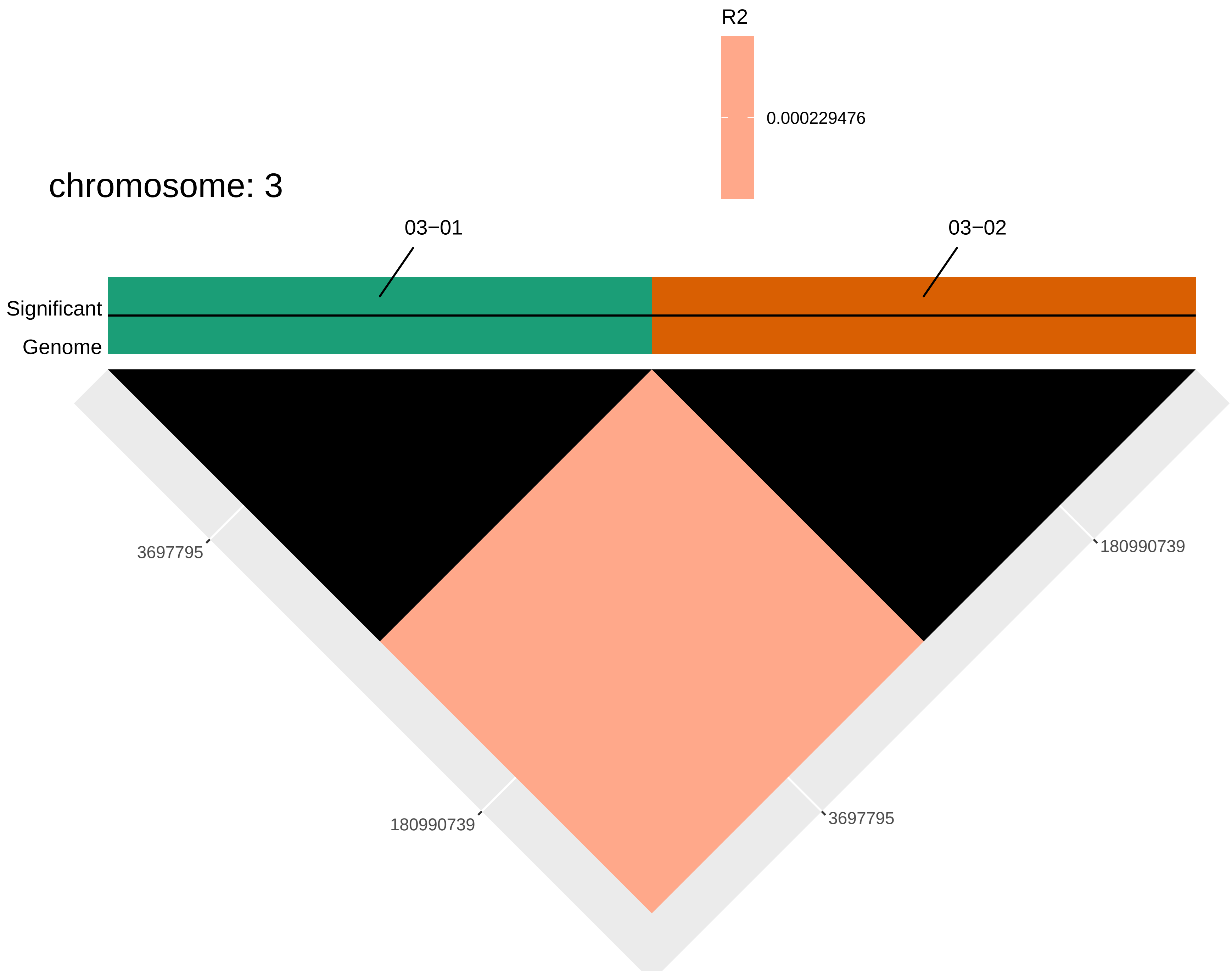

R2
0.000229476
chromosome: 3
03−01
03−02
Significant
Genome
3697795
180990739
180990739
3697795

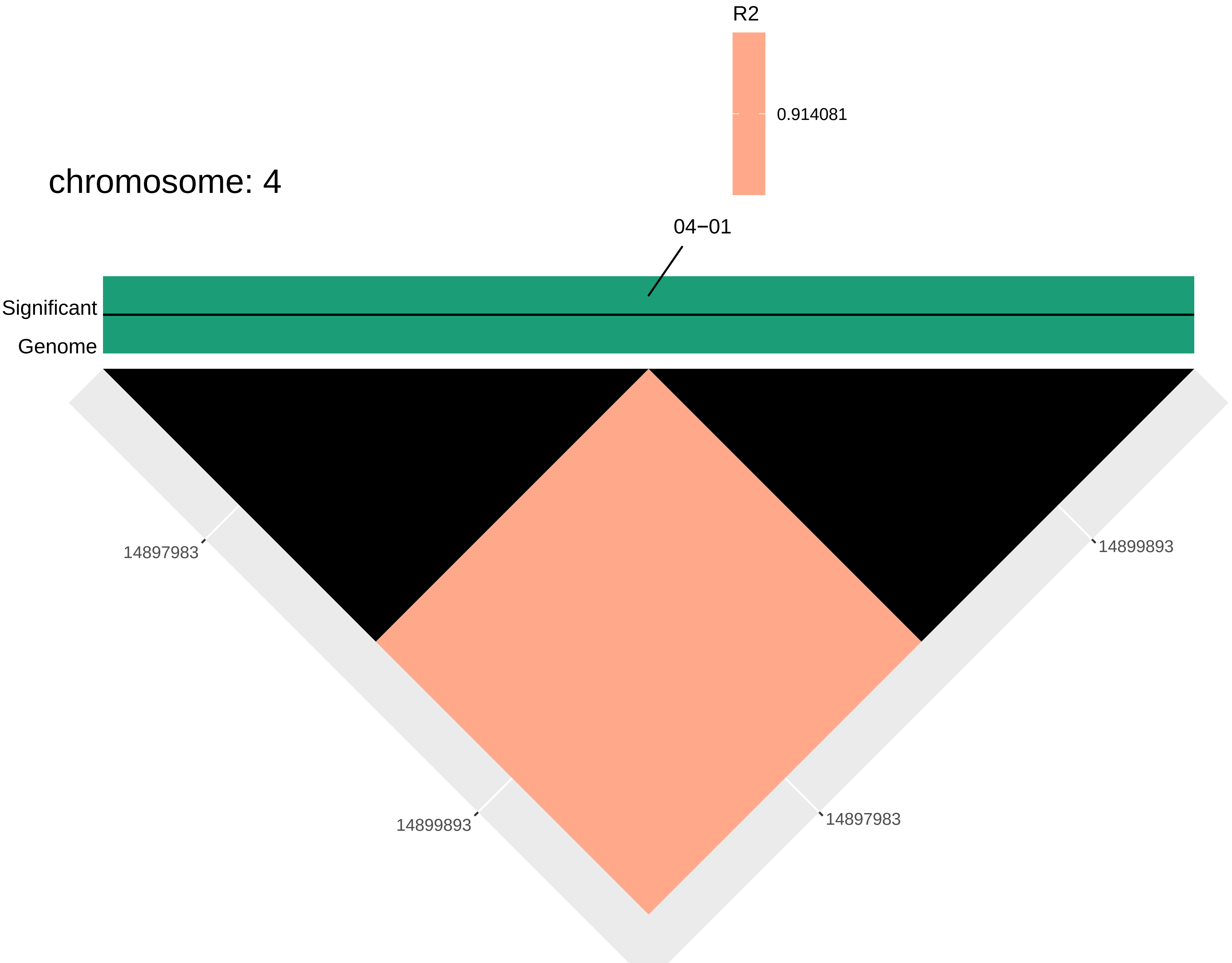

R2
0.914081
chromosome: 4
04−01
Significant
Genome
14897983
14899893
14899893
14897983

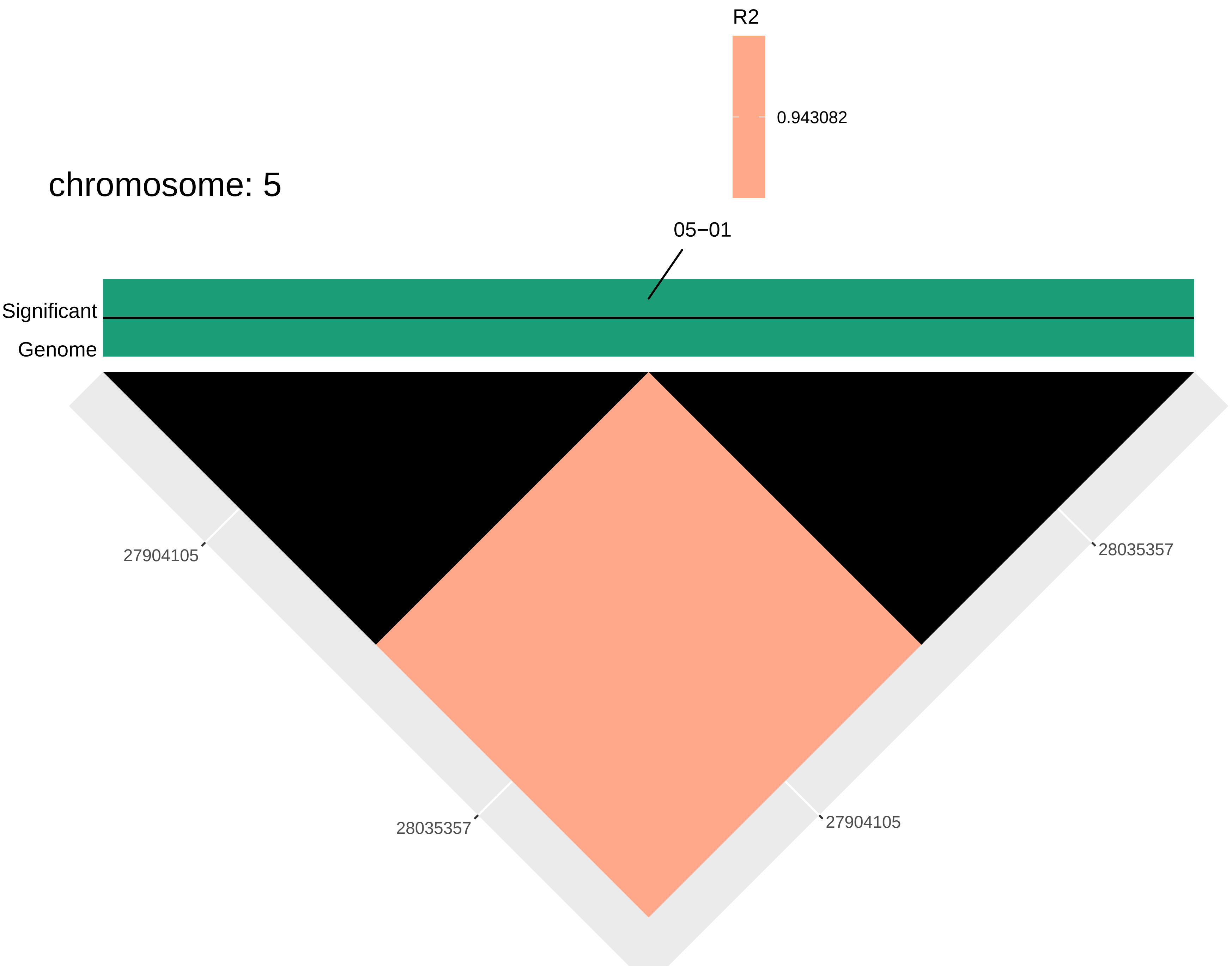

R2
0.943082
chromosome: 5
05−01
Significant
Genome
27904105
28035357
28035357
27904105

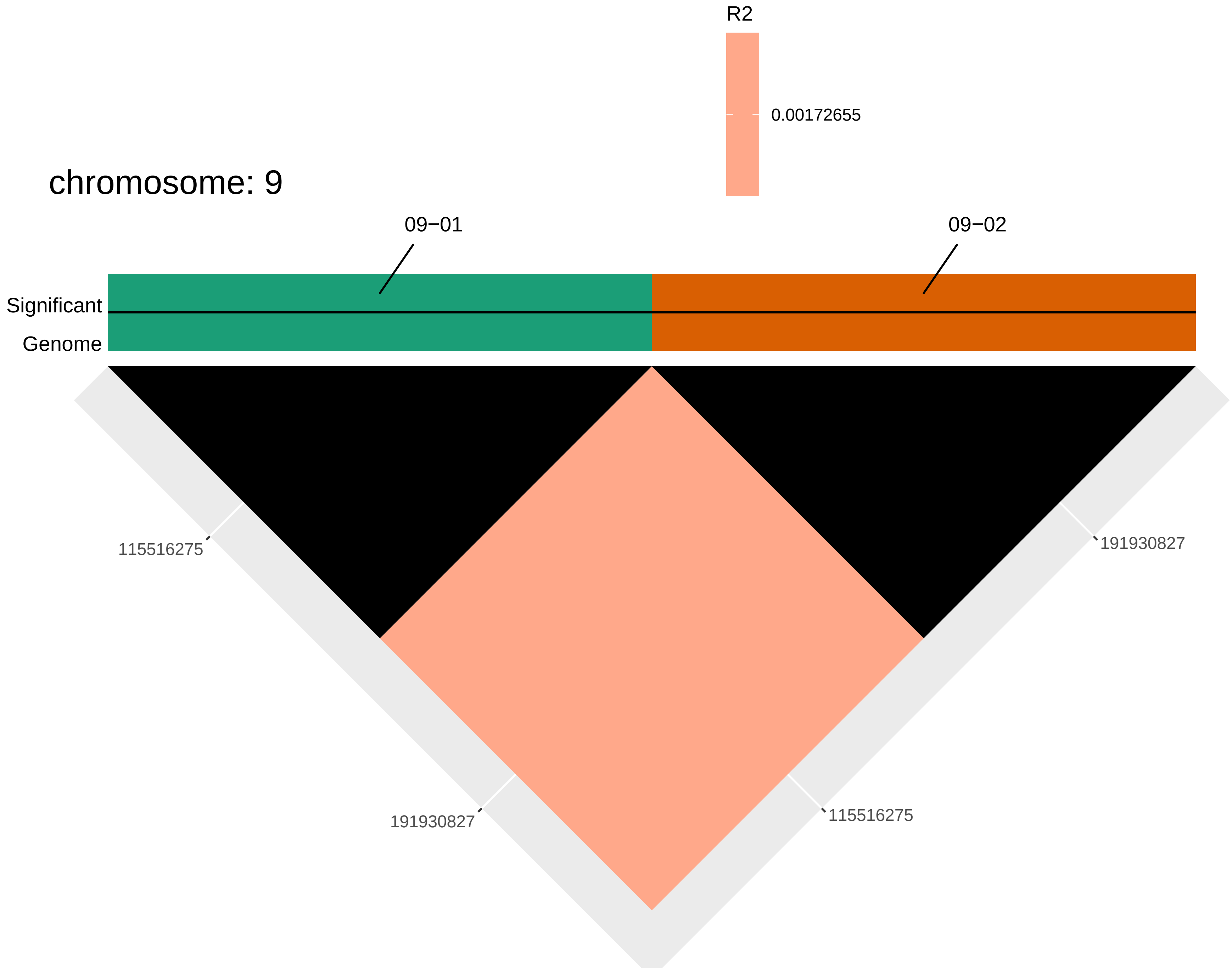
R2
0.00172655
chromosome: 9
09−01
09−02
Significant
Genome
115516275
191930827
191930827
115516275

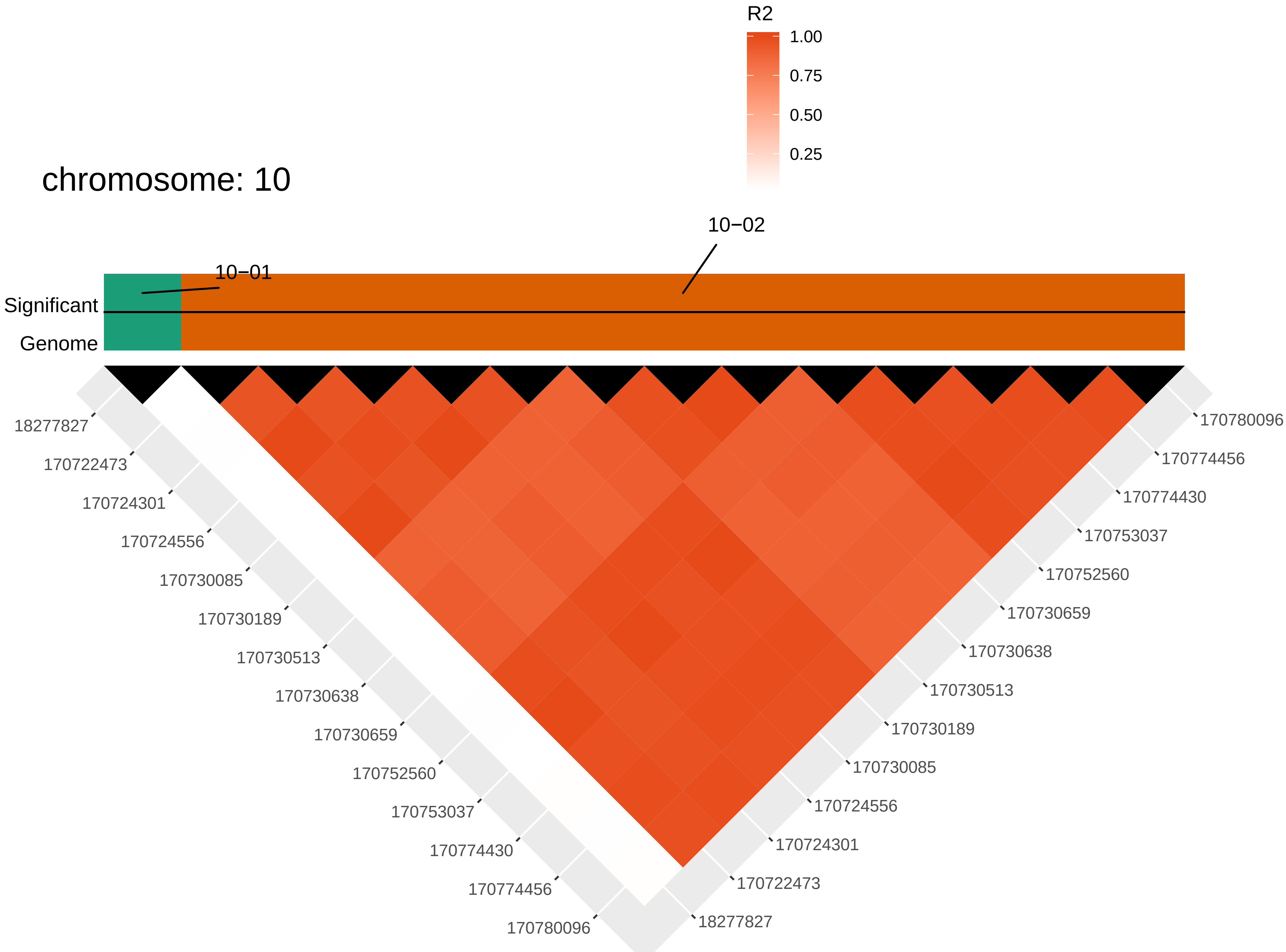

R2
1.00
0.75
0.50
0.25
chromosome: 10
10−02
10−01
Significant
Genome
18277827
170722473
170724301
170724556
170730085
170730189
170730513
170730638
170730659
170752560
170753037
170774430
170774456
170780096

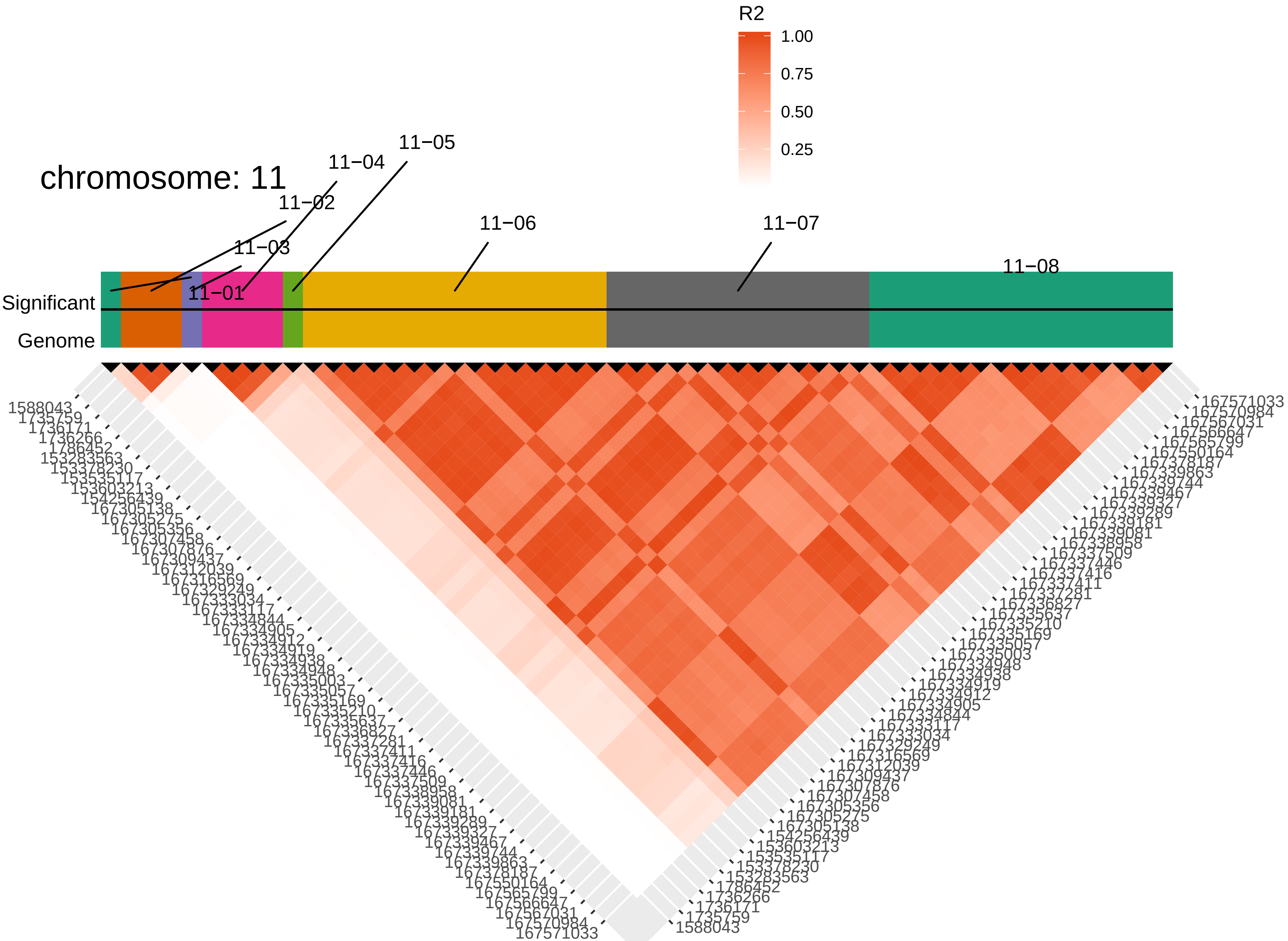

chromosome: 11
R2
1.00
0.75
0.50
0.25
11−01
11−02
11−03
11−04
11−05
11−06
11−07
11−08
Significant
Genome
1588043
1735759
1736171
1736266
1786452
153283563
153378230
153535117
153603213
154256439
167305138
167305275
167305356
167307458
167307876
167309437
167312039
167316569
167329249
167333034
167333117
167334844
167334905
167334912
167334919
167334938
167334948
167335003
167335057
167335169
167335210
167335637
167336827
167337281
167337411
167337416
167337446
167337509
167338958
167339081
167339181
167339289
167339327
167339467
167339744
167339863
167378187
167550164
167565799
167566647
167567031
167570984
167571033

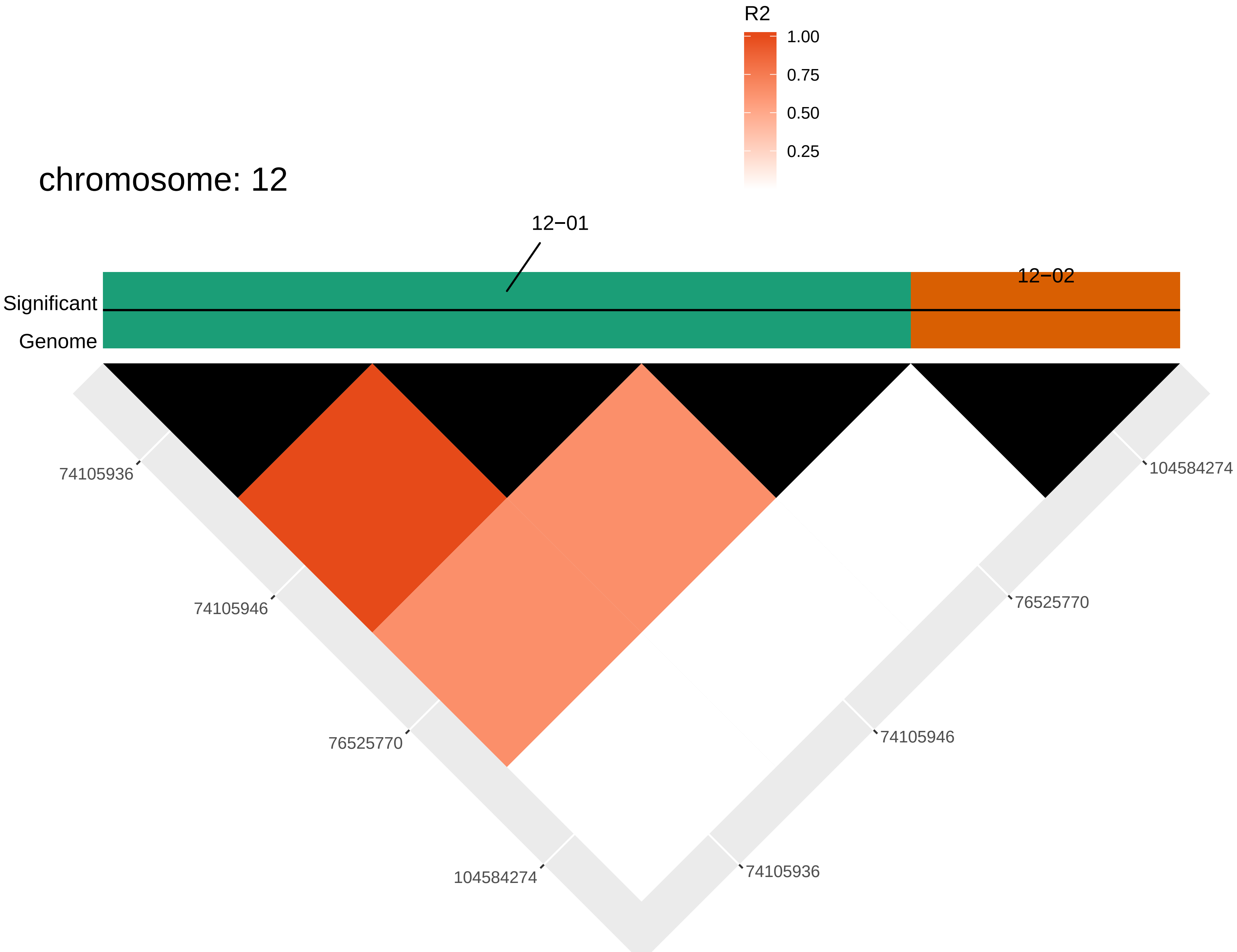

R2
1.00
0.75
0.50
0.25
chromosome: 12
12−01
12−02
Significant
Genome
74105936
74105946
76525770
104584274
104584274
76525770
74105946
74105936

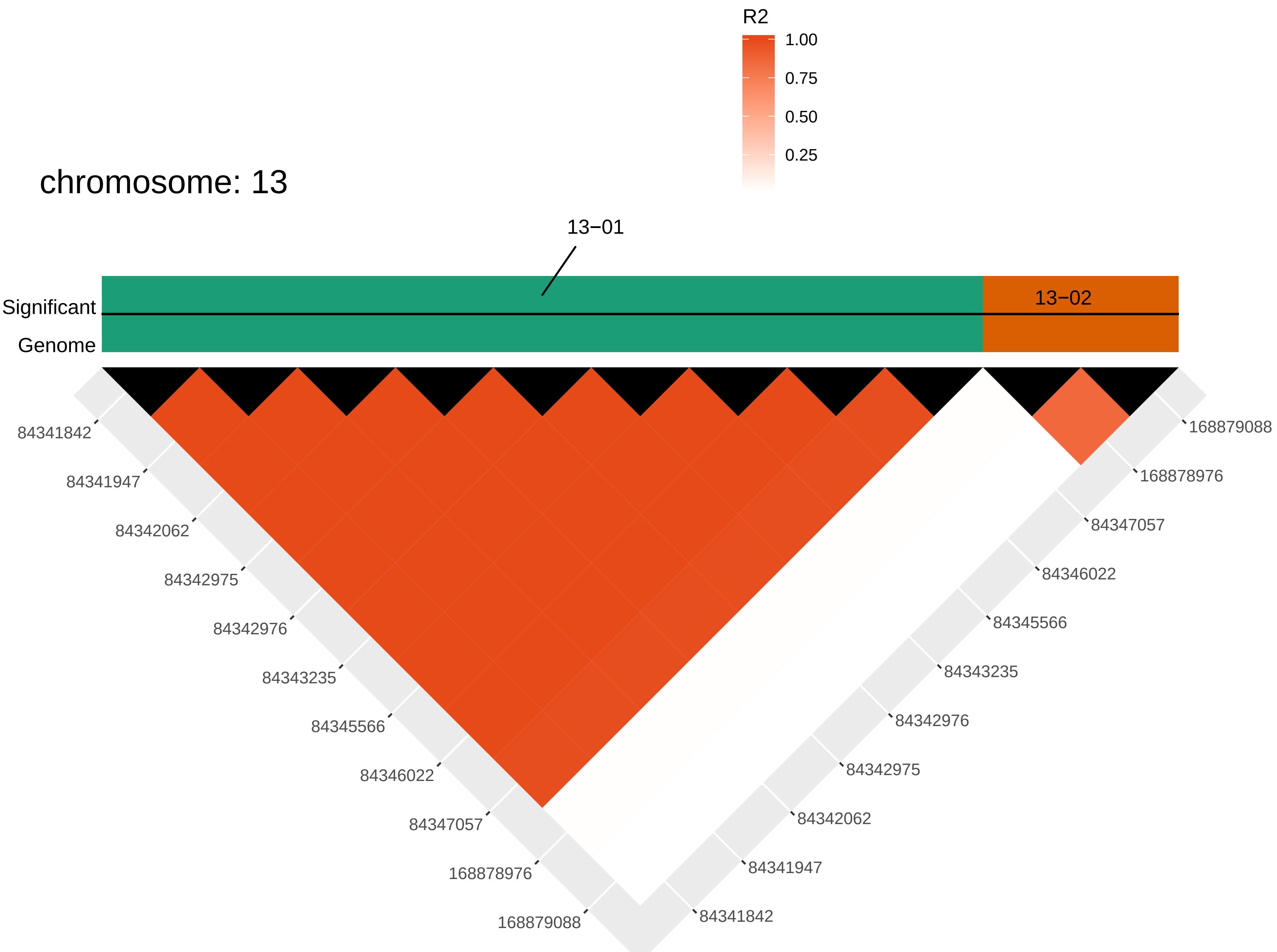
R2
1.00
0.75
0.50
0.25
chromosome: 13
13−01
13−02
Significant
Genome
84341842
84341947
84342062
84342975
84342976
84343235
84345566
84346022
84347057
168878976
168879088

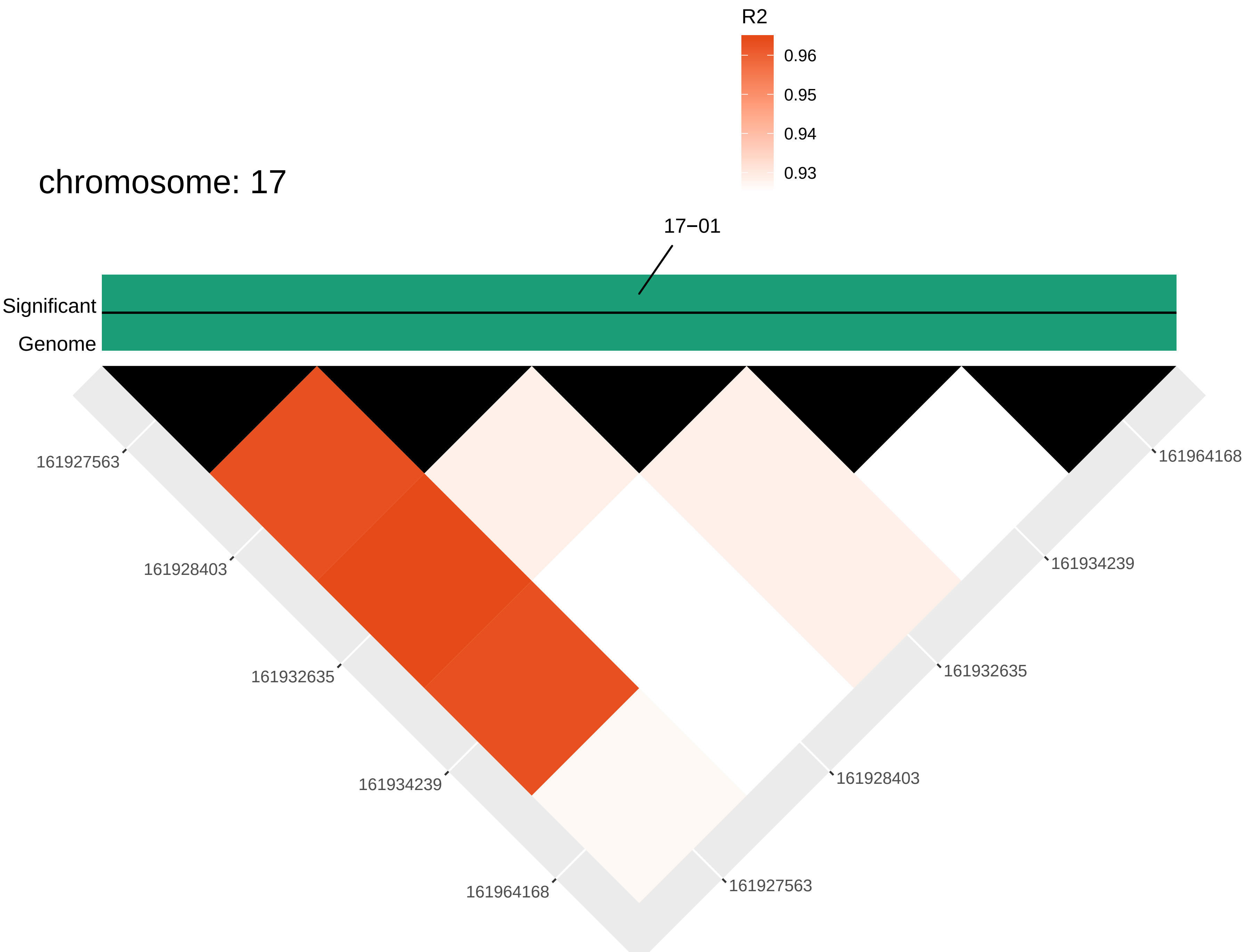

R2
0.96
0.95
0.94
0.93
chromosome: 17
17−01
Significant
Genome
161927563
161928403
161932635
161934239
161964168
161964168
161934239
161932635
161928403
161927563

**Figure S8:** *Manhattan plots resulting from GWA analyses for all traits. The red line is the significance threshold based on the modified Bonferroni correction (see text for details) and the blue line is the suggestive threshold based on (i.e., top 0.1% of all SNPs). Colored dots represent SNPs that are significant or suggestive for at least one trait. Color of dots is arbitrary.*

SD bottom

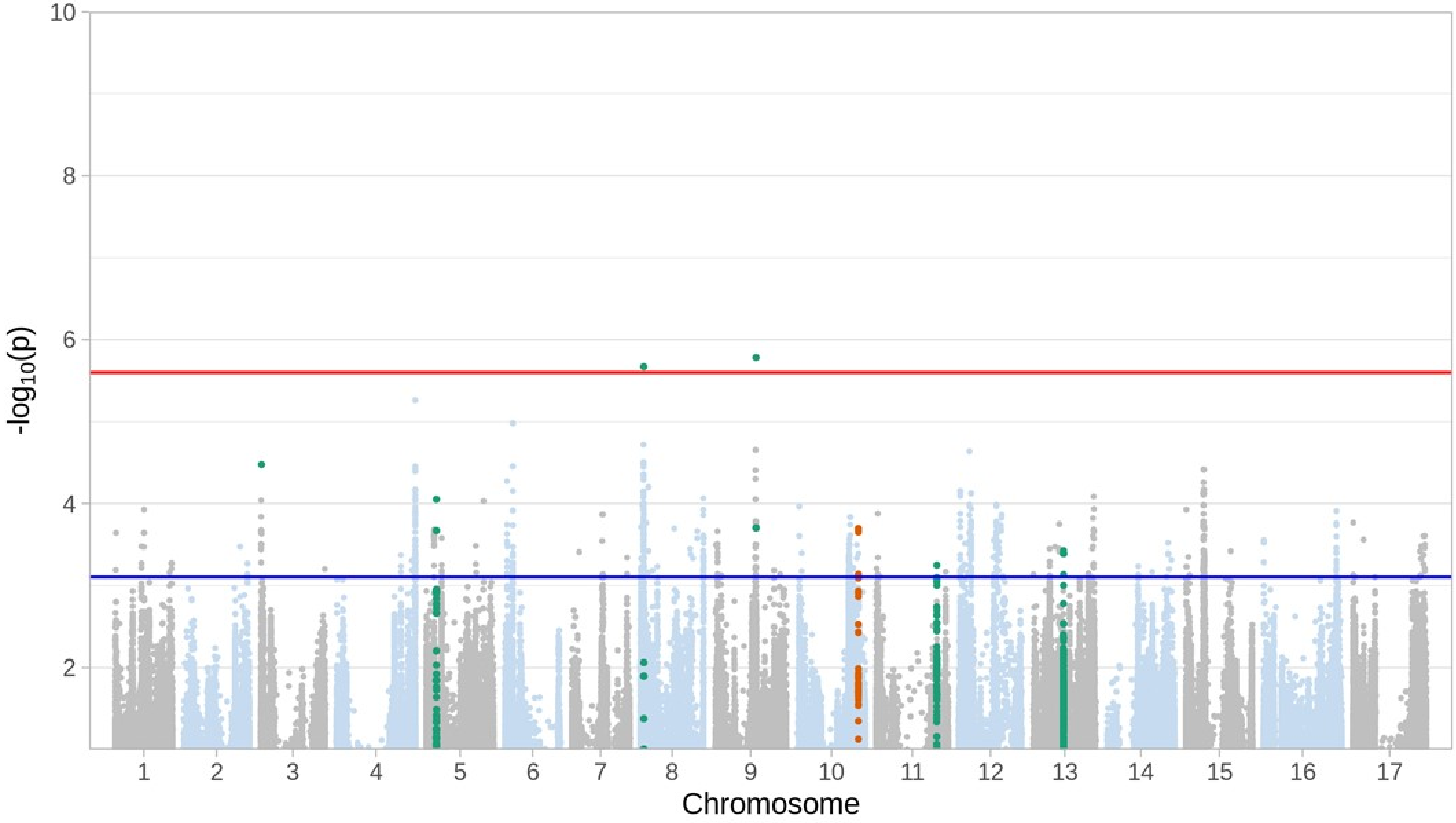

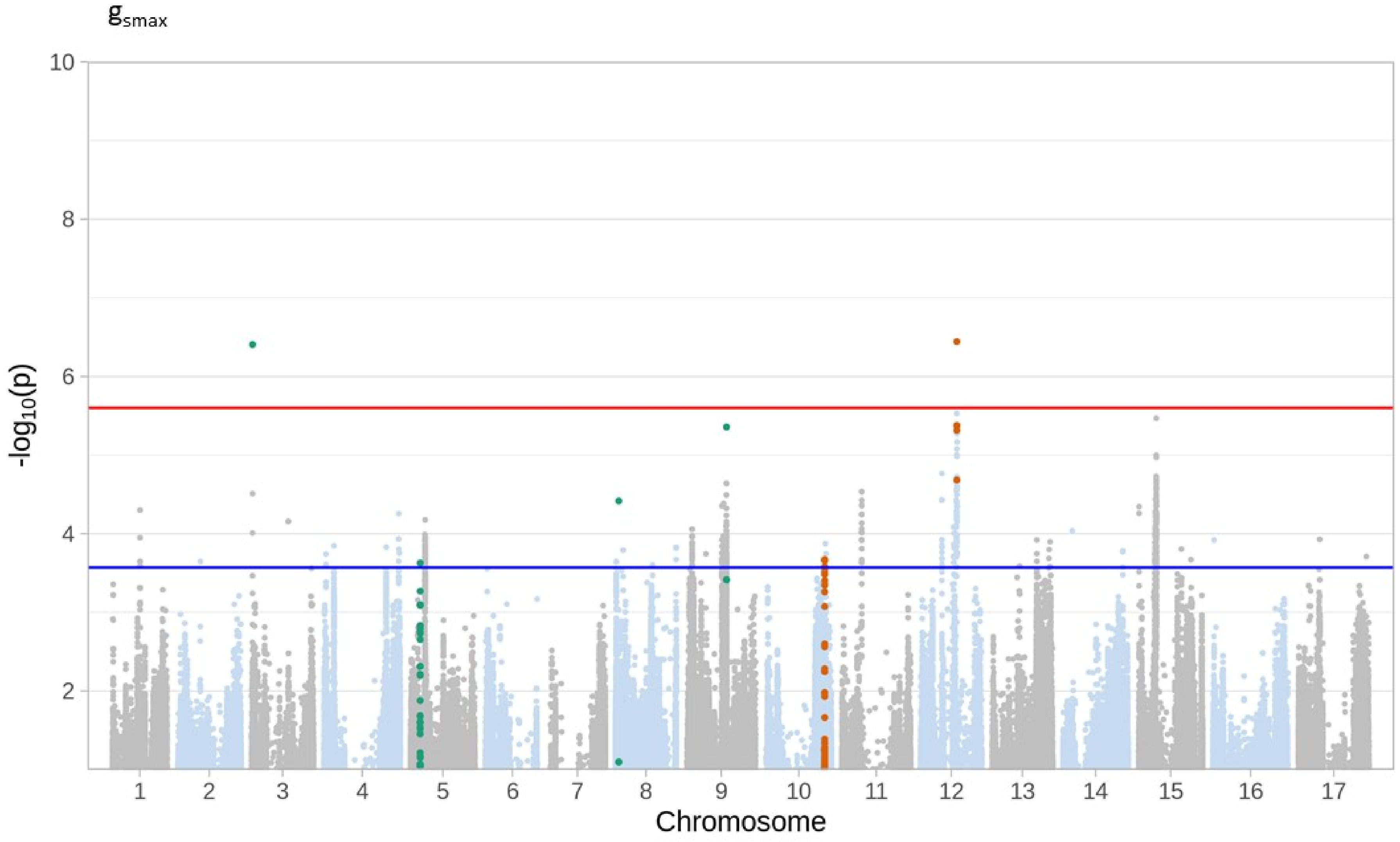

gsmax
10
8
6
4
2
-log10(p)
Chromosome

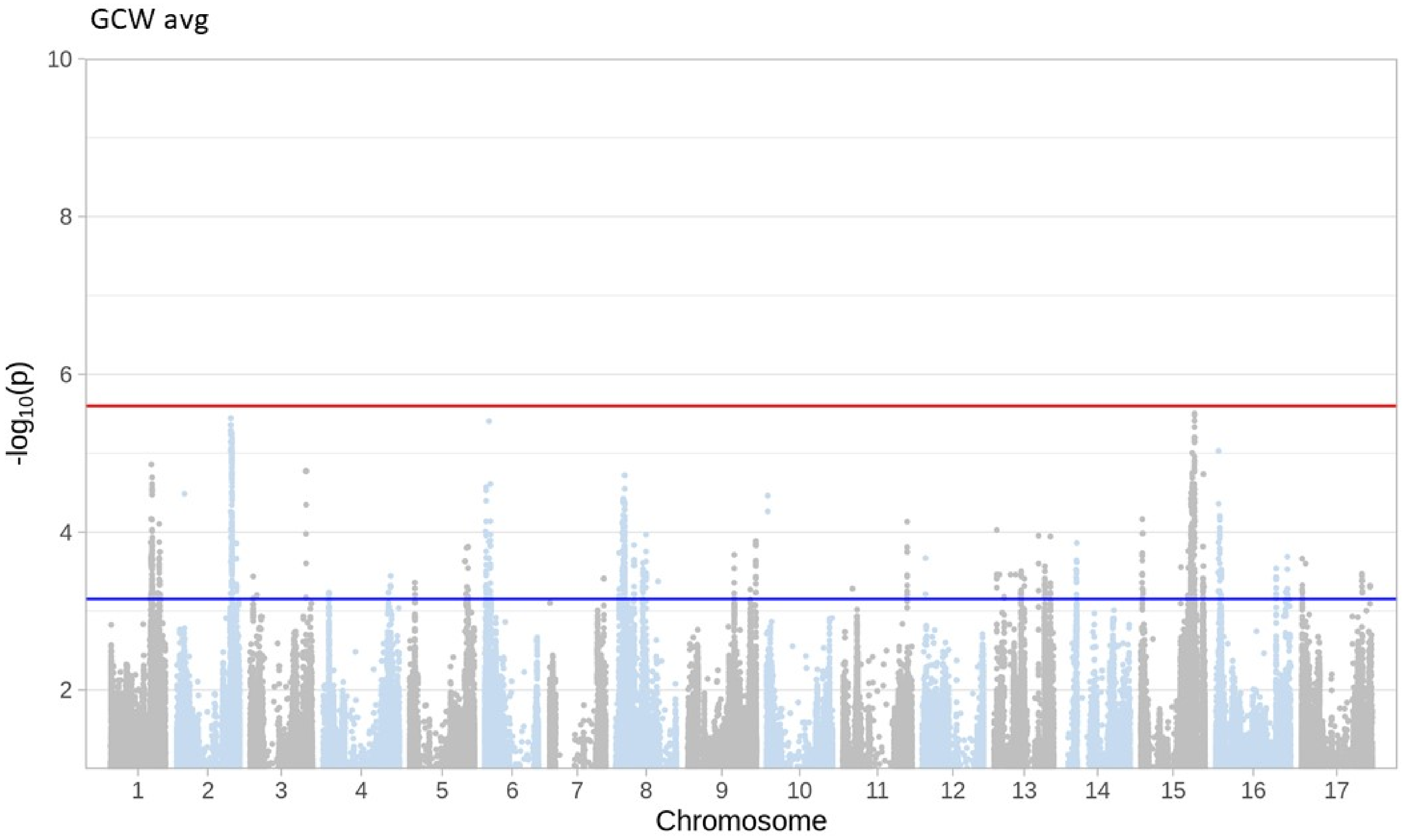
GCW avg
10
8
6
4
2
-log10(p)
Chromosome

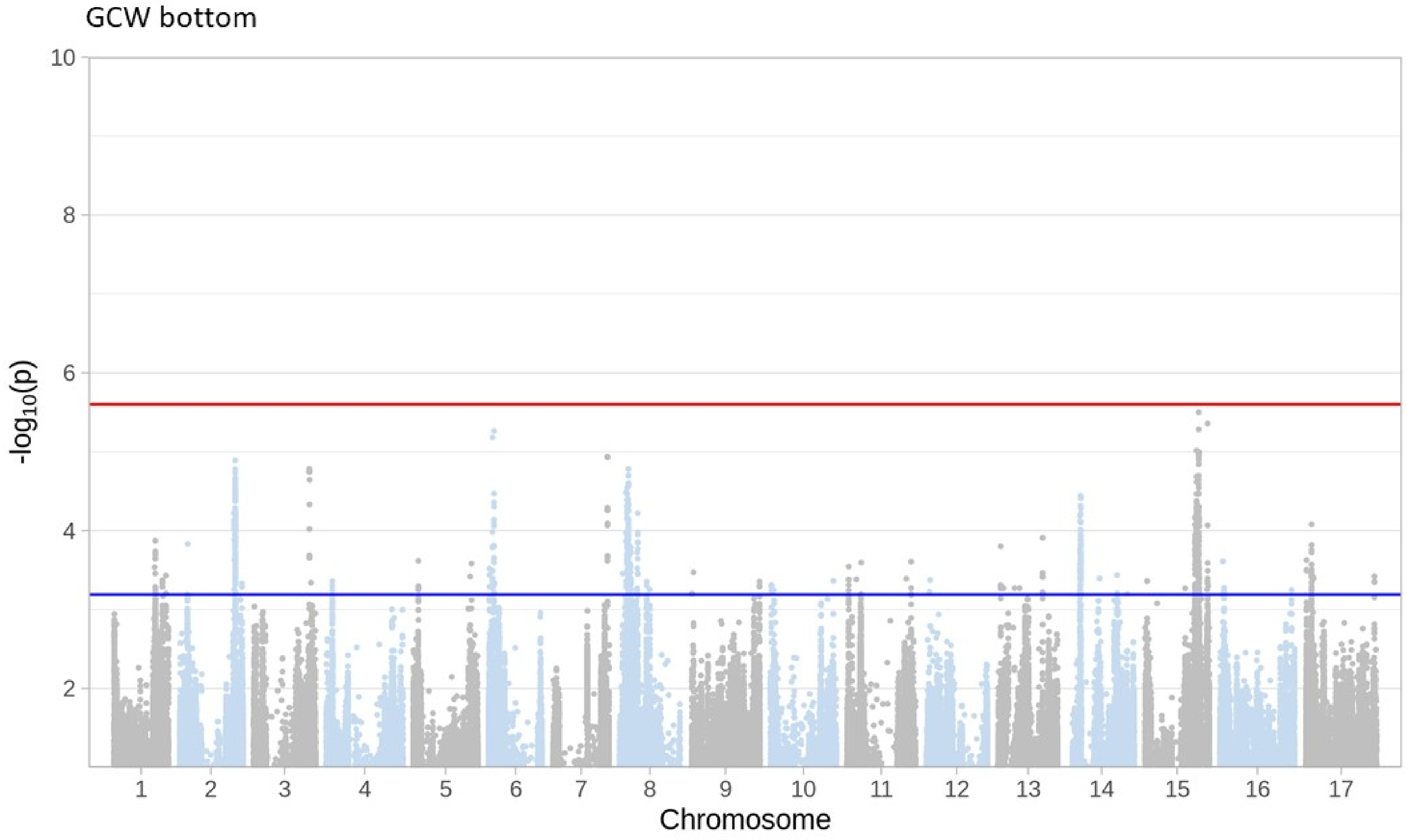
GCW bottom
10
8
6
4
2
$-\log_{10}(p)$
Chromosome

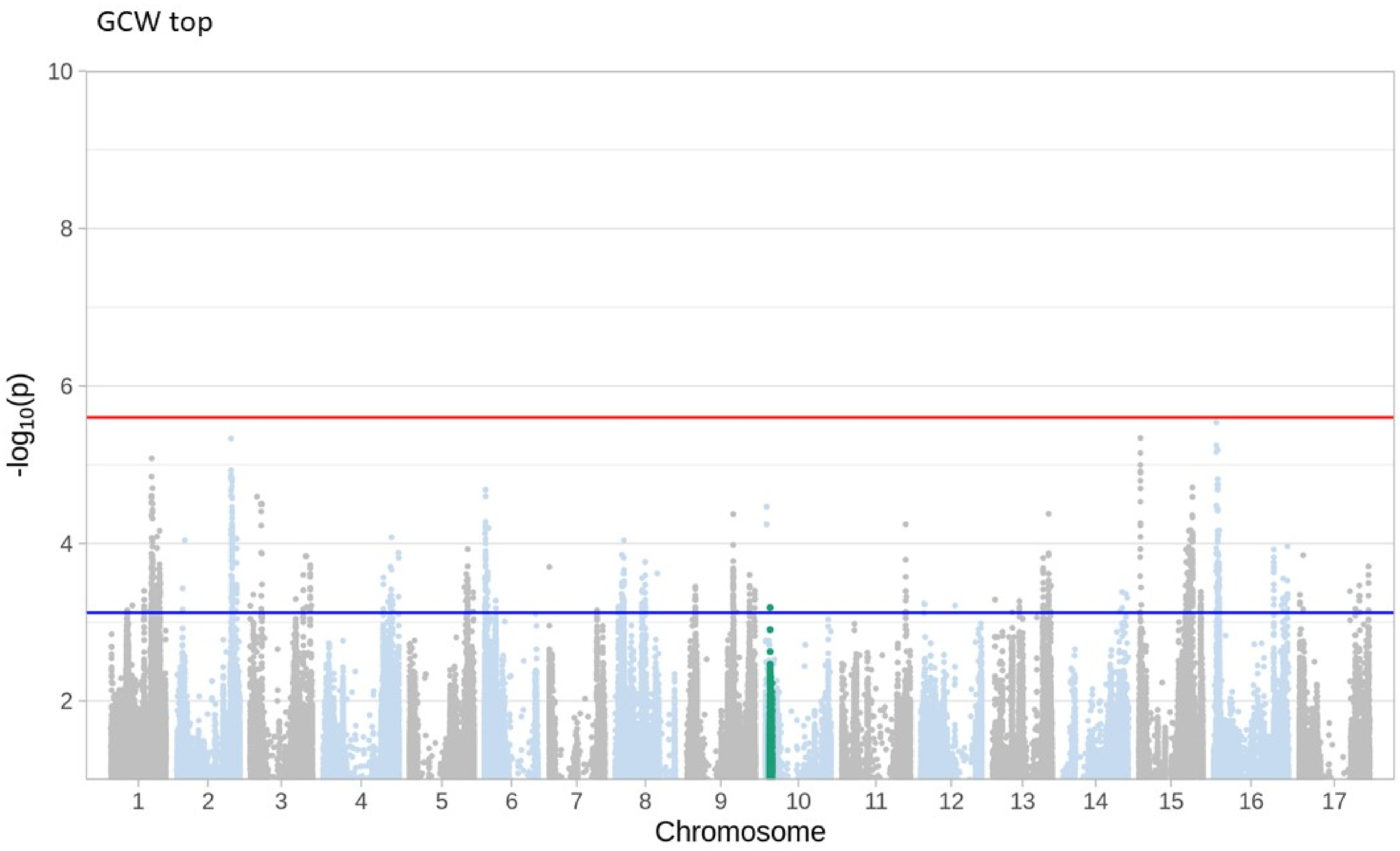
GCW top
10
8
6
4
2
-log10(p)
Chromosome

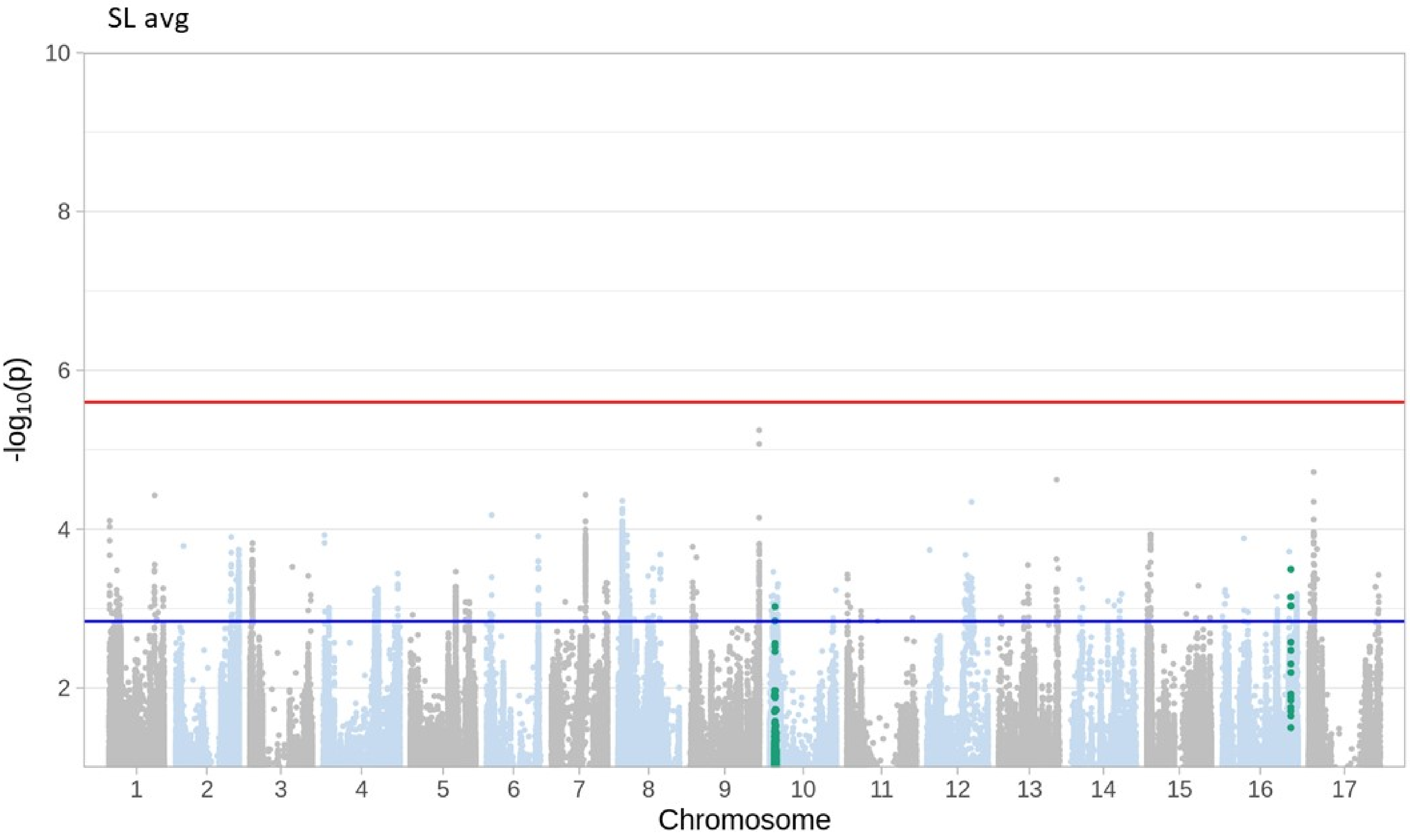

SL avg
-log10(p)
10
8
6
4
2
Chromosome

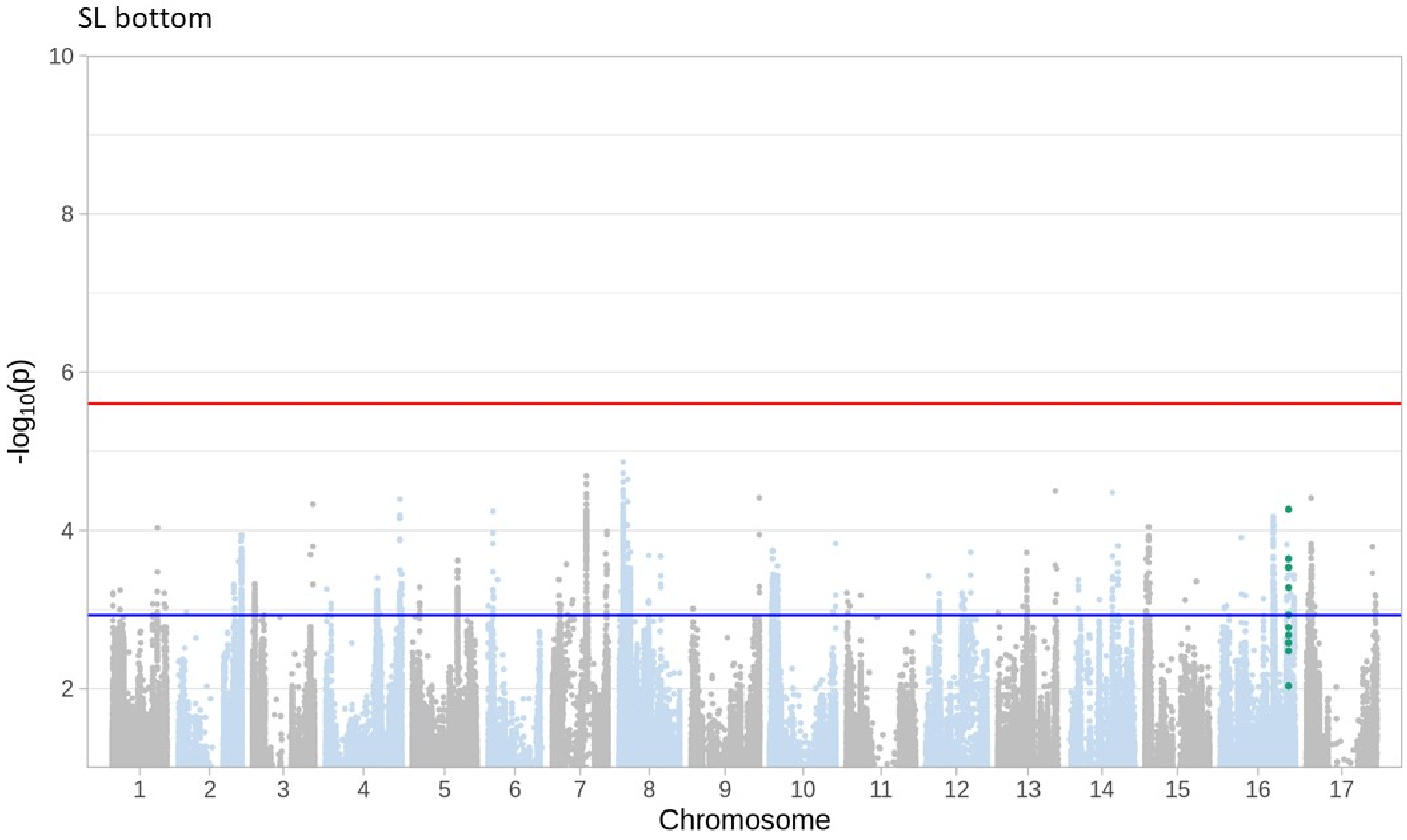
SL bottom
10
8
6
4
2
-log10(p)
Chromosome

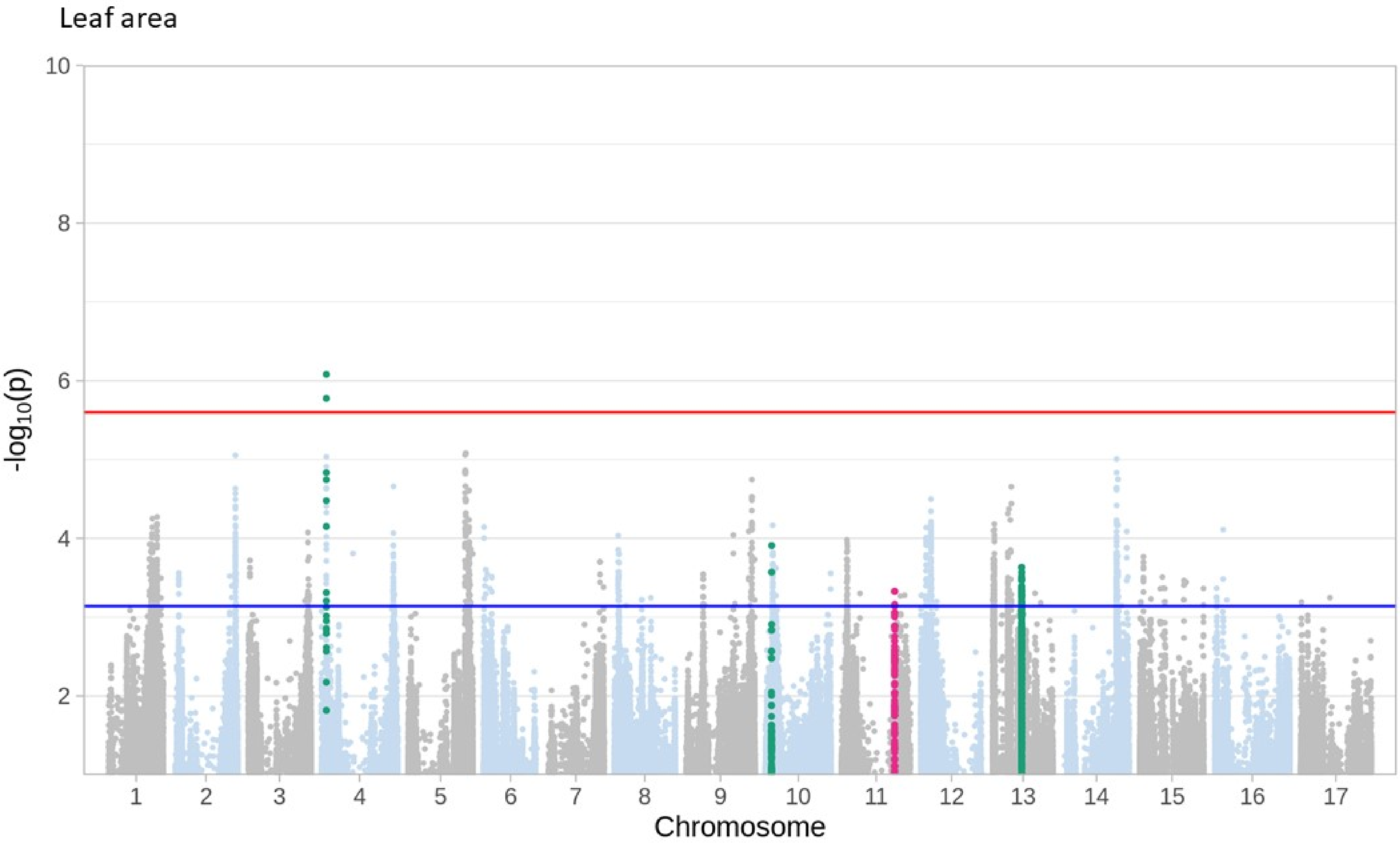

Leaf area
10
8
6
4
2
-log10(p)
Chromosome

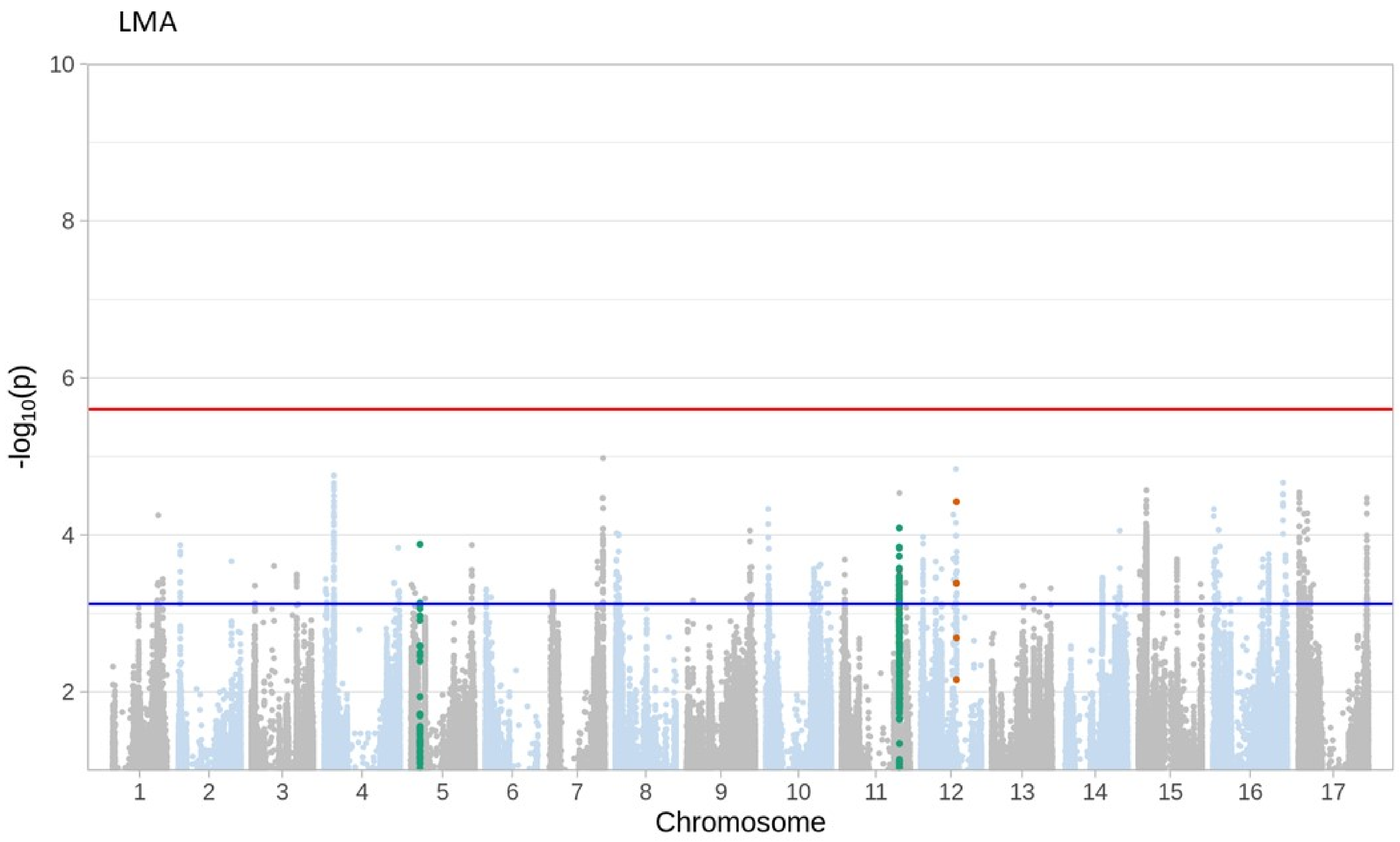
LMA
-log10(p)
10
8
6
4
2
1 2 3 4 5 6 7 8 9 10 11 12 13 14 15 16 17
Chromosome

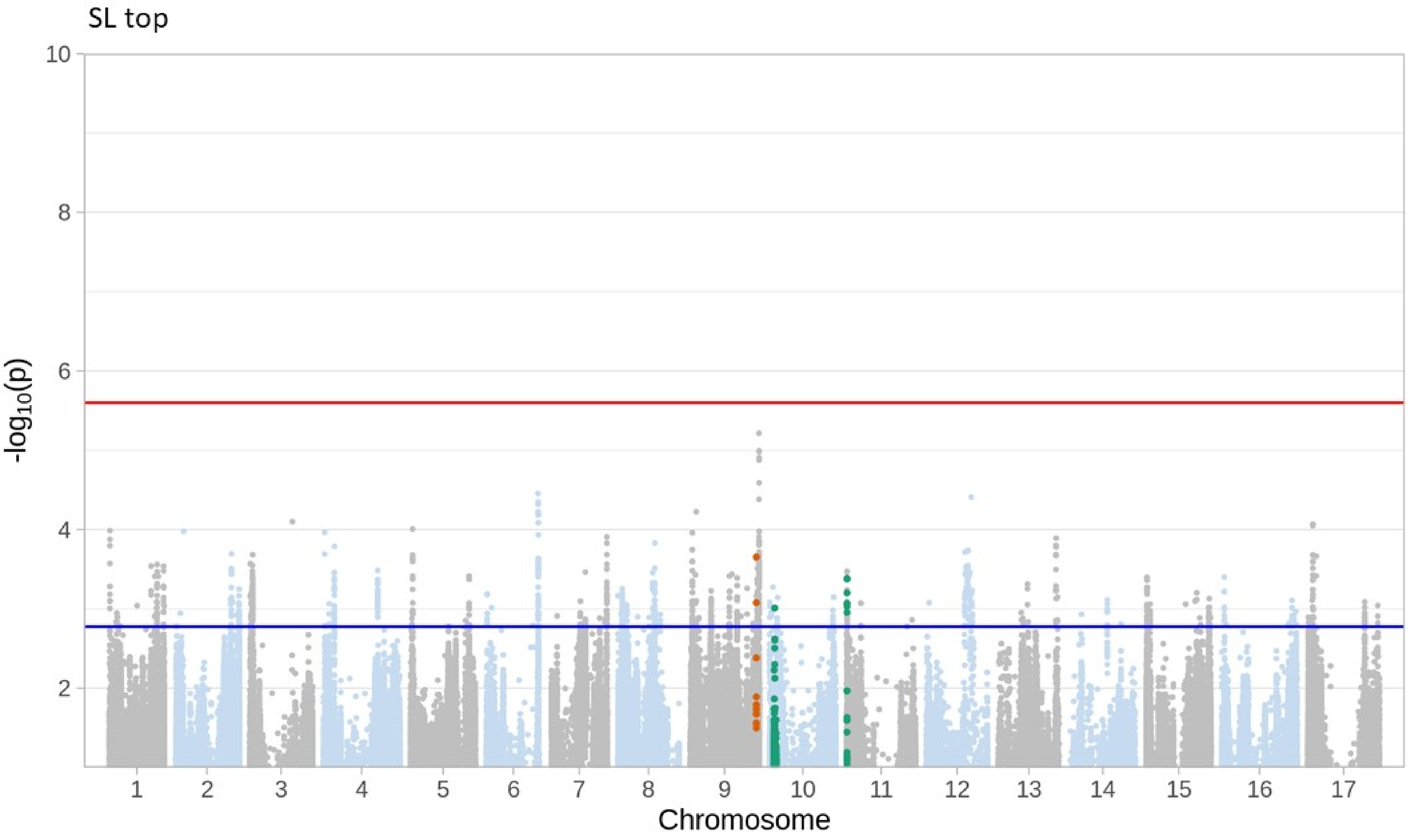
SL top
-log10(p)
10
8
6
4
2
Chromosome

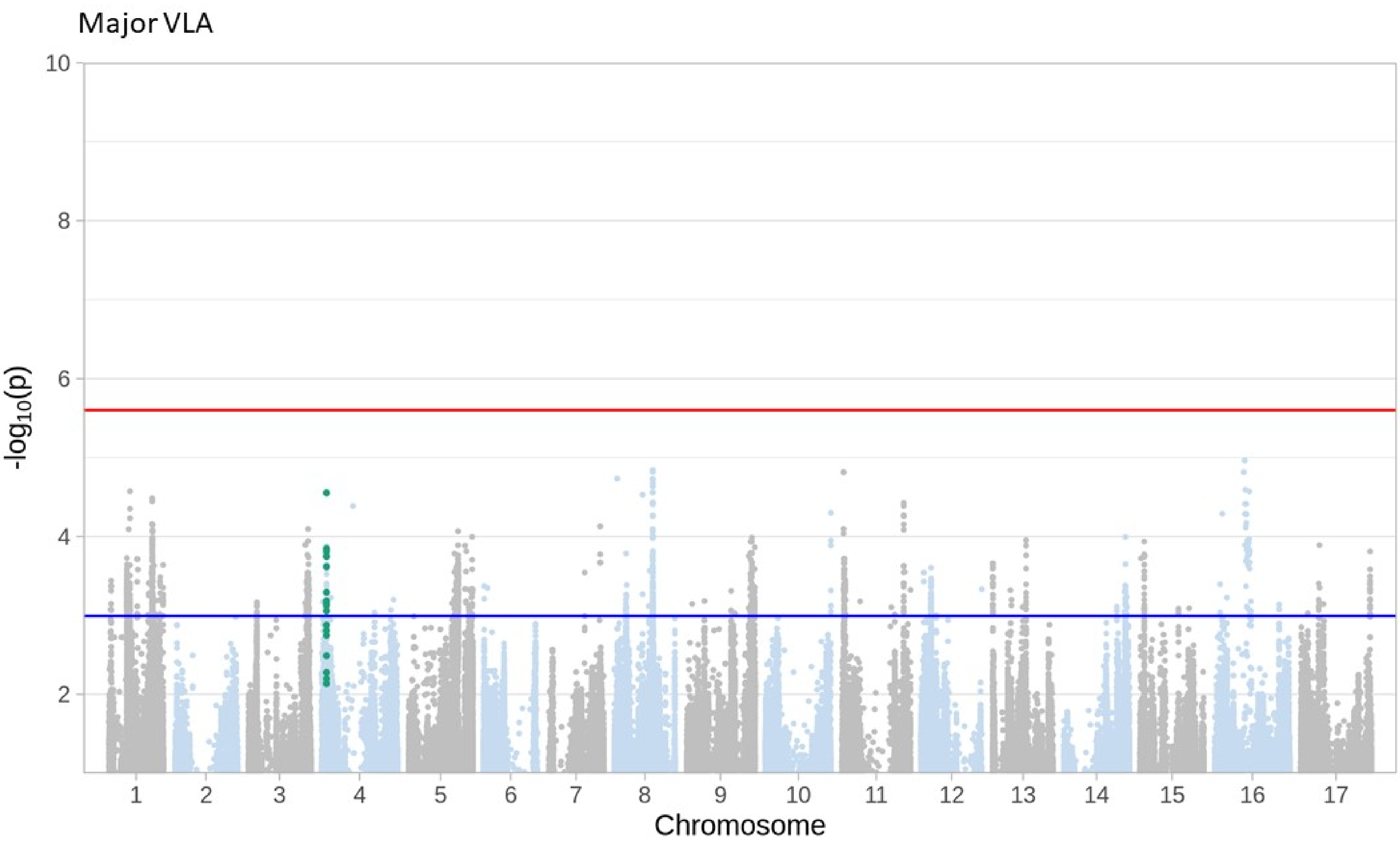
Major VLA
-log10(p)
10
8
6
4
2
Chromosome

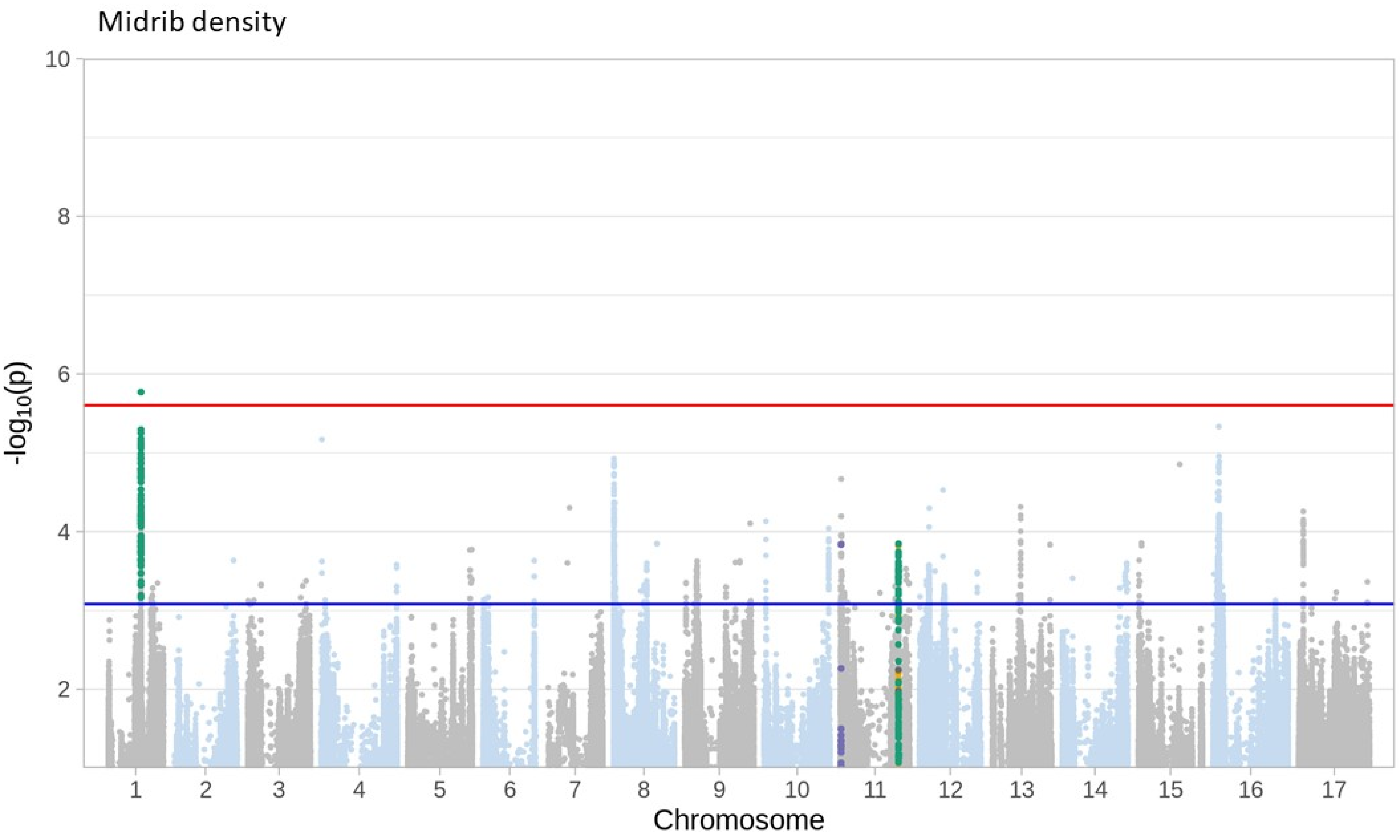

Midrib density
10
8
6
4
2
-log10(p)
Chromosome

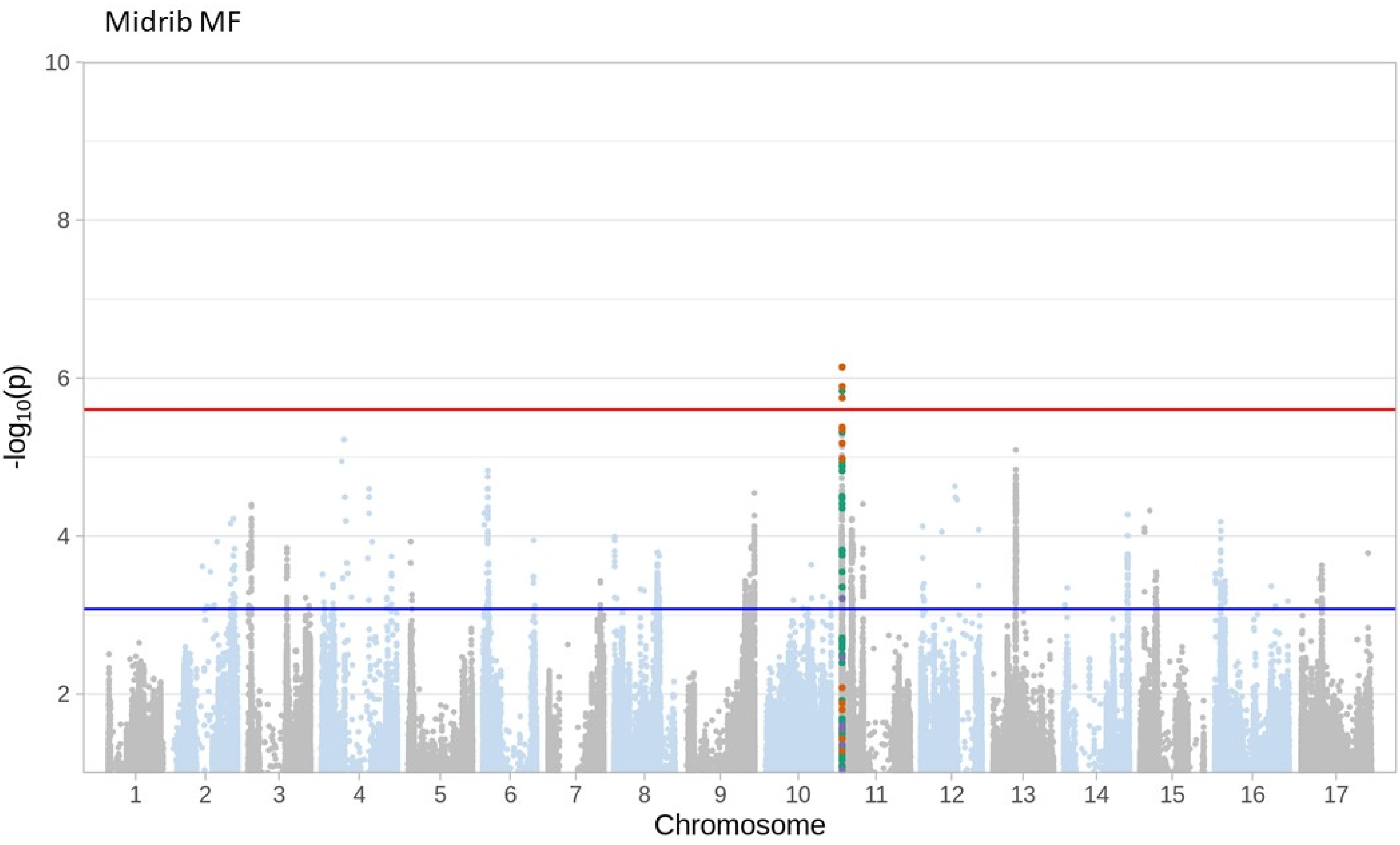
Midrib MF
-log10(p)
10
8
6
4
2
Chromosome

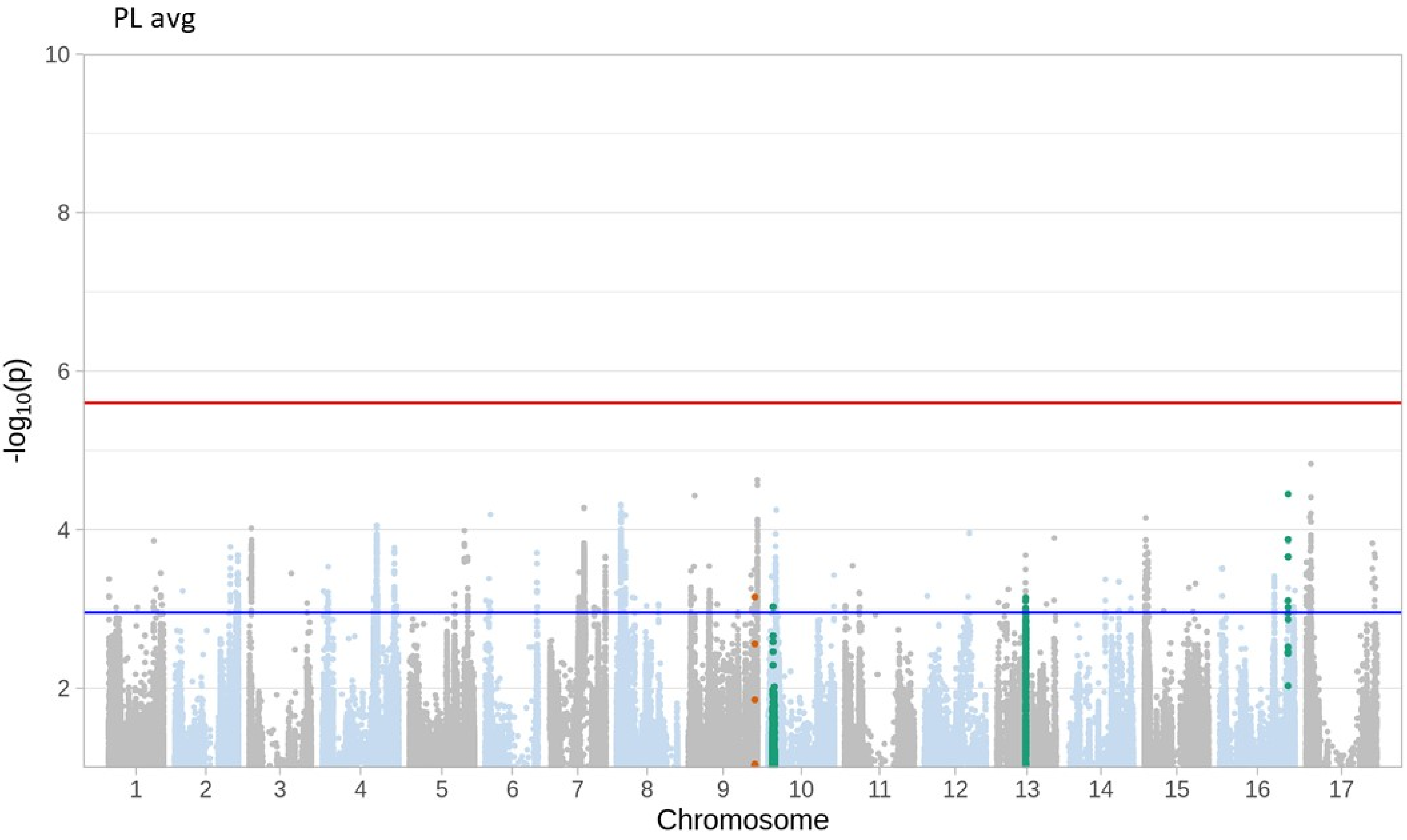
PL avg
10
8
6
4
2
-log10(p)
Chromosome

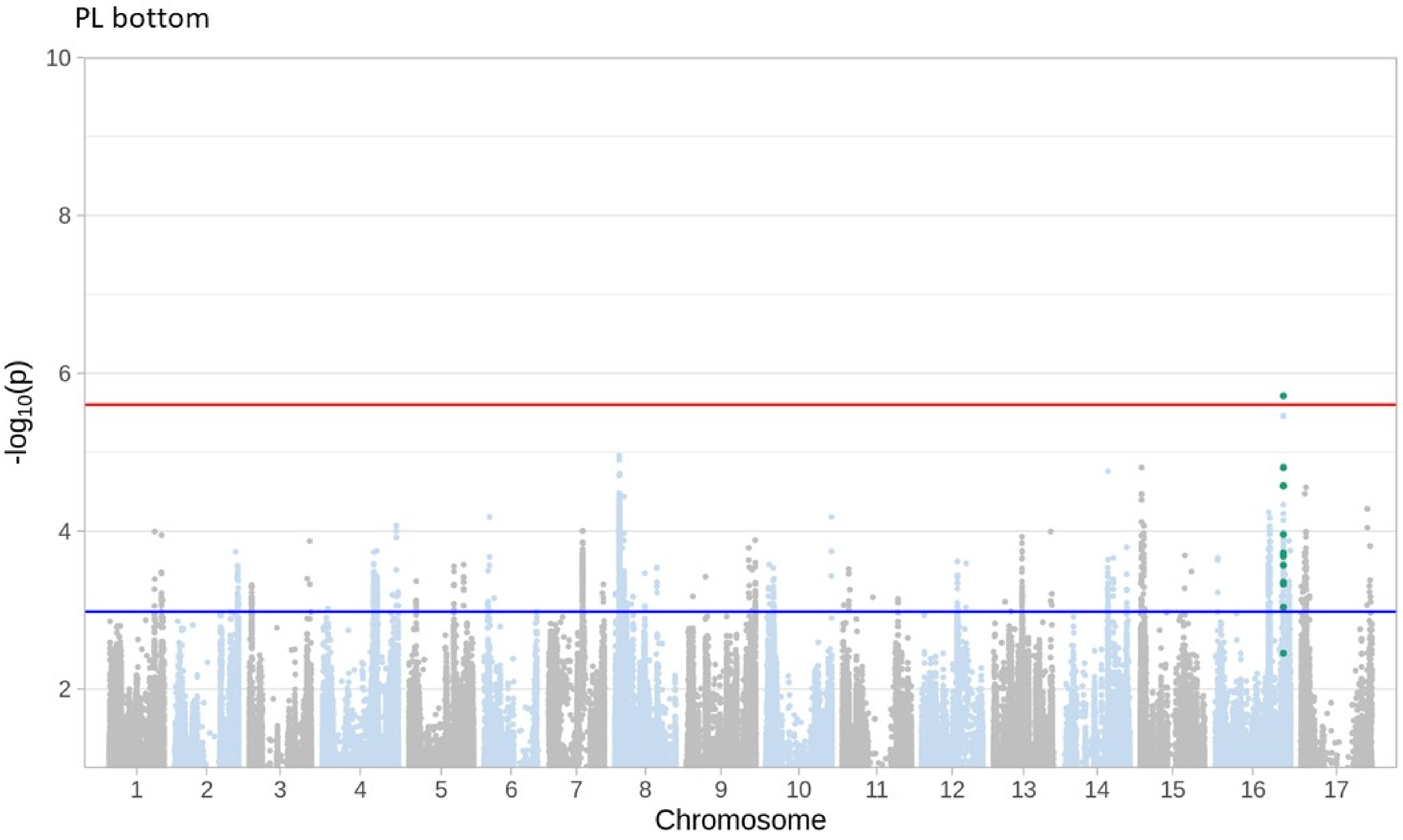
PL bottom
10
8
6
4
2
-log10(p)
Chromosome

PL top

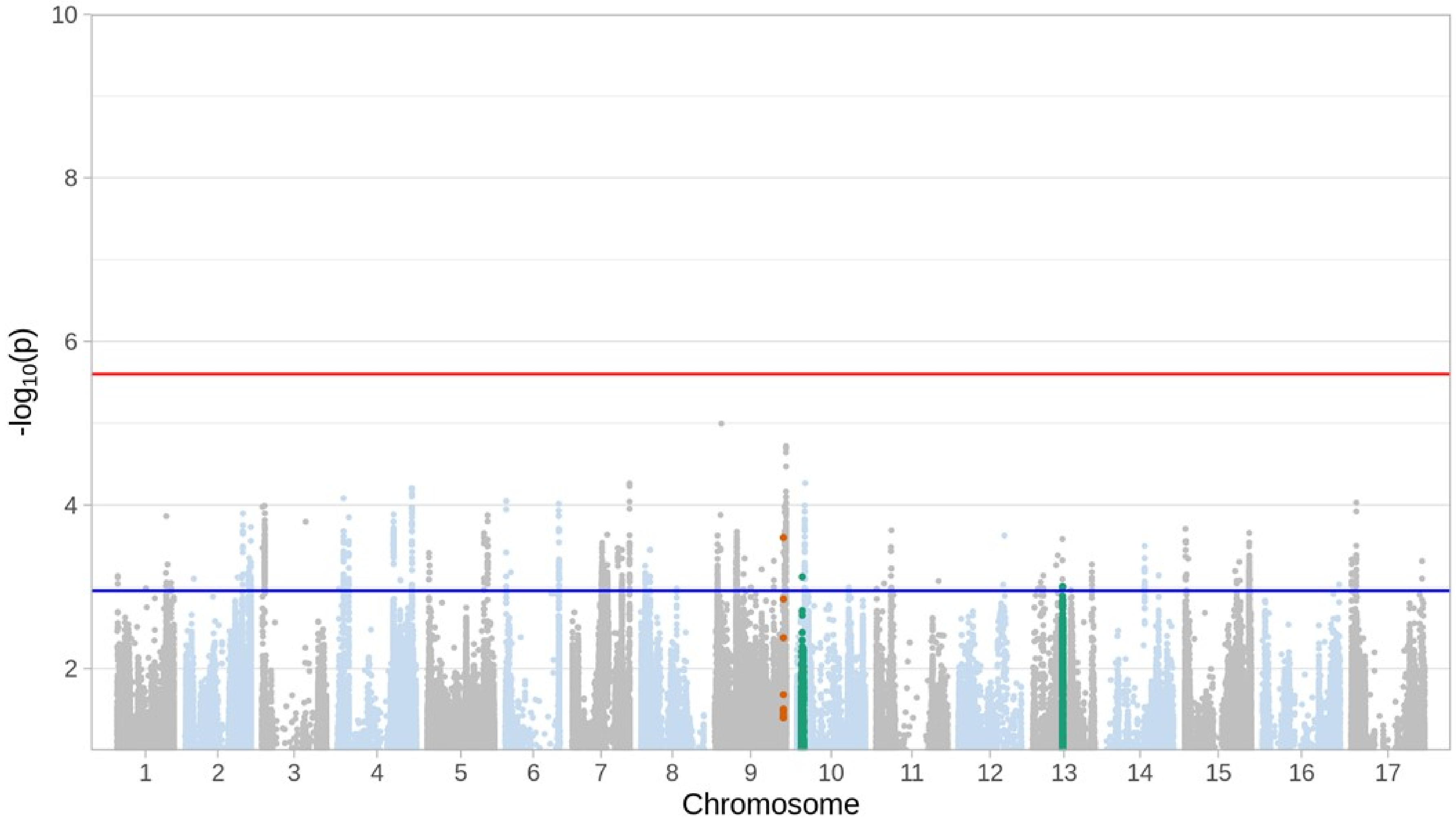

Stomatal ratio

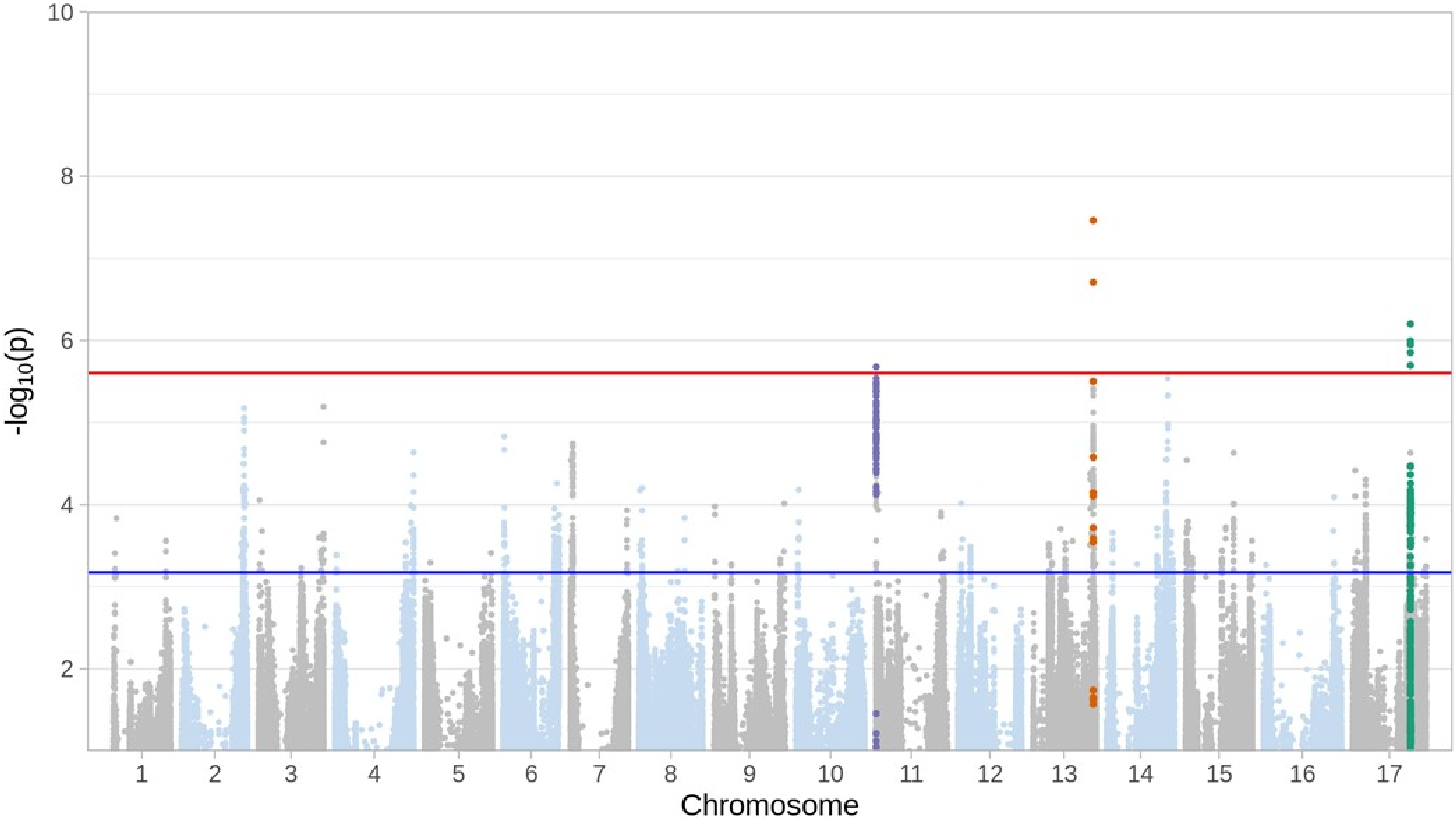

SD sum

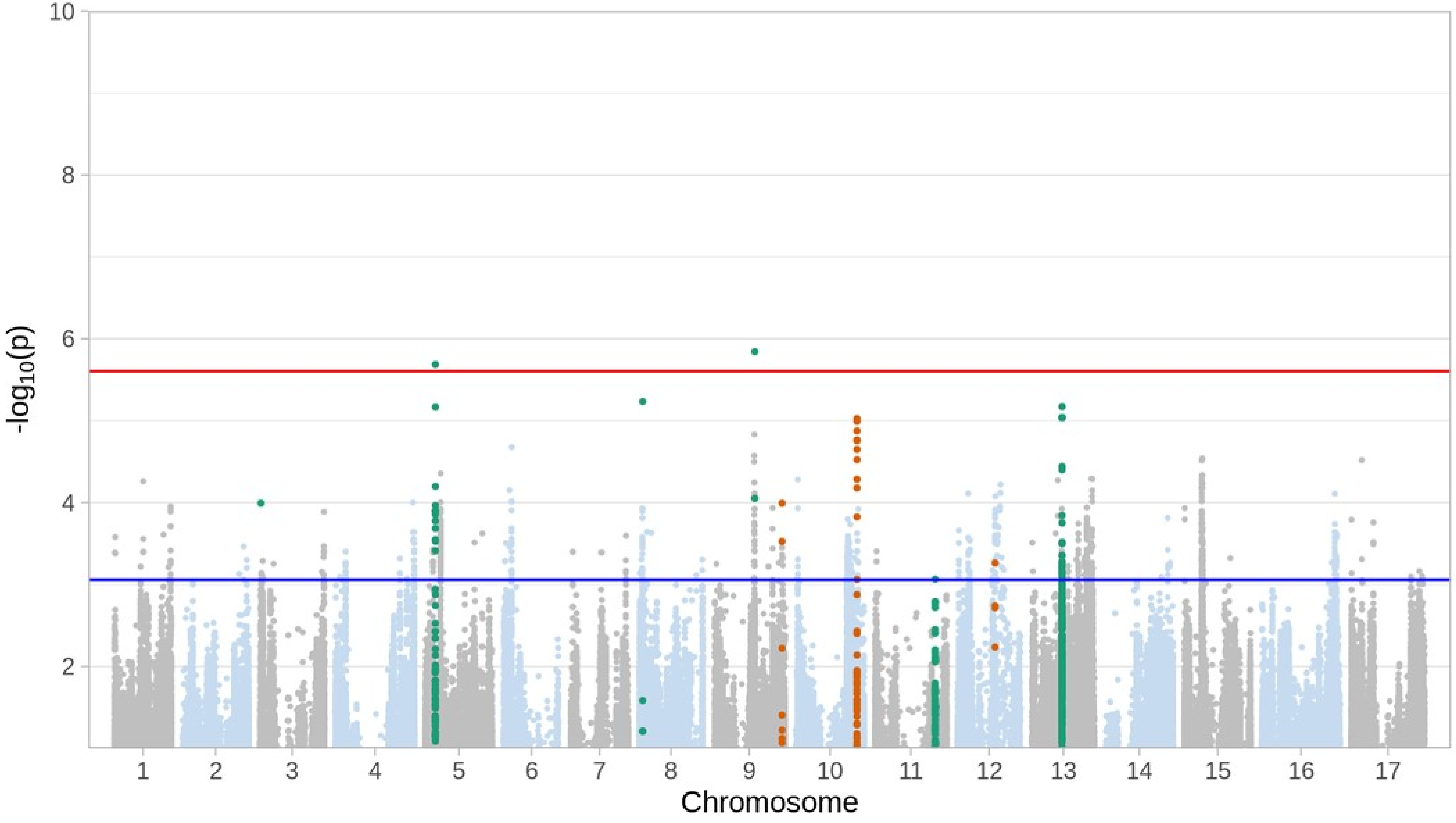

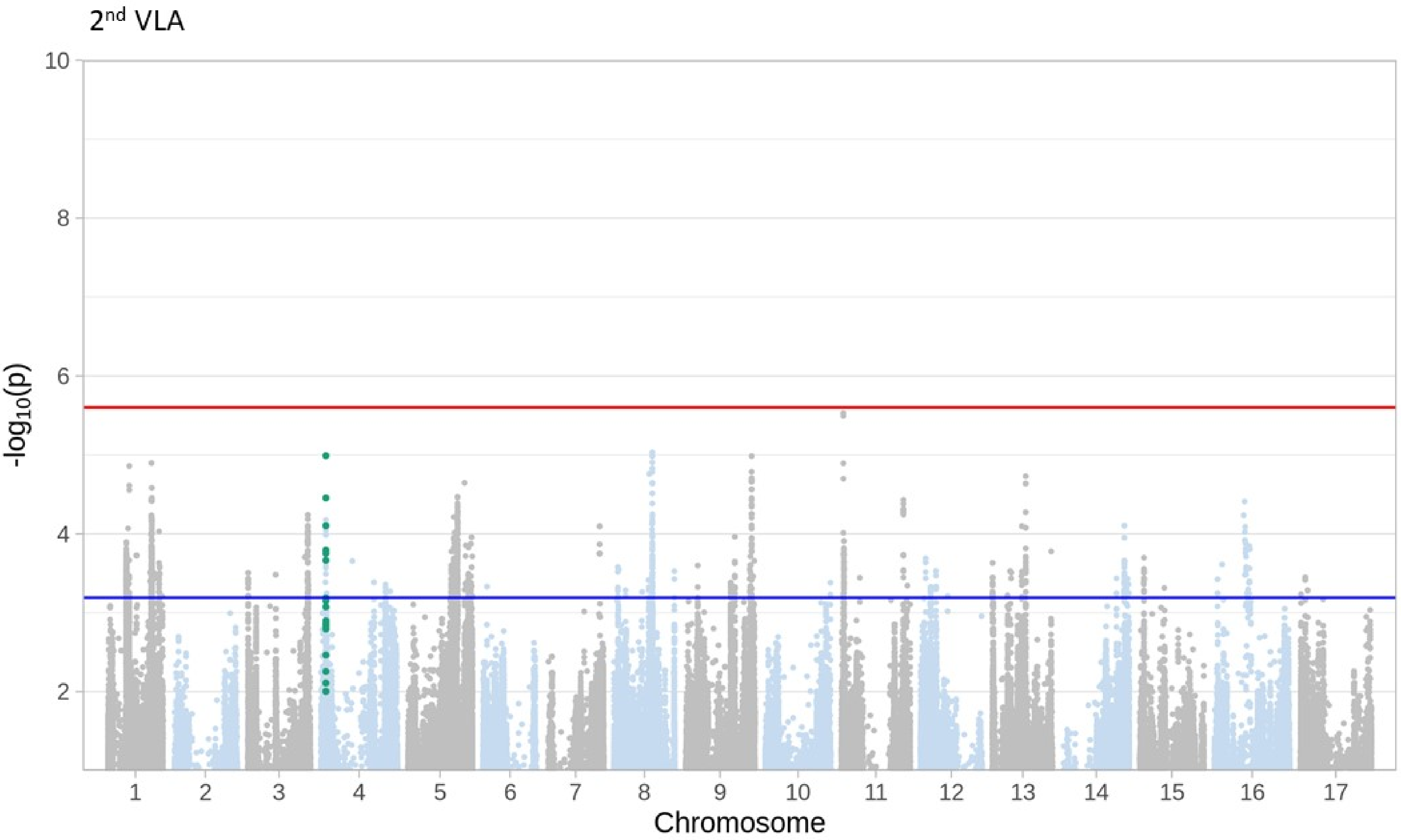

2nd VLA
10
8
6
4
2
-log10(p)
Chromosome

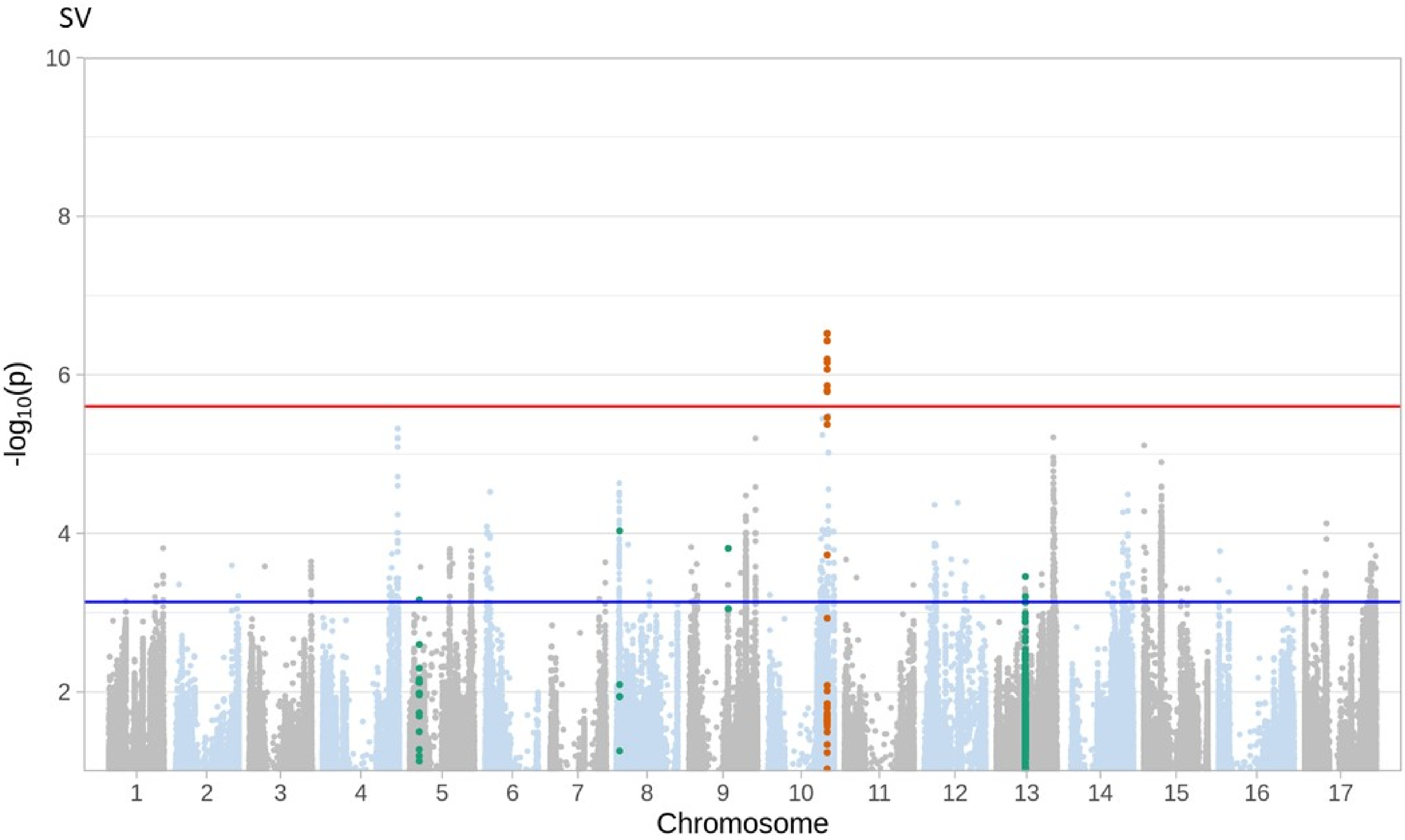
SV
10
8
6
4
2
-log10(p)
Chromosome

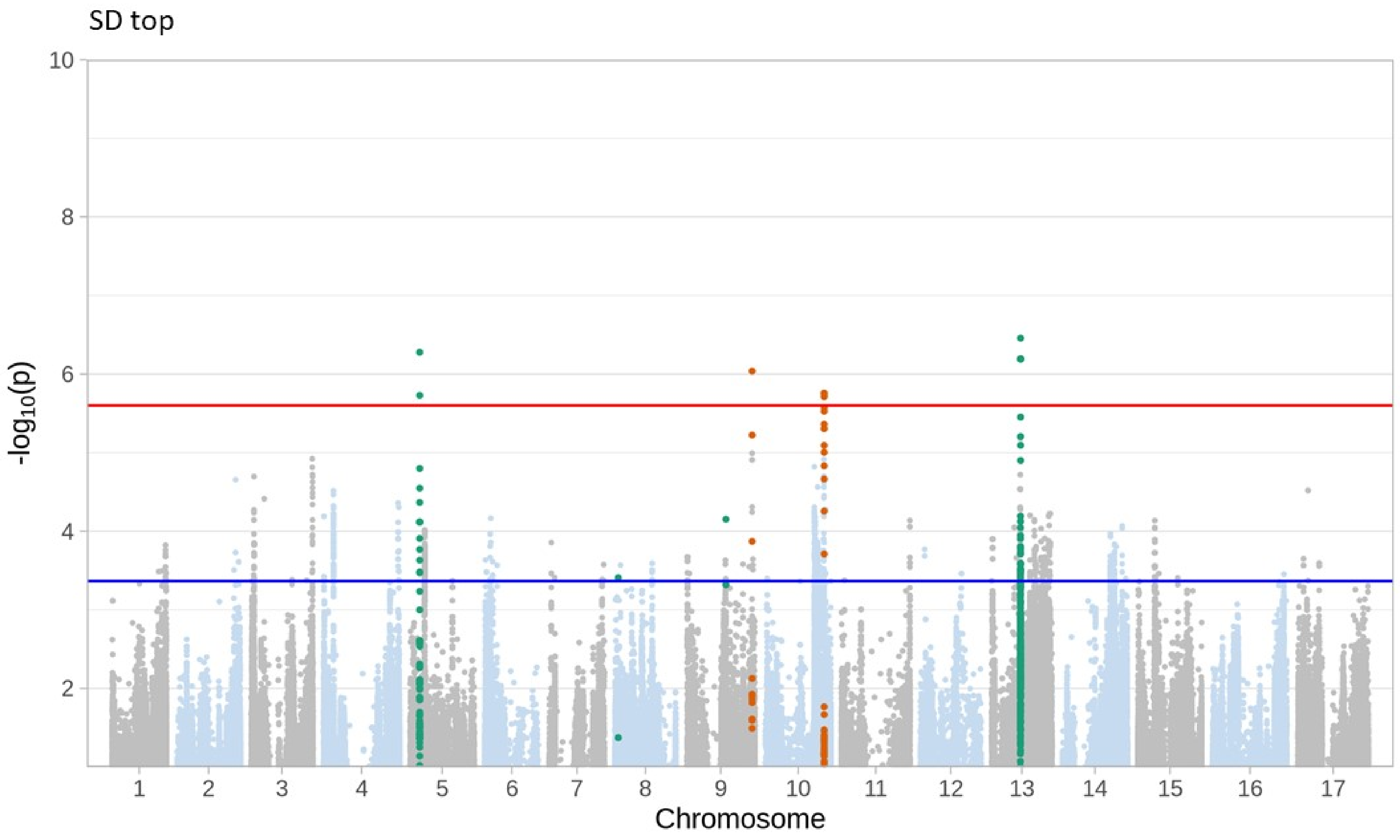
SD top
10
8
6
4
2
-log10(p)
Chromosome

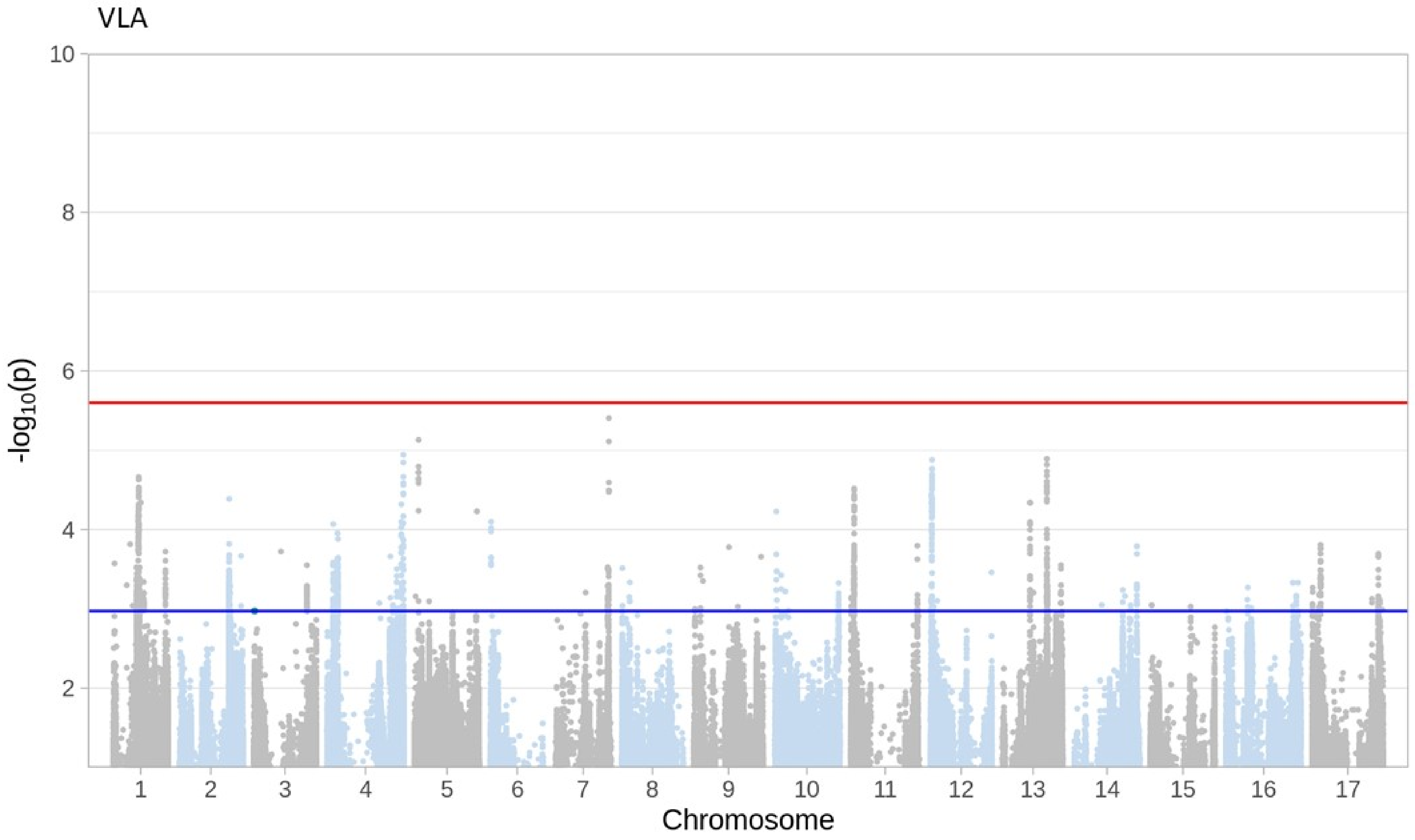
VLA
10
8
6
4
2
-log10(p)
Chromosome

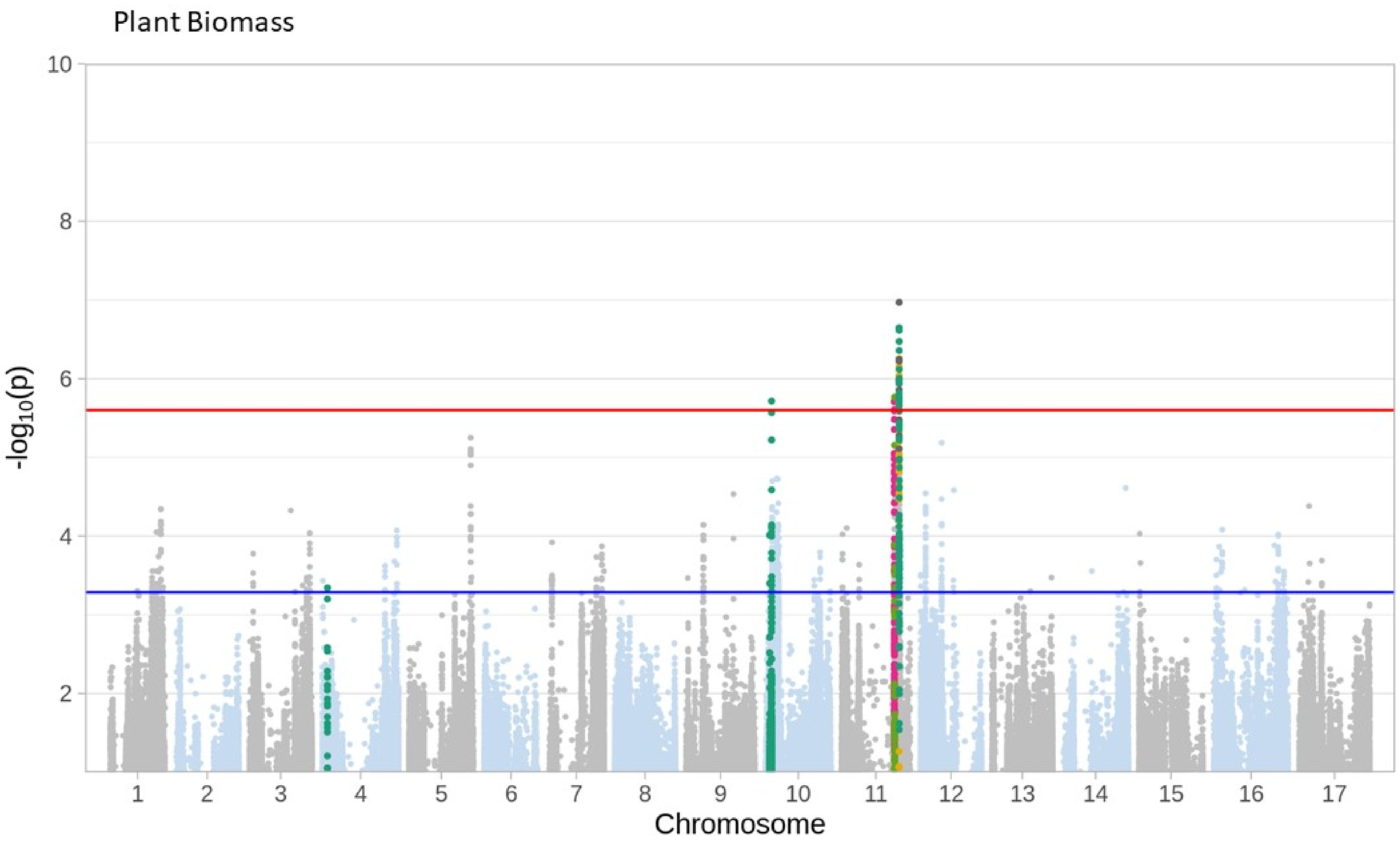
Plant Biomass
-log10(p)
10
8
6
4
2
Chromosome

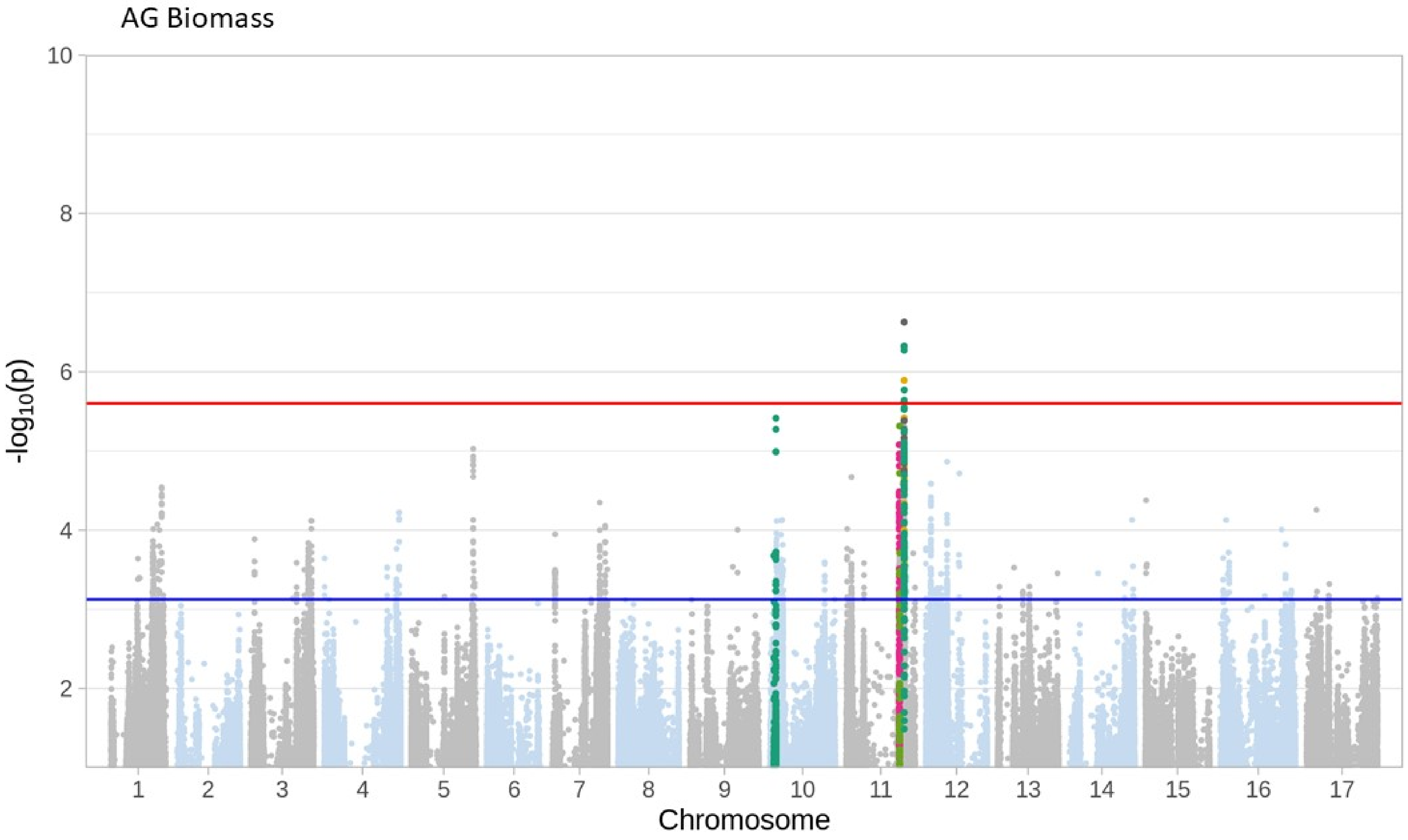
AG Biomass
10
8
6
4
2
-log10(p)
Chromosome

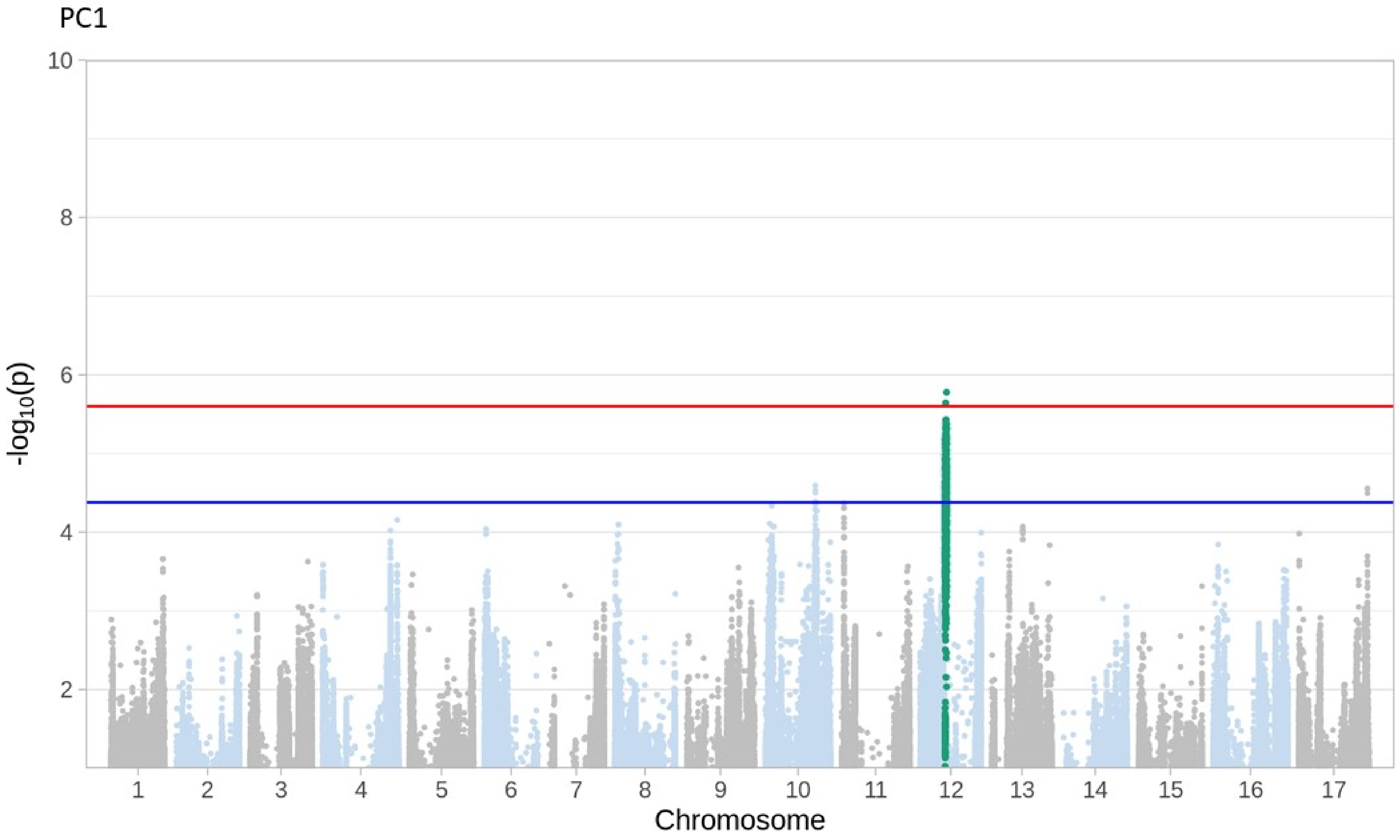
PC1
10
8
6
4
2
-log10(p)
Chromosome

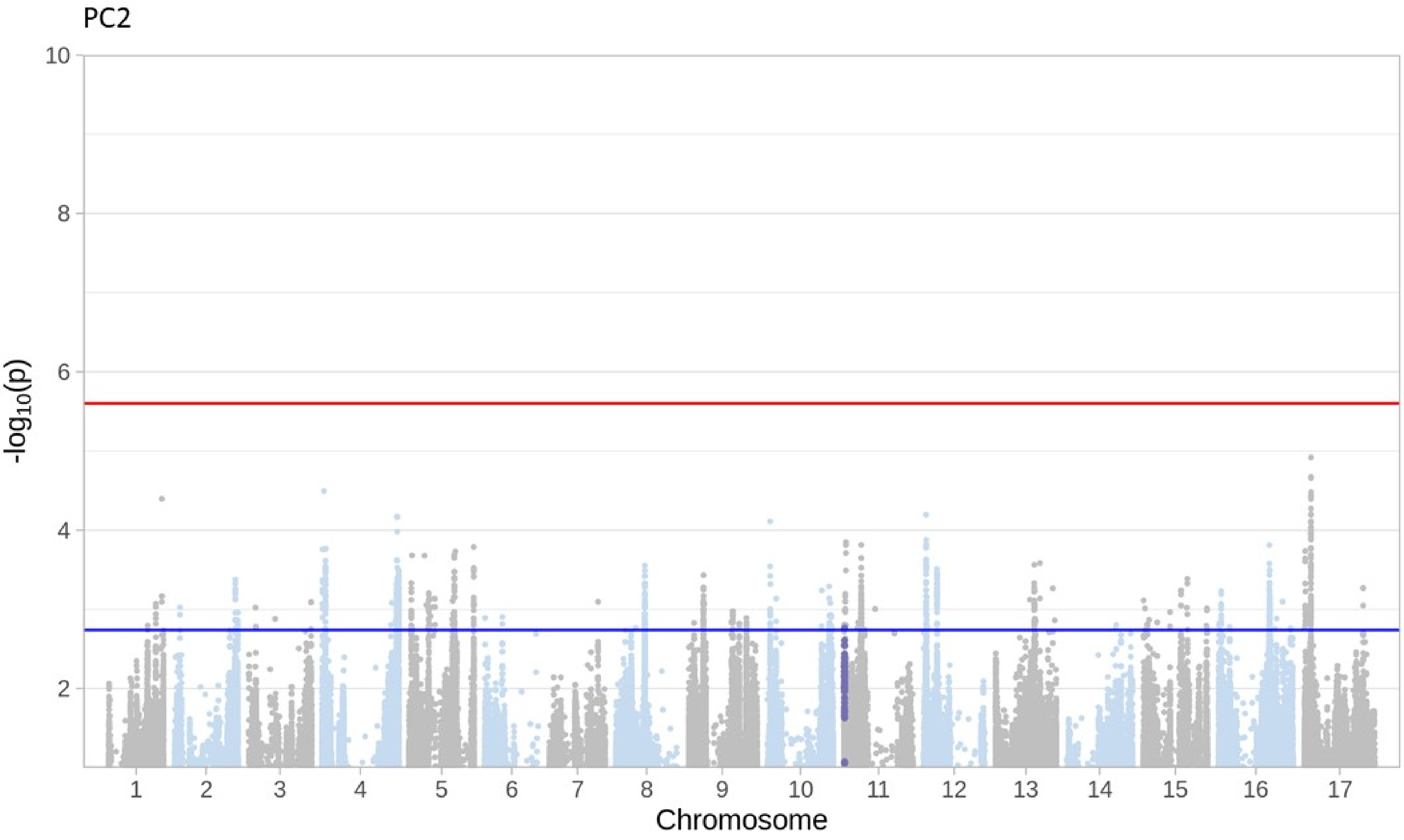
PC2
10
8
6
4
2
-log10(p)
1 2 3 4 5 6 7 8 9 10 11 12 13 14 15 16 17
Chromosome

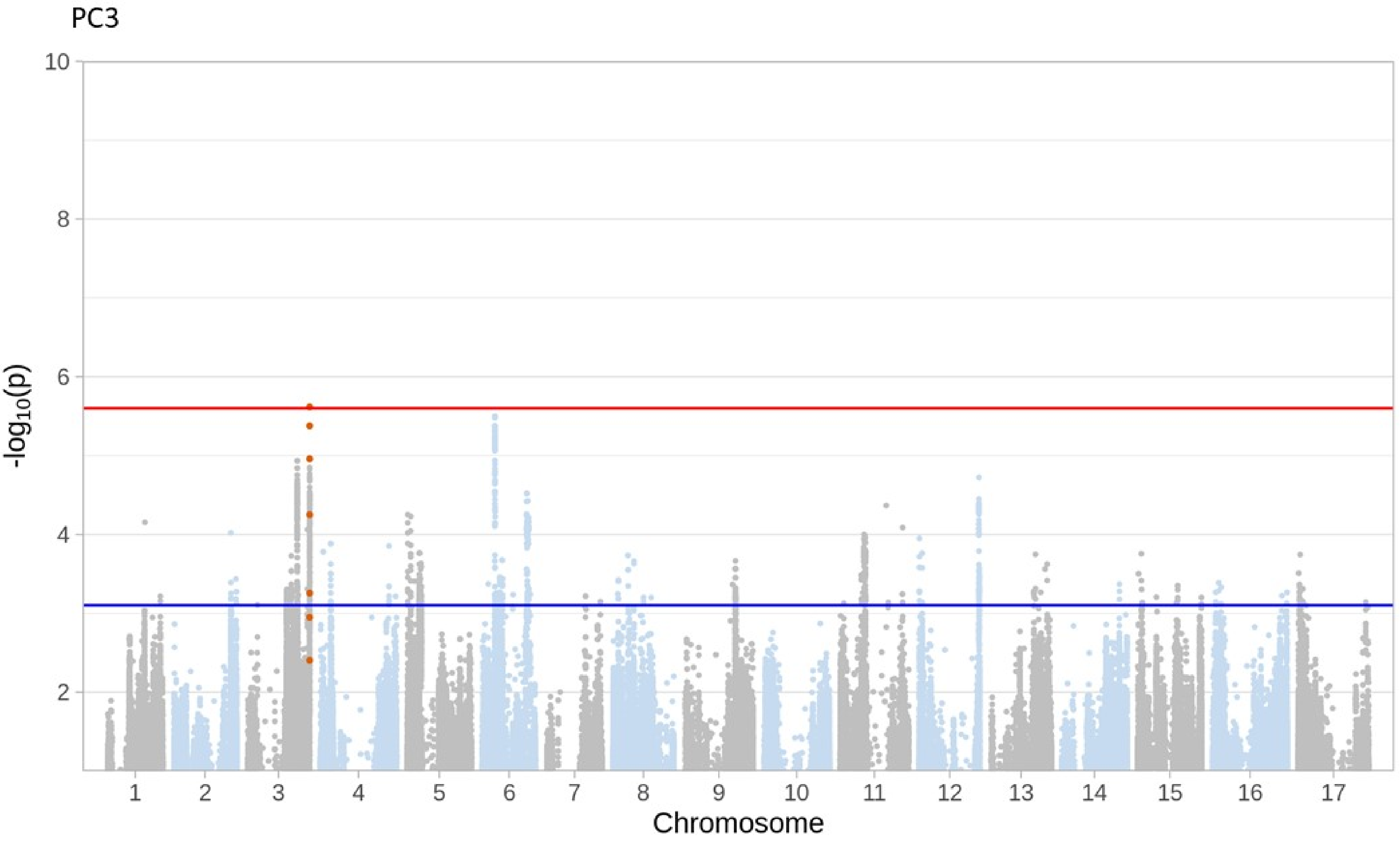

PC3
10
8
6
4
2
-log10(p)
Chromosome

**Figure S9**: *Distribution of the number of genes per significant region. Points represent the number of genes in each significant block. Note the log scale on the y-axis.*

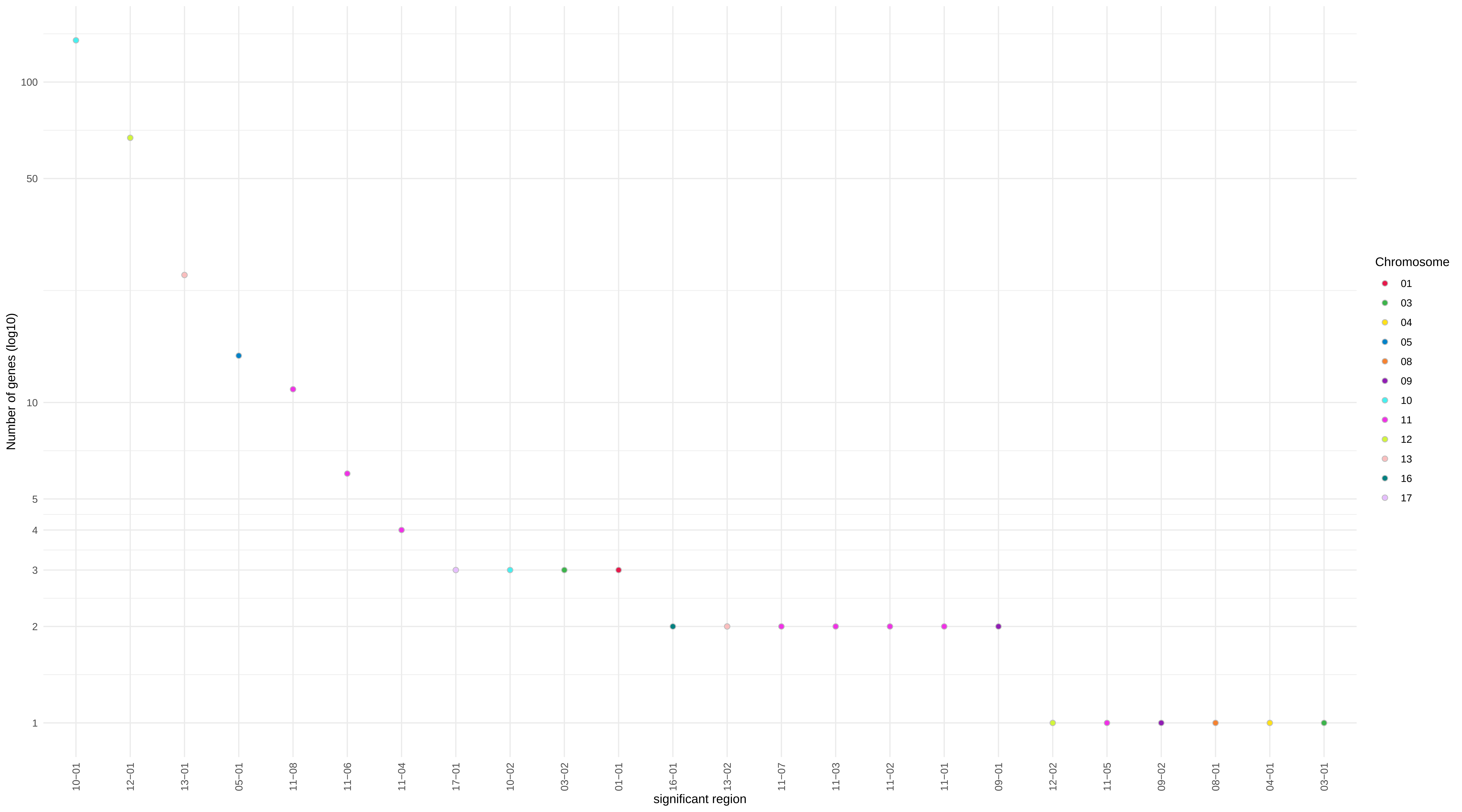

Number of genes (log10)
100
50
10
5
4
3
2
1
10−01
12−01
13−01
05−01
11−08
11−06
11−04
17−01
10−02
03−02
01−01
16−01
13−02
11−07
11−03
11−02
11−01
09−01
12−02
11−05
09−02
08−01
04−01
03−01
significant region
Chromosome
01
03
04
05
08
09
10
11
12
13
16
17

**Figure S10**: Visualization of *haplotypic blocks across the sunflower genome. Blocks are shown for each of the 17 sunflower chromosomes. Blocks are colored in an alternating fashion and colors are arbitrary. Black dots on the x-axis indicate singleton SNPs that did not fall within blocks.*

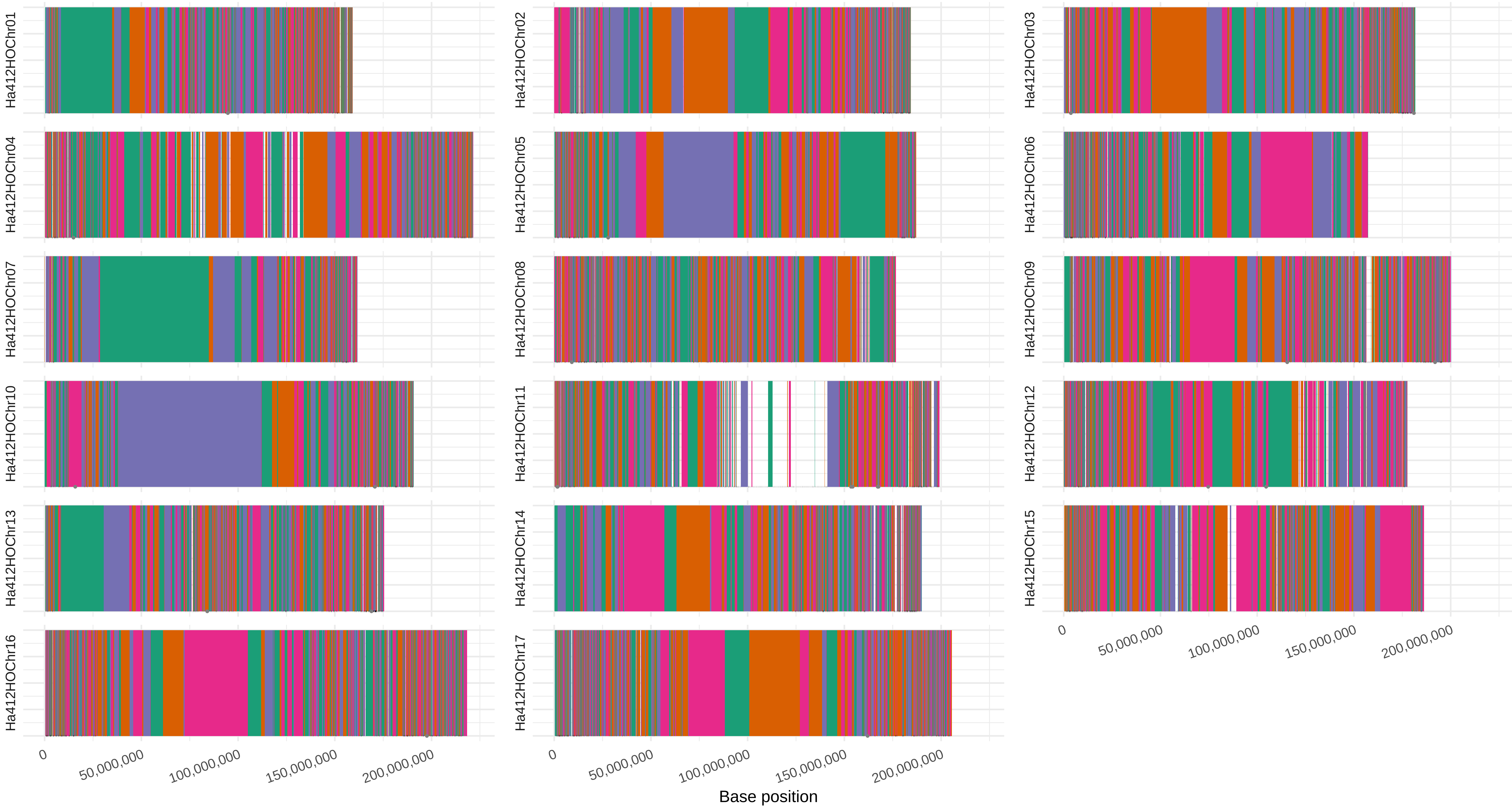
Ha412HOChr01
Ha412HOChr02
Ha412HOChr03
Ha412HOChr04
Ha412HOChr05
Ha412HOChr06
Ha412HOChr07
Ha412HOChr08
Ha412HOChr09
Ha412HOChr10
Ha412HOChr11
Ha412HOChr12
Ha412HOChr13
Ha412HOChr14
Ha412HOChr15
Ha412HOChr16
Ha412HOChr17
0
50,000,000
100,000,000
150,000,000
200,000,000
Base position

**Methods S1:** *Neural Network Architecture, Training and Prediction.*

The neural network followed that of the U-Net (Ronneberger *et al.*, 2015) deep neural network. High-dilation convolutions (Yu and Koltun, 2015) were integrated by replacing the U-Net architecture after the last max-pool through the first up convolution with the structure shown in Figure S1. In addition, all RELU activations were replaced with ELUs (Clevert *et al.*, 2015). The network architecture takes input images of size 572x572x3 pixels and outputs a segmented target image of 388x388x1 pixels that corresponds to the center of the 572x572 image. For training, input images were created by randomly selecting 388x388x3 regions from the leaf images along with the surrounding 92 pixels to create the 572x572x3 input image. As in Ronneberger et al. (2015), when surrounding data was missing at the edge of the image, the pixels were extrapolated by mirroring. The 572x572x3 bright-field leaf input images were independently normalized to have a mean of 0 and a standard deviation of 1. The network was coded using the pytorch (Paszke *et al.*, 2017) framework and trained using backpropagation using the Adam optimizer (Kingma and Ba, 2014). The loss function was weighted binary cross entropy. Non-vein pixels were given a weight of 0.05 and in-vein pixels a weight of 1. The weighting of the non-vein pixels was chosen using 3-fold cross validation of the training set of images using a grid search of 0.01, 0.05, 0.1, and 0.2 for the weight. Cross-validation training was performed out to 3000 batches, but there was no further decrease in the loss on the validation sets past 1900 batches (Figure S3).

Attempts at stitching 388x388 segmented tiles together to recreate the full size bright-field images resulted in incorrect segmentations at the boundaries between tiles. To avoid the boundary effects, the full-sized images (2584x1936 pixels) were mirror padded to 2876 x 2300 and passed through the network as one input. The images were normalized as described for the image tiles used in training. The resulting segmented images were cropped back to 2584x1936 pixels to remove padding pixels. The padding was used to both avoid edge effects – as done during network training – and to ensure all the convolution operations within the network were valid. The network outputs the probability a pixel is within a vein. The in-vein segmentation cutoff of 0.2 maximized the correlation between the number of pixels in the hand drawn vein lines and in that extracted with the network in a 3-way cross validation grid search of values spanning 0.2 to 0.9 with 0.1 increments.
